# *In vivo* base editing via single myotrophic adeno-associated viruses in dystrophic mouse muscle and satellite cells

**DOI:** 10.64898/2026.05.09.721064

**Authors:** Kuan-Hung Lin, Amy Lam, Samuel van Ooijen, Madeline Maier, Gracia Kassis, Regan Ellis, Kathleen Messemer, Jennifer Martin, Ramzi Khairallah, Amy Wagers

## Abstract

Duchenne muscular dystrophy (DMD) is the most common, lethal X-linked neuromuscular disorder of childhood and is caused by mutations in the *Dmd* gene that disrupt dystrophin expression. Although adeno-associated virus-mediated gene therapies hold tremendous promise for DMD treatment, their clinical applications have been limited by dose-dependent vector and genome-level toxicities. Here, we developed and tested a single-vector adenine base editing strategy as a potentially safer genome editing approach to recode the pathogenic nonsense mutation into a benign missense mutation in *mdx^4cv^*DMD mouse model. Delivered using a muscle-tropic adeno-associated virus (MyoAAV) at a clinically-feasible dose (4E13 VG/kg), this strategy enabled detectable molecular recoding of the *mdx^4cv^* mutation in mice ranging in age from 3 days to 6 months. Yet, the overall efficiency and therapeutic impact of *in vivo* base editing with this system was highest in mice treated at the juvenile stage, with animals administered MyoAAV vectors at 3 weeks of age showing robust recovery of dystrophin expression and significant improvement in muscle contractile properties only one month later. Notably, introduction of adenine base editors either earlier in development, in neonatal mice, or later, in adulthood, yielded substantially lower editing efficiencies, particularly in muscle satellite cells whose editing is essential to ensure durable rescue of dystrophin expression in growing and regenerating muscle. Taken together, these results demonstrate the therapeutic potential of single-vector adenine base editing for DMD and underscore the importance of recipient age and disease stage in achieving optimal treatment outcomes for this and other genetic muscle disorders.

## Introduction

Duchenne muscular dystrophy (DMD) is an inherited progressive muscle degenerative disease caused by deletions, point mutations, duplications, or inversions in the *Dmd* gene, which encodes the dystrophin protein. DMD mutations cause genetic frame-shifts or premature truncation of the dystrophin reading frame, leading to loss of protein expression^1^. At the cellular level, loss of dystrophin in myofibers disrupts the dystrophin-associated protein complex (DAPC), compromising sarcolemma stability, disrupting cellular signaling, and increasing vulnerability to muscle contraction-induced injury^1^. Restoring dystrophin in myofibers effectively rescues muscle function in several animal models^2^. Yet, because myofibers are post-mitotic and rely on postnatal muscle stem cells (satellite cells) for regular replacement and turnover^3^, whether restoration of dystrophin in myofibers alone will provide enduring therapeutic benefit remains unclear. Inefficient rescue of *Dmd* in satellite cells could, over time, lead to the accretion and replacement of previously DMD-restored myonuclei with new myonuclei derived from unmodified satellite cells. As healthy, rescued myofibers become increasingly replaced by fibers bearing a predominance of pathogenic myonuclei, the benefits of dystrophin protein restoration could wane.

Further emphasizing the importance of DMD gene rescue in satellite cells, recent evidence has revealed a critical cell-autonomous role for dystrophin in these cells^3^. Satellite cells lacking intact *Dmd* exhibit higher apoptotic and senescence rates and reduced differentiation capacity^4^, ultimately leading to inadequate muscle regeneration^1,5^. Thus, effective long-term rescue of muscle function in DMD patients requires targeting of both myofibers and satellite cells to ensure complete and long-lasting functional restoration.

Adeno-associated viruses (AAVs) have frequently been used to deliver gene therapy cargoes because of their high transduction rates, gene transfer without host genome integration, relatively low toxicity, and adjustable tissue tropism by use of different serotypes^6^. In pre-clinical studies focused on Duchenne muscular dystrophy, multiple AAV-based gene therapy strategies have been documented, in some cases achieving restoration of dystrophin expression in over 90% of myofibers. Tested strategies include delivery of full-length dystrophin^7^, CRISPR-Cas9-based exon excision^8^, homology-directed repair^9^, base editing^10,11^, and prime editing^11^. Compared to ectopic expression of dystrophin proteins, genome editing-based approaches offer the advantage of introducing a permanent genomic correction into the mutant *Dmd* locus. Additionally, unlike CRISPR-Cas9-based approaches, base editing^12^ and prime editing^13^ do not cause DNA double-strand breaks and thus present a lower risk of genotoxicity^14^. However, the limited packaging capacity of AAV,^15^ which is smaller than that required for first-generation base editing- and prime editing-based strategies, has thus far necessitated use of an intein-split dual AAV delivery approach that divides the encoded base editor or prime editor into amino-terminal (N-terminal) and carboxy-terminal (C-terminal) parts that can only be reassembled within cells transduced by both vectors^16^. For these reasons, the design and manufacturing of AAV vectors for base and prime editing have been complicated and usually require an initial screen to identify the optimal intein sequences to be used and the best location for genome editor splitting. Furthermore, the need for co-transduction of heterotypic AAVs frequently necessitates an undesirable high AAV dose for these strategies.

Recent advancements in size-minimized adenine base editor (ABE) designs, including those that can fit within a single AAV vector, provide a potential solution to this packaging challenge. ABE is derived from the bacterial enzyme tRNA-specific adenosine deaminase A (tadA), which converts adenosine to inosine in tRNA molecules^17^. Because inosine is recognized as guanine by DNA polymerase, adenines converted to inosine eventually are replaced by guanine through DNA repair and replication, thereby achieving an A to G conversion. Gaudelli et al. took advantage of this biochemical activity of tadA to engineer a programmable ABE, evolving and fusing tadA with a catalytically impaired Streptococcus pyogenes Cas9 mutant (dSpCas9)^12^. When a SpCas9 Protospacer Adjacent Motif (PAM) compatible gRNA is provided, the fused dSpCas9 domain allows ABE to be guided to the gRNA target site and instills A.T → G.C conversions (depending on whether the provided gRNA targets the sense or antisense strand of DNA). This A.T → G·C capacity offers the potential to correct half of all known pathogenic single-nucleotide mutations^12^.

Multiple gene therapies utilizing base editors to treat inherited and also chronic recurring infectious diseases that affect various tissues have been designed and are currently entering clinical trials^18^. With the phage-assisted continuous evolution approach, Richter et al. further evolved the original TadA7.10 to TadA8e, achieving increased editing efficiency and broadened Cas domain compatibility^19^. This breakthrough made it possible to engineer smaller ABEs by fusing TadA8e with a smaller dCas9 domain derived from Cas9 found in other bacterial species, such as SaCas9, SauriCas9, CjCas9, and Nme2Cas9^20^. These size-minimized ABEs for the first time enabled the packaging of an ABE with its compatible gRNA in a single AAV vector. Use of such a single-vector approach for targeting of rare muscle satellite cells in vivo should support higher rates of gene correction than achieved with previously reported gene therapies that employed dual, or even triple, AAV delivery approaches.

In this study, we designed a single-vector MyoAAV-based and ABE-mediated gene therapy approach, in which we utilized the size-minimized SaABE8e (TadA8e fused with nuclease-deactivated SaCas9) to treat the *mdx^4cv^* DMD disease mouse model. We chose to use MyoAAV, a muscle-tropic AAV serotype, due to its greater rate of transduction of cardiomyocytes, myofibers, and satellite cells and its lower transduction of liver hepatocytes, compared to conventional AAVs^21^. The liver-detargeting property of MyoAAV^21^ also reduces hepatotoxicity – the most common adverse event observed in AAV gene therapy^22^.

The *mdx^4cv^* mouse line carries a C→T point mutation in exon 53, which causes a CAA→TAA nonsense mutation. By systemically delivering SaABE8e-required gene correction components via MyoAAV at a clinically feasible dose (4E13 viral genome/kg) to *mdx^4cv^* mice, the TAA nonsense mutation was successfully converted to a TGG missense mutation in myofibers, cardiomyocytes, and satellite cells. Treated mice also exhibited rescued dystrophin expression in myofibers and cardiomyocytes. *In silico* protein structure prediction showed that the rod-like structure formed by the wild-type amino acid sequence was minimally affected by the introduced missense mutation, strongly suggesting that dystrophin bearing the therapy-generated missense mutation remained functional. Consistent with this prediction, muscle physiology assessments demonstrated significant improvements in the force output of the gastrocnemius muscles of treated mice. Finally, to determine the optimal developmental window for dystrophin rescue in *Dmd*-edited mice, we evaluated differences in base editing efficiency among mice treated at neonatal, juvenile, or adult stages, and observed the highest editing efficiency in juvenile (P21-28) mice.

Altogether, these data provide proof-of-principle demonstration that a systemically administered, single AAV vector-based, ABE-mediated genomic medicine strategy can effectively recode the *Dmd* nonsense mutation in endogenous myofibers, cardiomyocytes, and satellite cells, rescuing functional dystrophin expression and muscle contractile activity. They further reveal the importance of recipient age and stage of disease progression for achieving optimal editing efficiency and maximal therapeutic benefit. Results from these pre-clinical studies should aid the successful development and deployment of clinical base editing strategies for DMD and other, currently incurable, genetic diseases of muscle.

## Results

### A single-AAV vector for SaABE8e-mediated genome editing of the *mdx^4cv^* nonsense mutation

Treatment of DMD with a single vector AAV-based *in vivo* gene editing approach provides numerous potential benefits, including reducing the vector dose required for gene therapy and simplifying AAV manufacturing. We chose to test such an editing approach in the DMD model *mdx^4cv^*, which carries a nonsense mutation in exon 53 of the *Dmd* gene (CAA>TAA), as these mice exhibit near-complete loss of Dystrophin protein^23^. To conform to the packaging limits of a single-AAV vector (<5.2 kb ^15^), we designed our strategy around a recently engineered size-minimized adenine base editor (SaABE8e)^20^. As shown in **Fig. 1A**, we identified two SaABE8e gRNAs (4cv-gRNA1 and 4cv-gRNA2) with the potential to convert the *mdx^4cv^* nonsense mutation (TAA) to a tryptophan coding sequence (TGG), thereby rescuing the protein reading frame.

**Fig. 1.**
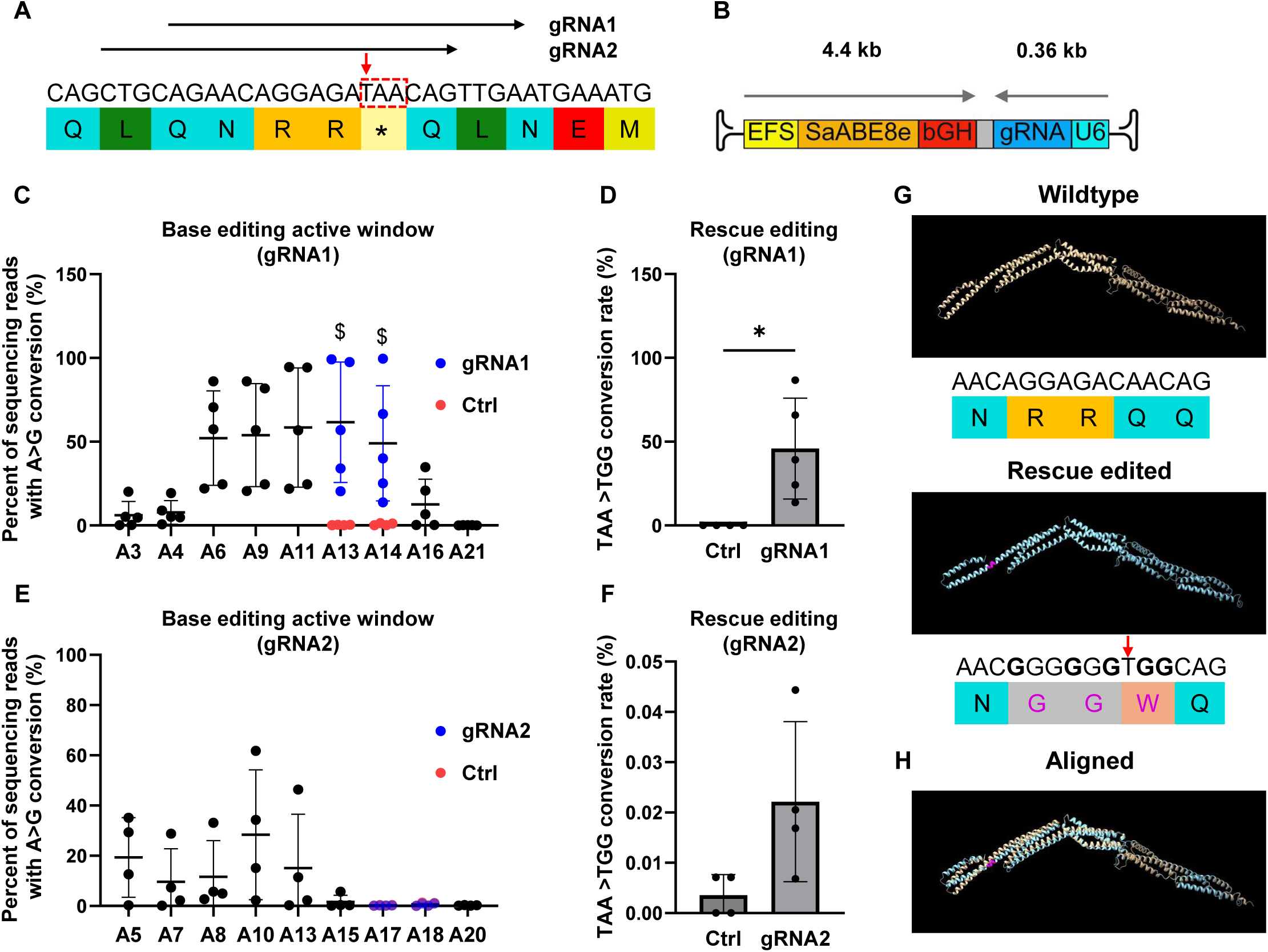
SaABE8e-mediated genome editing bypasses *mdx^4cv^* nonsense mutation in FACS-isolated satellite cells *in vitro*. (A) Genomic sequence, encoded amino acids and gRNA recognition sequences in the *mdx^4cv^*allele (red box). Red arrow highlights the C>T mutation. (B) Schematic of AAV-SaABE8e^4cv^ vector. Arrows indicate transcript orientation of EFS SaABE8e and U6 sgRNA cassettes. (C-F) FACS-isolated satellite cells from juvenile *mdx^4cv^* male mice (Fig. S1) were transfected with AAV-SaABE8e plasmids carrying 4cv-gRNA1 (C-D) or 4cv-gRNA2 (E-F), or with EGFP encoding plasmid to indicate transfection efficiency (Fig. S2). A>G conversion rates were calculated from sequencing reads and normalized to transfection efficiency for each experiment. (C,E) Normalized A>G conversion rates calculated for individual adenines (A) within each gRNA-targeting site. Adenine conversions in the *mdx^4cv^* premature stop codon are shown in dark blue (A13A14 for gRNA1 and A17A18 for gRNA2). Normalized A>G conversion rates for these adenines from control cells (ctrl) transfected with EGFP plasmid only are plotted in red. $ indicates a significant difference in A>G conversion rates (determined by unpaired T test) between 4cv-gRNA1 or 4cv-gRNA2 and control-transfected satellite cells. (D,F) Normalized conversion rates for reads in which the *mdx^4cv^* nonsense mutation is converted to a non-stop (missense) mutation (TAA>TGG, defined as rescue editing). n = 4-5 biological replicates/group; D and F analyzed with unpaired T-test. Data are shown as mean ± SD. * indicates P < 0.05. (G) *In silico* predicted central rod domain structure of wildtype dystrophin protein (top) and the most abundant rescue variant detected in SaABE8e-gRNA1 edited *mdx^4cv^* satellite cells (bottom). Genomic sequence and encoded amino acids are shown beneath each structure. In the rescue-edited variant, guanines converted from adenine are shown in bold, with resulting changes to the amino acid sequence highlighted in magenta. The red arrow indicates the C>T mutation site. (H) Alignment of *in silico* predicted wild-type and rescue-edited structures shows near-perfect overlap.

### SaABE8e-mediated genome editing bypasses the *mdx^4cv^* nonsense mutation in FACS-isolated satellite cells *in vitro*

To test the efficiency of 4cv-gRNA1 and 4cv-gRNA2 for recoding the *mdx^4cv^* mutation, we constructed SaABE8e expression constructs carrying each gRNA (SaABE8e^4cv-gRNA1^ or SaABE8e^4cv-gRNA2^) (**Fig. 1B**), using a recombinant AAV cargo plasmid as backbone. We then transfected each plasmid into FACSorted primary mouse satellite cells (Sca-1^-^CD11b^-^Ter119^-^CD45^-^CD29^+^CXCR4^+^), isolated as previously described^24^ from juvenile (P21) male *mdx^4cv^* mice (**Fig. S1**). To estimate transfection efficiency for each batch of experiments, an extra well of cells was transfected with an EGFP reporter plasmid. Four days after transfection, genomic DNA was harvested from cultured cells for amplicon sequencing to determine editing rates at the *4cv* mutation site (**Fig. 1C-F**), with the frequency of sequenced reads bearing SaABE8e-mediated A>G conversions at each adenine at the gRNA targeting site defined as “conversion rate”. To correct for batch-to-batch differences in transfection efficiency, conversion rates were further normalized to the rate of EGFP positivity (%EGFP+, determined by flow cytometry of EGFP plasmid-transfected cells, **Fig. S2**) for each experiment.

For both gRNAs, SaABE8e exhibited high activity for editing adenines between the 6^th^ and 14^th^ position at the gRNA targeting sites (representative allele frequency tables are shown in **Fig. S3**), consistent with previously reported data^19^ (**Fig. 1C** and **1E**). The A>G conversion rate at the most efficiently edited adenine (A13) in SaABE8e^4cv-gRNA1^ transfected satellite cells was 61.7 ± 35.9% (**Fig. 1C**), whereas SaABE8e^4cv-gRNA2^ showed a conversion rate of only 28.4 ± 25.9% at the most efficiently edited adenine (A10, **Fig. 1E**). Additionally, in SaABE8e^4cv-gRNA1^ transfected satellite cells, 45.9 ± 30.1% of reads exhibited conversion of both adenines in the TAA nonsense mutation, bypassing the premature stop and converting this codon to TGG, encoding tryptophan (rescue editing, **Fig. 1D**). In comparison, only 0.02% of reads in SaABE8e^4cv-gRNA2^ transfected satellite cells showed this double conversion rescue edit (**Fig. 1F**). This difference is likely attributable to the lower overall editing efficiency of SaABE8e within the 4cv-gRNA2 targeting window and the location of the 4cv-gRNA2 PAM site, which places the adenines targeted for rescue editing (A17 and A18) outside of the optimal SaABE8e active window (**Fig. 1E**).

Although 4cv-gRNA1 editing rescued the *mdx^4cv^* nonsense mutation, SaABE8e also converted other adenines to guanines within the active window (**Fig. S3**), causing an additional change of up to 5 amino acids at the target site (wild-type: QNRRQQ > RGGGWR). To predict the likelihood that such missense mutations introduced by SaABE8e^4cv-gRNA1^ could impact the structure or function of the encoded protein, we performed *in silico* modeling of the dystrophin central rod domain structure formed by the wild-type amino acid sequence and the most abundant *mdx^4cv^* rescue variant in SaABE8e^4cv-gRNA1^-transfected satellite cells (**Fig. 1G**). As shown in **Fig. 1H**, the rescue variant shows very little difference in the predicted structure, suggesting that the modified dystrophin protein produced by SaABE8e^4cv-gRNA1^-mediated rescue editing is likely to be functional. Thus, we proceeded with 4cv-gRNA1 as the best candidate for further testing *in vivo*.

### Systemic delivery of MyoAAV-SaABE8e^4cv^ in juvenile *mdx^4cv^*mice rescues dystrophin deficiency and restores muscle function

Adeno-associated viruses (AAVs) are known for their adjustable tissue tropism with different serotypes^6^. A recent study^21^ identified a new family of muscle-targeted “MyoAAVs” that exhibit markedly enhanced efficiency for transducing skeletal muscle myofibers and satellite cells, as well as cardiomyocytes – all cell types of high relevance to DMD pathology. Notably, MyoAAVs also showed significantly reduced liver tropism, which minimizes the risk of hepatotoxicity commonly associated with conventional AAV vectors^22^. Thus, to test the in vivo efficacy of our base editing approach, we packaged SaABE8e**^4cv-gRNA1^** into MyoAAV2A (hereafter referred to as MyoAAV-SaABE8e^4cv^), an evolved MyoAAV subtype that transduces mouse skeletal muscle with particularly high efficiency. As a negative control, we constructed a single AAV SaABE8e plasmid carrying a non-targeting gRNA (SaABE8e^NT^) and packaged it into MyoAAV2A (MyoAAV-SaABE8e^NT^). Equal amounts (4E13 vector genomes/body mass (VG/kg)) of either MyoAAV-SaABE8e^4cv^ or MyoAAV-SaABE8e^NT^ were delivered systemically, by retro-orbital (RO) injection, into postnatal day 21 (P21) male *mdx^4cv^* mice.

Tissues harvested from injected mice one month later demonstrated efficient restoration of dystrophin protein expression in the tibialis anterior (TA), diaphragm, and heart of MyoAAV-SaABE8e^4cv^ injected mice (**Fig. 2A**). Quantification of immunostained sections indicated that 30.8 ± 6.4% of myofibers in TA, 16.5 ± 5.9% of myofibers in diaphragm, and 33.7 ± 6.2% of cardiomyocytes in hearts of MyoAAV-SaABE8e^4cv^ injected *mdx^4cv^* mice were dystrophin-positive (**Fig. 2B**). At the same time, dystrophin-positive myofibers and cardiomyocytes were essentially undetectable in the TA, diaphragm, and heart of control MyoAAV-SaABE8e^NT^ injected *mdx^4cv^* mice (**Fig. 2A, B**). Western blot analysis confirmed dystrophin rescue in MyoAAV-SaABE8e^4cv^ injected *mdx^4cv^* mice to levels 10.5 ± 8.0%, 8.1 ± 6.6 %, and 16.8 ± 7.1% of WT, in the TA, diaphragm, and heart, respectively (**Fig. S4 and 2C**). We also detected restoration of dystrophin-associated protein complex (DAPC) components at the myofiber surface in dystrophin-expressing cells of MyoAAV-SaABE8e^4cv^, but not MyoAAV-SaABE8e^NT^, injected *mdx^4cv^* mice (**Fig. S5**). These data confirm that the rescued dystrophin protein mediates functional interactions with these essential sarcolemmal proteins.

**Fig. 2.**
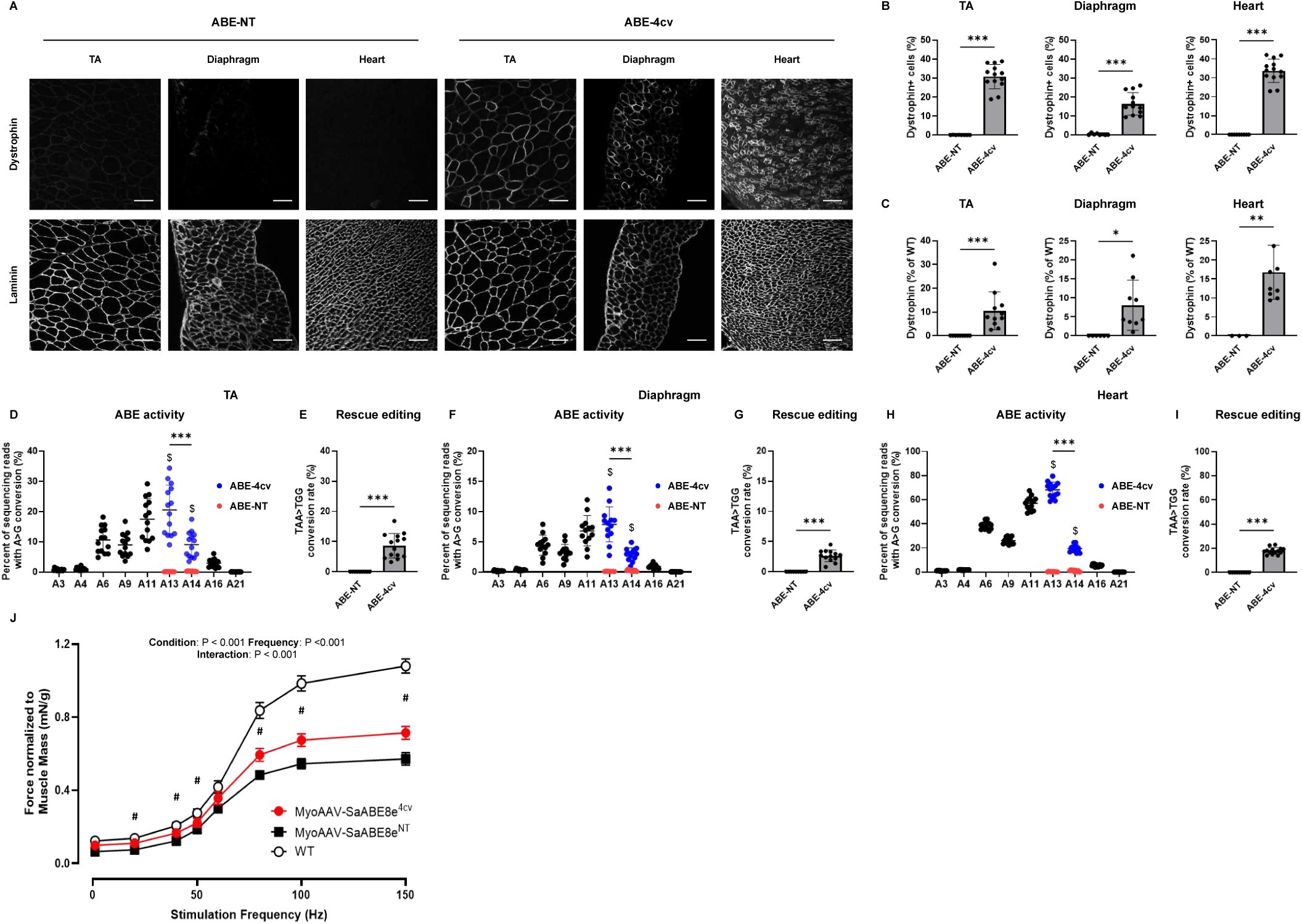
Systemic delivery of MyoAAV-SaABE8e^4cv^ restores dystrophin expression in *mdx^4cv^*mice. Juvenile (P21) male *mdx^4cv^* mice were injected retro-orbitally with 4E13 VG/kg MyoAAV-SaABE8e^4cv^ (ABE-4cv) or MyoAAV-SaABE8e^NT^ (ABE-NT), and tissues were harvested one month later. (A) Representative single-channel immunofluorescence images from ABE-NT or ABE-4cv injected mice showing the same field of tibialis anterior (TA), diaphragm, or heart stained for dystrophin (top) or laminin (bottom), as indicated. Scale bar = 100 µm. (B) Quantification of dystrophin-positive myofibers or cardiomyocytes, calculated as percent (%) of total cells (derived from laminin staining). (C) Semi-quantitative analysis of dystrophin protein expression in TA, diaphragm, and heart of mice injected with MyoAAV-SaABE8e^NT^ or MyoAAV-SaABE8e^4cv^. Representative blots are shown in Fig. S4. (D-I) cDNA from TA (D-E), diaphragm (F-G), or heart (H-I) were analyzed by amplicon sequencing at the *mdx^4cv^* mutation site. Representative allele frequency tables are shown in Fig. S6. (D,F,H) A>G conversion rates for individual adenines (A) of MyoAAV-SaABE8e^4cv^-injected *mdx^4cv^* mice were calculated from sequencing reads. Adenine conversions in the *mdx^4cv^* premature stop codon from MyoAAV-SaABE8e^4cv^-injected mice are shown in dark blue (A13 and A14). A13 and A14 A>G conversion rates from control, MyoAAV-SaABE8e^NT^-injected *mdx^4cv^* mice are shown in red. $ indicates a significant difference between MyoAAV-SaABE8e^4cv^- and MyoAAV-SaABE8e^NT^-injected *mdx^4cv^* mice in A>G conversion rates on the indicated adenine (determined by unpaired T-test). (E,G,I) Percent of reads in which the *mdx^4cv^*nonsense mutation is converted to a non-stop (missense) mutation (TAA>TGG). (J) Male wildtype (WT), MyoAAV-SaABE8e^4cv^-injected *mdx^4cv^* mice, and MyoAAV-SaABE8e^NT^-injected *mdx^4cv^* mice were analyzed for *in vivo* gastrocnemius muscle force output. Force output (mN) was normalized to muscle mass (g) and plotted against the stimulation frequency (Hz). # indicates significant difference (P < 0.05) between the MyoAAV-SaABE8e^4cv^- and MyoAAV-SaABE8e^NT^-injected *mdx^4cv^* mice. n = 8-14 biological replicates/group; B, C, E, G, and I are analyzed with unpaired T-test, D, F, and H are analyzed with one-way repeated measures ANOVA, J is analyzed with two-way repeated measures ANOVA. Data are shown as mean ± SD. *, **, and *** indicate P < 0.05, < 0.01, and <0.001, respectively.

To determine how efficiently *in vivo* base editing can recode the *mdx^4cv^* mutation at a molecular level, we sequenced genomic DNA and cDNA from TA, diaphragm, and heart of *mdx^4cv^* mice injected with MyoAAV-SaABE8e^4cv^ or MyoAAV-SaABE8e^NT^, focusing specifically on SaABE8e activity at the *mdx^4cv^* mutation site. Interestingly, while *in vitro* editing results suggested that editing rates at positions A13 and A14 were comparable for TAA→TGG conversion (**Fig. 1C**), data from *in vivo* edited TA, diaphragm and heart indicated that the A>G conversion rate for adenine 14 (A14) was significantly lower than the rate at A13 for both cDNA (**Fig. 2D, F, H**) and genomic DNA (**Fig. S7**), making the A14 A>G conversion rate limiting for rescue editing of the *mdx^4cv^* nonsense mutation in tissue. Overall, ABE activity *in vivo* appeared to vary significantly across tissues. Comparing A>G conversion at A13 (the most efficiently converted adenine), SaABE8e editing rates detected at the cDNA level were 20.6 ± 8.2% in the TA, 7.9 ± 2.9% in the diaphragm, and 68.2 ± 6.4% in the heart (**Fig. 2D, F, H**). Due to the significantly lower A14 A>G conversion rate in tissues in vivo, rescue editing (TAA→TGG) was detected at substantially lower efficiency in these same tissues (8.6 ± 4.1% (TA), 2.6 ± 1.0% (Diaphragm), and 17.8 ± 2.7% (Heart); **Fig. 2E, J, I**).

We also performed *in silico* analysis to predict possible off-target edits for 4cv-gRNA1 and found that none of these predicted off-targets are located in coding regions of the mouse genome (**Fig. S8A**). Amplicon sequencing of genomic DNA from the TAs and hearts of *mdx^4cv^* mice administered MyoAAV-SaABE8e^4cv^ at the two loci with the highest predicted off-target scores detected only minimal SaABE8e activity on Chr15: 48935095 in heart (0.7 ± 0.06% A>G conversion rate at A9 (with the highest conversion rates) at this locus in MyoAAV-SaABE8e^4cv^ injected mice compared to 0.12 ± 0.04% in MyoAAV-SaABE8e^NT^ injected mice) (**Fig. S8B-C**). In contrast, conversion rates at A13 were much higher (11.8 ± 2.0%) at the on-target site (**Fig. S7**).

Finally, to assess the impact of gene editing on muscle function in *mdx^4cv^* mice, which have a relatively severe muscle function defect due to a low frequency of revertant fibers ^25^, we subjected male *mdx^4cv^* mice to *in situ* measurement of muscle contractile force four weeks after administration of MyoAAV-SaABE8e^4cv^ or MyoAAV-SaABE8e^NT^ (as a control). Gastrocnemius muscles demonstrated significantly improved force output (normalized to muscle weight) in MyoAAV-SaABE8e^4cv^ injected mice, compared to MyoAAV-SaABE8e^NT^ injected mice (**Fig. 2J**); however, susceptibility to repetitive eccentric contraction-induced injury was not different from controls (**Fig. S9**). This result may indicate that a higher frequency of dystrophin-expressing muscle fibers, or a longer duration of dystrophin restoration, is needed to protect from mechanical damage than is required to boost muscle force production.

### Systemic delivery of MyoAAV-SaABE8e^4cv^ in juvenile *mdx^4cv^* mice sustains production of edited Dystrophin protein throughout adulthood in the heart

To determine the durability of MyoAAV-SaABE8e^4cv^-mediated therapeutic editing, we again injected P21 juvenile male *mdx^4cv^* mice with 4E13 VG/kg of MyoAAV-SaABE8e^4cv^ or MyoAAV-SaABE8e^NT^ and evaluated these mice at 6 months post-injection. As shown in **Fig. 3A-B**, compared to mice harvested at 1-month post-injection, the frequencies of dystrophin-expressing cells were lower in TA (6.6 ± 5.3%) but comparable in diaphragm (13.2 ± 4.3%) and heart (36.9 ± 4.1%). Unexpectedly, western blot analysis showed that dystrophin protein levels were markedly lower in the TA and diaphragm of MyoAAV-SaABE8e^4cv^-injected mice at this time point (1.4 ± 1.1% of WT levels in TA and 0.3 ± 0.1% of WT levels in diaphragm at 6 months versus 10.5 ± 8.0% in TA and 8.1 ± 6.6% in diaphragm at 1 month), whereas dystrophin levels in heart remained comparable or even increased (32.8 ± 10.8% of WT, **Fig. 3C and Fig. S10**) at 6 months versus 1 month (16.8 ± 7.1% of WT, **Fig. 2C and Fig. S4**) post-injection.

**Fig. 3.**
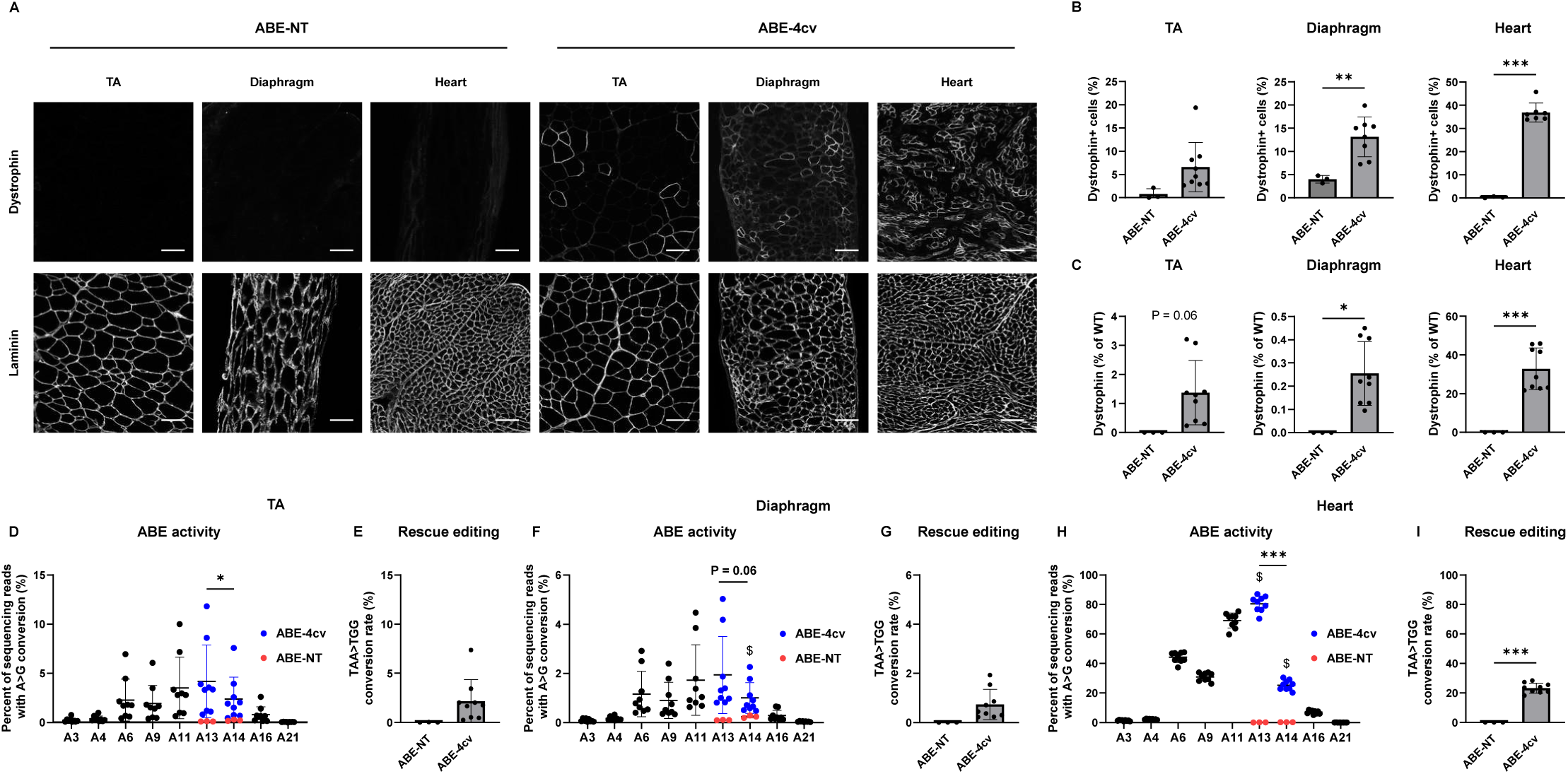
Systemic delivery of MyoAAV-SaABE8e^4cv^ in *mdx^4cv^* mice sustains production of edited Dystrophin protein through adulthood. Juvenile (P21) male *mdx^4cv^*mice were injected retro-orbitally with 4E13 VG/kg MyoAAV-SaABE8e^4cv^ or MyoAAV-SaABE8e^NT^, and tissues harvested six months later. (A) Representative single-channel immunofluorescence images from ABE-NT or ABE-4cv injected mice showing the same field of tibialis anterior (TA), diaphragm, or heart stained for dystrophin (top) or laminin (bottom), as indicated. Scale bar = 100 µm. (B) Quantification of dystrophin-positive myofibers or cardiomyocytes, calculated as percent (%) of total cells (derived from laminin staining). (C) Semi-quantitative analysis of dystrophin protein expression in TA, diaphragm, and heart of mice injected with MyoAAV-SaABE8e^NT^ or MyoAAV-SaABE8e^4cv^. Representative blots are shown in Fig. S10. (D-I) cDNA from TA (D-E), diaphragm (F-G), or heart (H-I) were analyzed by amplicon sequencing at the *mdx^4cv^*mutation site. Representative allele frequency tables are shown in Fig. S11. (D,F,H) A>G conversion rates for individual adenines (A) of MyoAAV-SaABE8e^4cv^-injected *mdx^4cv^* mice were calculated from sequencing reads. Adenine conversions in the *mdx^4cv^* premature stop codon from MyoAAV-SaABE8e^4cv^-injected mice are shown in dark blue (A13 and A14). A13 and A14 A>G conversion rates from control, MyoAAV-SaABE8e^NT^-injected *mdx^4cv^* mice are shown in red. $ indicates a significant difference between MyoAAV-SaABE8e^4cv^- and MyoAAV-SaABE8e^NT^-injected *mdx^4cv^* mice in A>G conversion rates on the indicated adenine (determined by unpaired T-test. (E,G,I) Percent of reads in which the *mdx^4cv^*nonsense mutation is converted to a non-stop (missense) mutation (TAA>TGG). n = 3-9 biological replicates/group; B, C, E, G, and I were analyzed with unpaired T-test, D, F, and H were analyzed with one-way repeated measures ANOVA. Data are shown as mean ± SD. *, **, and *** indicate P < 0.05, < 0.01, and <0.001, respectively.

To quantify the persistence of MyoAAV-SaABE8e^4cv^-mediated edits, we sequenced genomic DNA and cDNA from the TA, diaphragm, and heart of *mdx^4cv^* mice injected with MyoAAV-SaABE8e^4cv^ or MyoAAV-SaABE8e^NT^. Similar to results at one-month post-MyoAAV-SaABE8e^4cv^ injection, data from *in vivo* edited TA, diaphragm, and heart indicated that the A>G conversion rate for A14 was still significantly lower than at A13 for both cDNA (**Fig. 3D, F, H**) and genomic DNA (**Fig. S12**). Mirroring results from Western blotting, MyoAAV-SaABE8e^4cv^-mediated edits detected in cDNA from the TA and diaphragm were substantially lower in mice analyzed at 6 months after AAV injection, as compared with mice analyzed at 1 month, whereas editing rates in heart cDNA were comparable. Specifically, the A>G conversion rates at A13 detected in cDNA were 4.2 ± 3.7% in the TA, 1.9 ± 1.6% in the diaphragm, and 80.4 ± 5.3% in the heart (**Fig. 3D, F, H**). Regarding rescue editing rates, we detected 2.2 ± 2.2% in TA, 0.7 ± 0.6% in Diaphragm, and 23.3 ± 3.4% in Heart (**Fig. 3E, G, I**).

### Inefficient rescue of dystrophin deficiency in neonatal *mdx^4cv^* mice following systemic delivery of MyoAAV-SaABE8e^4cv^

Duchenne Muscular Dystrophy (DMD) is a progressive disease, and symptoms appear in the majority of DMD patients around age 2-4^26^. Treating early in disease progression could pause DMD pathology at a milder stage and thus might more effectively rescue dystrophic symptoms^26,27^. To assess the effects of MyoAAV-SaABE8e^4cv^-mediated dystrophin restoration in younger mice, we injected neonatal *mdx^4cv^* mice (P3) with 4E13 VG/kg of MyoAAV-SaABE8e^4cv^ or MyoAAV-SaABE8e^NT^ through intraperitoneal (IP) or retro-orbital (RO) injection. One month after injection, IP delivery of MyoAAV-SaABE8e^4cv^ led to 5.6 ± 1.6% of myofibers in TA, 33.2 ± 7.6% of myofibers in the diaphragm, and 17.8 ± 6.4% of cardiomyocytes becoming dystrophin-positive (**Fig. 4A-B**). RO delivery of MyoAAV-SaABE8e^4cv^ led to 10.1 ± 2.8% of myofibers in TA, 17.7 ± 5.7% of myofibers in the diaphragm, and 17.4 ± 6.3% of cardiomyocytes to become dystrophin-positive (**Fig. 4A-B**). Western blot analysis demonstrated that IP delivery of MyoAAV-SaABE8e^4cv^ rescued dystrophin protein production in MyoAAV-SaABE8e^4cv^-injected *mdx^4cv^* mice to levels 0.9 ± 0.5%, 50.9 ± 13.7%, and 14.3 ± 8.1% of the WT in TA, diaphragm and heart, respectively (**Fig. 4C and Fig. S13).** On the other hand, RO delivery of MyoAAV-SaABE8e^4cv^ rescued dystrophin levels to 0.7 ± 0.5%, 14.9 ± 9.2%, and 5.3 ± 3.3% of the WT in TA, diaphragm, and heart, respectively (**Fig 4C and Fig. S13**).

**Fig. 4.**
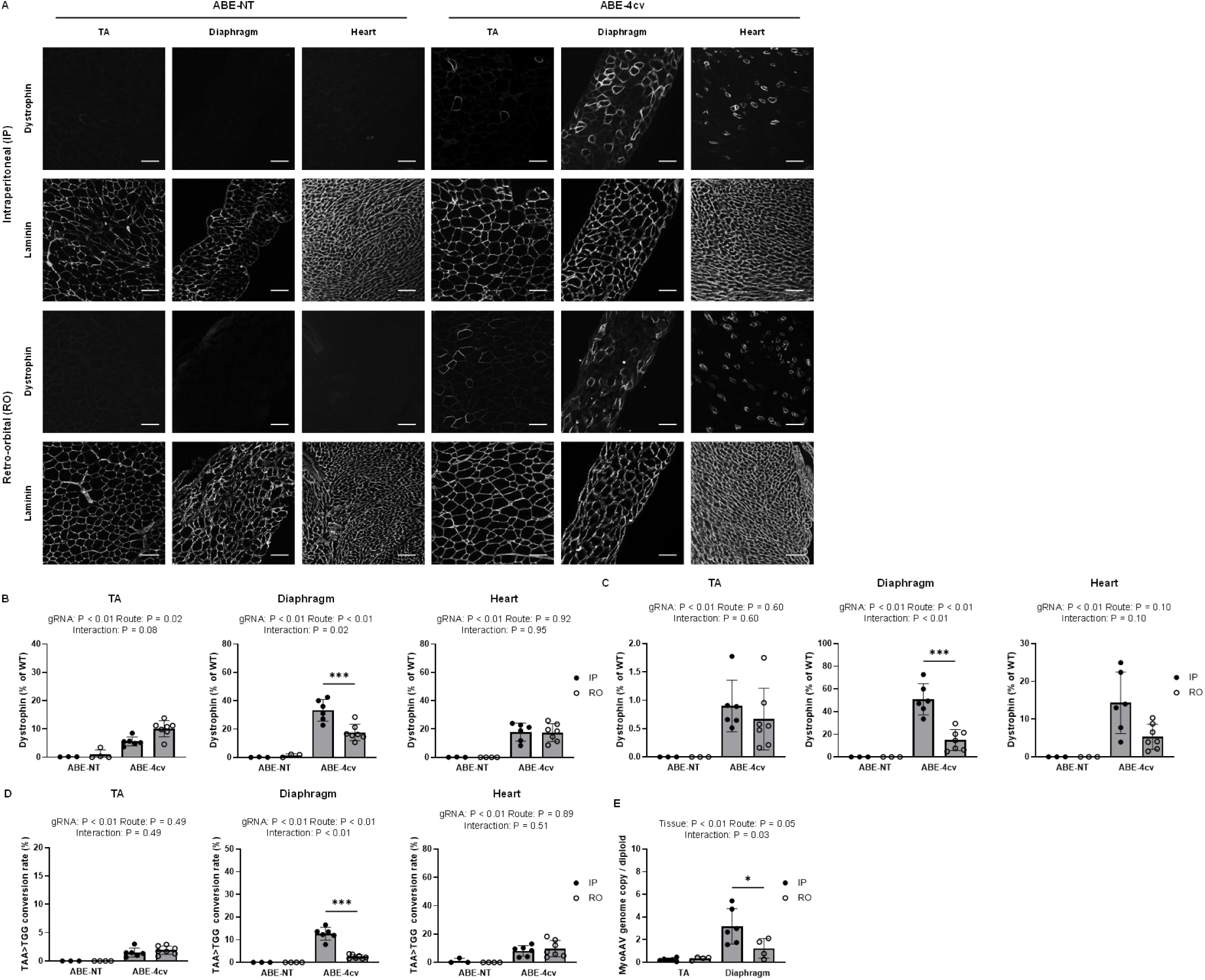
Inefficient rescue of dystrophin deficiency in TA and heart following systemic delivery of MyoAAV-SaABE8e^4cv^ in neonatal *mdx^4cv^* mice. Neonatal (P3) male *mdx^4cv^* mice were injected intraperitoneally (IP) or retro-orbitally (RO) with 4E13 VG/kg MyoAAV-SaABE8e^4cv^ (ABE-4cv) or MyoAAV-SaABE8e^NT^ (ABE-NT), and tissues were harvested one month later. (A) Representative single-channel immunofluorescence images from ABE-NT or ABE-4cv injected mice showing the same field of tibialis anterior (TA), diaphragm, or heart stained for dystrophin (top) or laminin (bottom), as indicated. Scale bar = 100 µm. (B) Quantification of dystrophin-positive myofibers or cardiomyocytes, calculated as percent (%) of total cells (derived from laminin staining). (C) Semi-quantitative analysis of dystrophin protein expression in TA, diaphragm, and heart of mice injected with MyoAAV-SaABE8e^NT^ or MyoAAV-SaABE8e^4cv^. Representative blots are shown in Fig. S13. (D) cDNA from TA, diaphragm, and heart was analyzed by amplicon sequencing at the *mdx^4cv^* mutation site. A>G conversion rates of individual adenines are shown in Fig. S14, and representative allele frequency tables are shown in Fig. S15. Percent of reads in which the *mdx^4cv^* nonsense mutation is converted to a non-stop (missense) mutation (TAA>TGG) is shown for the indicated conditions. (E) Ratio of MyoAAV vector genome copy number to diploid genome for MyoAAV-SaABE8e^4cv^ IP or RO injected *mdx^4cv^* mice in TA and diaphragm. n = 3-7 biological replicates/group; B-E were analyzed with two-way ANOVA. Data are shown as mean ± SD. *, **, and *** indicate P < 0.05, < 0.01, and <0.001, respectively.

In contrast to the TA and heart, in which no obvious differences in dystrophin restoration were detected between IP and RO MyoAAV-SaABE8e^4cv^ injected *mdx^4cv^* mice, IP injection restored substantially greater dystrophin expression than RO injection in the diaphragm (**Fig. 4A-C**). Interestingly, albeit sequencing of cDNA from TA, diaphragm, and heart showed comparable rescue editing rates in the TA and heart between IP and RO injected *mdx^4cv^* mice, rescue editing rates was significantly higher in the diaphragm following IP delivery of MyoAAV-SaABE8e^4cv^ (**Fig. 4D**). Further analysis of adenine conversion across all targetable positions within the 4cv-gRNA1 targeting site revealed that A14 conversion at both cDNA and genomic DNA levels remained significantly lower than A13 and therefore represented the limiting factor for rescue editing under both delivery routes (**Fig. S14**). These results indicate an overall increased adenine conversion efficiency in the diaphragm following IP injection, likely due to apposition of this muscle to the peritoneal cavity, exposing it to a higher concentration of virus with this route of delivery.

To explore this possibility, we quantified the MyoAAV-SaABE8e^4cv^ vector genomes (VGs) present in the TA and diaphragm of IP versus RO MyoAAV-SaABE8e^4cv^ injected *mdx^4cv^* mice. As shown in **Fig. 4E**, delivery of 4E13 VG/kg of MyoAAV-SaABE8e4cv via IP injection resulted in VG to diploid ratios of 0.24 ± 0.14 in TA and 3.18 ± 1.57 in diaphragm, whereas RO injection yielded ratios of 0.34 ± 0.14 in TA and 1.24 ± 0.87 in diaphragm. These results demonstrate that IP delivery in P3 *mdx^4cv^*mice achieved higher AAV transduction efficiency than RO delivery in the diaphragm. In addition, both delivery routes produced substantially higher transduction in the diaphragm than in the TA muscle, which positively correlates with the enhanced dystrophin restoration observed in the diaphragm (**Fig. 4A–C**). Overall, our data demonstrates that the copy number of MyoAAV-SaABE8e^4cv^ genomes retained within cells in target tissues is a major determinant of dystrophin restoration in treated P3 *mdx^4cv^*mice.

As previous studies have demonstrated efficient transduction of neonatal mouse muscle by conventional and myotropic AAVs^28,29^, we hypothesized that the rapid dilution of AAV genomes in fast-growing neonatal skeletal muscle and heart might underlie the low dystrophin restoration observed following MyoAAV-SaABE8e^4cv^ delivery to P3 *mdx^4cv^*mice. To test this hypothesis, we administered 4E13 VG/kg of MyoAAV-SaABE8e^4cv^ or MyoAAV-SaABE8e^NT^ to P7 *mdx^4cv^* mice via RO injection. At P7, cardiomyocytes have largely lost the ability to divide^30^, whereas myofibers continue to rapidly acquire new myonuclei^31^. Consistent with our hypothesis, dystrophin restoration in TA and diaphragm remained minimal; however, 30.3 ± 4.7% of cardiomyocytes in P7-injected animals showed restored dystrophin expression, and total dystrophin levels in the heart reached 16.4 ± 5.3% of the WT (**Fig. S16A–C**). The same pattern was also observed in SaABE8e-mediated genome editing rates, in which we detected A13 conversion rates of 5.6 ± 2.3% in TA, 6.0 ± 1.2% in diaphragm, and 62.1 ± 4.9% in heart at cDNA level (**Fig. S18**). For rescue editing rates, we detected 1.5 ± 0.7% in TA, 1.7 ± 0.5% in diaphragm, and 16.2 ± 2.1% in heart (**Fig. S16D**). Notably, despite the superior rescue editing in the P7 heart, ABE-mediated rescue of the *mdx^4cv^* mutation remained limited by low A14 A>G conversion in both cDNA and genomic DNA analyses (**Fig. S18**).

### Diminished efficiency of dystrophin restoration in young adult *mdx^4cv^*mice systemically administered MyoAAV-SaABE8e^4cv^

In both DMD patients and rodent models, dystrophic symptoms and complications, such as the number of necrotic myofibers, muscle fibrotic progression, lipid deposition, and inflammatory response, worsen progressively with age^32,33^. These symptoms not only create physical barriers for AAV delivery^34^, but also lead to irreversible muscle damage. To examine whether the efficiency of dystrophin restoration by base editing is diminished when older mice are treated with MyoAAV-SaABE8e^4cv^, we injected 12-week-old (12W) *mdx^4cv^* mice, which are generally considered equivalent to ∼20-30-year-old humans^35^, retro-orbitally with 4E13 VG/kg of MyoAAV-SaABE8e^4cv^ or MyoAAV-SaABE8e^NT^. Quantification of immunostained tissue sections indicated that 19.5 ± 7.1% of myofibers in the TA, 16.8 ± 6.1% of myofibers in the diaphragm, and 22.7 ± 4.2% of cardiomyocytes in MyoAAV-SaABE8e^4cv^ injected *mdx^4cv^* mice were dystrophin-positive at one month post-injection (**Fig. 5A-B**).

**Fig. 5.**
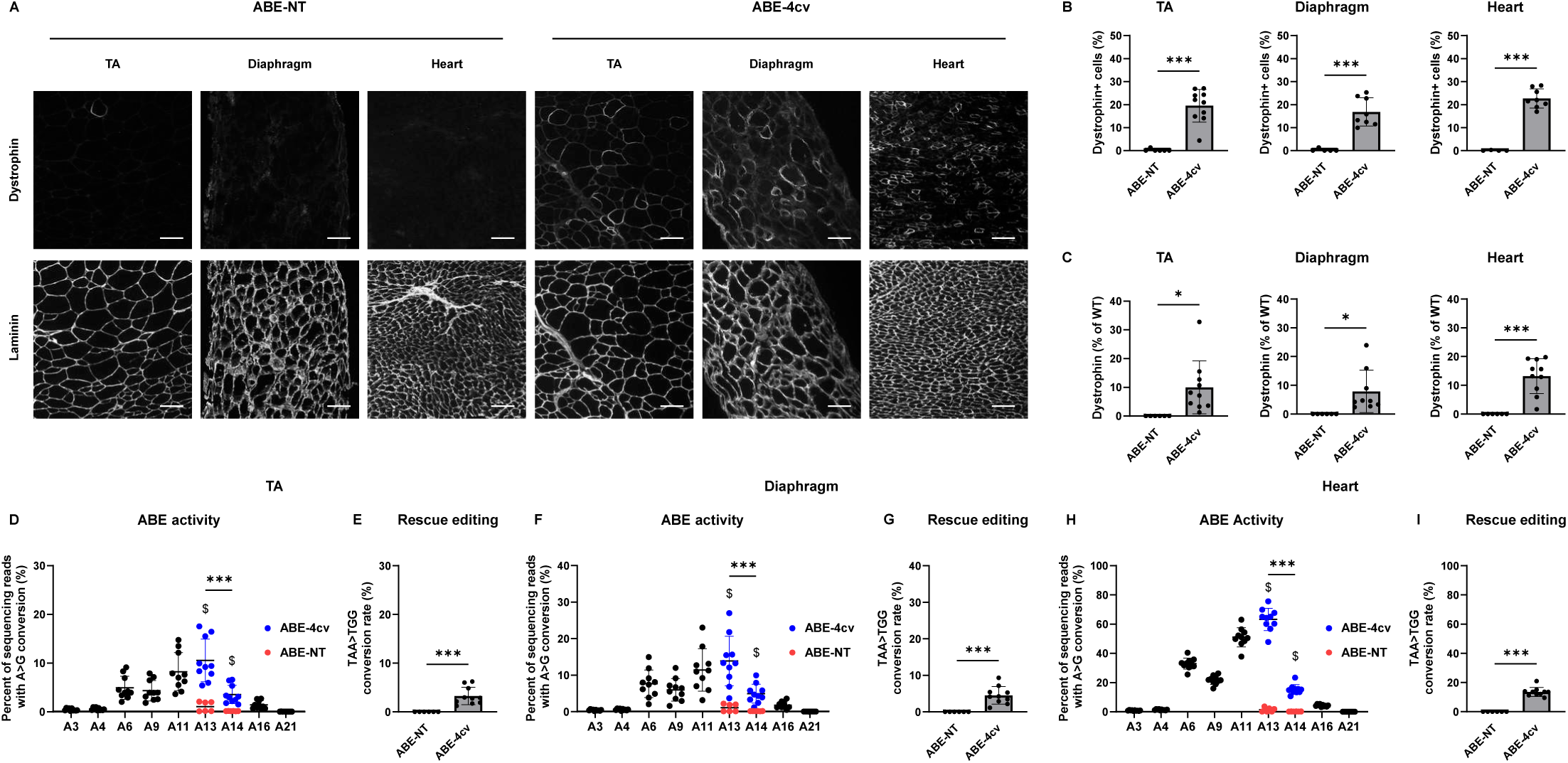
Diminished efficiency of dystrophin restoration following systemic delivery of MyoAAV-SaABE8e^4cv^ to young adult *mdx^4cv^* mice. Young adult (12-week-old) male *mdx^4cv^* mice were injected retro-orbitally (RO) with 4E13 VG/kg MyoAAV-SaABE8e^4cv^ or MyoAAV-SaABE8e^NT^, and tissues harvested one month later. (A-B) Representative single-channel immunofluorescence images from ABE-NT or ABE-4cv injected mice showing the same field of tibialis anterior (TA), diaphragm, or heart stained for dystrophin (top) or laminin (bottom), as indicated. Scale bar = 100 µm. (B) Quantification of dystrophin-positive myofibers or cardiomyocytes, calculated as percent (%) of total cells (derived from laminin staining). (C) Semi-quantitative analysis of dystrophin protein expression in TA, diaphragm, and heart of mice injected with MyoAAV-SaABE8e^NT^ or MyoAAV-SaABE8e^4cv^. Representative blots are shown in Fig. S20. (D-I) cDNA from TA (D-E), diaphragm (F-G), and heart (H-I) were analyzed by amplicon sequencing at the *mdx^4cv^*mutation site. Representative allele frequency tables are shown in Fig. S22. (D,F,H) A>G conversion rates for individual adenines (A) of MyoAAV-SaABE8e^4cv^-injected *mdx^4cv^* mice were calculated from sequencing reads. Adenine conversions in the *mdx^4cv^*premature stop codon from MyoAAV-SaABE8e^4cv^-injected mice are shown in dark blue (A13 and A14). A13 and A14 A>G conversion rates from control, MyoAAV-SaABE8e^NT^-injected *mdx^4cv^* mice are shown in red. $ indicates a significant difference between MyoAAV-SaABE8e^4cv^- and MyoAAV-SaABE8e^NT^-injected *mdx^4cv^* mice in A>G conversion rates on the indicated adenine (determined by unpaired T-test. (E,G,I) Percent of reads in which the *mdx^4cv^*nonsense mutation is converted to a non-stop (missense) mutation (TAA>TGG). n = 6-10 biological replicates/group; B, C, E, G, and I were analyzed with unpaired T-test, D, F, and H were analyzed with one-way repeated measures ANOVA. Data are shown as mean ± SD. *, **, and *** indicate P < 0.05, < 0.01, and <0.001, respectively.

Western blot analysis determined dystrophin rescue in MyoAAV-SaABE8e^4cv^ injected *mdx^4cv^* mice to levels 10.0 ± 9.2%, 7.8 ± 7.4%, and 13.2 ± 6.1% of WT, in the treated TA, diaphragm, and heart, respectively (**Fig. 5C and Fig. S20**). For SaABE8e-mediated genome editing rates, we again sequenced genomic DNA and cDNA from TA, diaphragm, and heart. In the TA, diaphragm, and heart, the A>G conversion rates at A14 were still significantly lower than at A13 for both cDNA (**Fig. 5D, F, H**) and genomic DNA (**Fig. S21**). We detected A13 conversion rates of 10.6 ± 4.4% in TA, 13.9 ± 6.8% in diaphragm, and 63.3 ± 7.6% in heart at the cDNA level (**Fig. 5D, F, H**). For rescue editing, we detected 3.3 ± 1.8% in TA, 4.5 ± 2.4% in diaphragm, and 13.5 ± 3.1% in heart (**Fig. 5E, G, I**).

To further assess the therapeutic window for MyoAAV-SaABE8e^4cv^-based gene therapy in this dystrophic model, we injected 6-month-old (6M) *mdx^4cv^*mice with 4E13 VG/kg of MyoAAV-SaABE8e^4cv^ or MyoAAV-SaABE8e^NT^ via RO injection. Quantitative analysis of dystrophin immunostaining indicated that 9.2 ± 3.2% of myofibers in TA, 10.2 ± 3.9% of myofibers in diaphragm, as well as 13.6 ± 2.2% cardiomyocytes of MyoAAV-SaABE8e^4cv^ injected *mdx^4cv^* mice were dystrophin-positive (**Fig. S23A-B**). Western blotting showed dystrophin restoration to 1.7 ± 1.0%, 9.6 ± 5.0%, and 6.1 ± 1.0% of WT levels in TA, diaphragm, and heart, respectively, following MyoAAV-SaABE8e4cv treatment of *mdx^4cv^* mice (**Fig. S23C and Fig. S24**). For SaABE8e-mediated genome editing rates, the A>G conversion rates at A14 were still significantly lower than at A13 at both cDNA (**Fig. S23D, F, H**) and genomic DNA (**Fig. S25**) levels in TA, diaphragm, and heart. We determined A13 conversion rates of 8.8 ± 4.9% in TA, 14.0 ± 2.4% in diaphragm, and 55.0 ± 10.7% in heart, as well as rescue editing rates of 2.4 ± 1.7% in TA, 4.4 ± 1.2% in diaphragm, and 13.0 ± 7.5% in heart (**Fig. S23D-I**).

### Systemic delivery of MyoAAV-SaABE8e^4cv^ achieved rescue editing in muscle satellite cells, with the highest efficiency observed in juvenile and early young adult treated *mdx^4cv^* mice

The absence of dystrophin in DMD patients and rodent models leads to myofiber fragility and degeneration, which in turn activates satellite cell-mediated regeneration. However, because the newly formed myofibers still lack dystrophin, they remain vulnerable to further damage, resulting in recurrent cycles of degeneration and regeneration. In *mdx^4cv^* mice, this regenerative/degenerative cycle peaks between 3 and 8 weeks of age. Therefore, if a gene therapy strategy fails to recode the *mdx^4cv^* mutation in satellite cells, the durability of dystrophin restoration in myofibers would be expected to diminish over time. To examine whether delivery of MyoAAV-SaABE8e^4cv^ affected rescue editing in satellite cells, we FACSorted these cells, using previously validated methods^24^, from *mdx^4cv^*mice that were RO-injected with MyoAAV-SaABE8e^4cv^ at P3, P7, P21, 12W, or 6M of age. As a control condition, we also FACSorted satellite cells from age-matched *mdx^4cv^* mice RO-injected with MyoAAV-SaABE8e^NT^. Genomic DNA from these FACSorted satellite cells was extracted for amplicon sequencing at the *mdx^4cv^* mutation site.

These analyses revealed a significant difference in rescue editing rates between satellite cells from *mdx^4cv^* mice injected with MyoAAV-SaABE8e^4cv^ and MyoAAV-SaABE8e^NT^ (**Fig. 6A**). Supporting this result, as shown in **Fig. 6B-F** and **Fig. S27**, reads bearing A>G conversions were detected in satellite cells from mice of all ages, including from P3 MyoAAV-SaABE8e^4cv^ injected mice, which showed the lowest A13 A>G conversion rate (0.14 ± 0.07%). Additionally, the age of MyoAAV-SaABE8e^4cv^ injection also has a main effect on rescue editing rates. Compared with age-matched controls, injected with MyoAAV-SaABE8e^NT^, injection of MyoAAV-SaABE8e^4cv^ yielded significantly higher rescue editing rates in satellite cells from *mdx^4cv^* mice injected at P21 (2.73 ± 1.46% rescued edited reads) and 12W (1.48 ± 1.53% rescue edited reads) (**Fig. 6B, C**). Interesting, in satellite cells from P21 injected mice, A13 and A14 conversion rates were comparable (**Fig. 6B**), similar to the editing outcomes observed following SaABE8e^4cv-gRNA1^ plasmid transfection-induced in these cells *ex vivo* (**Fig. 1C**). In contrast, as shown in **Fig. 6C**, satellite cells from 12W-injected mice displayed an editing pattern similar to that observed in whole muscle tissue, in which the A>G conversion rate at A14 (1.84 ± 1.95%) was significantly lower than at A13 (3.28 ± 2.85%). Collectively, these results demonstrate that systemic delivery of MyoAAV-SaABE8e^4cv^ induces SaABE8e-mediated genome editing in satellite cells in *mdx^4cv^* mice, with the highest rescue editing efficiencies observed in mice treated at juvenile and young adult stages.

**Fig. 6.**
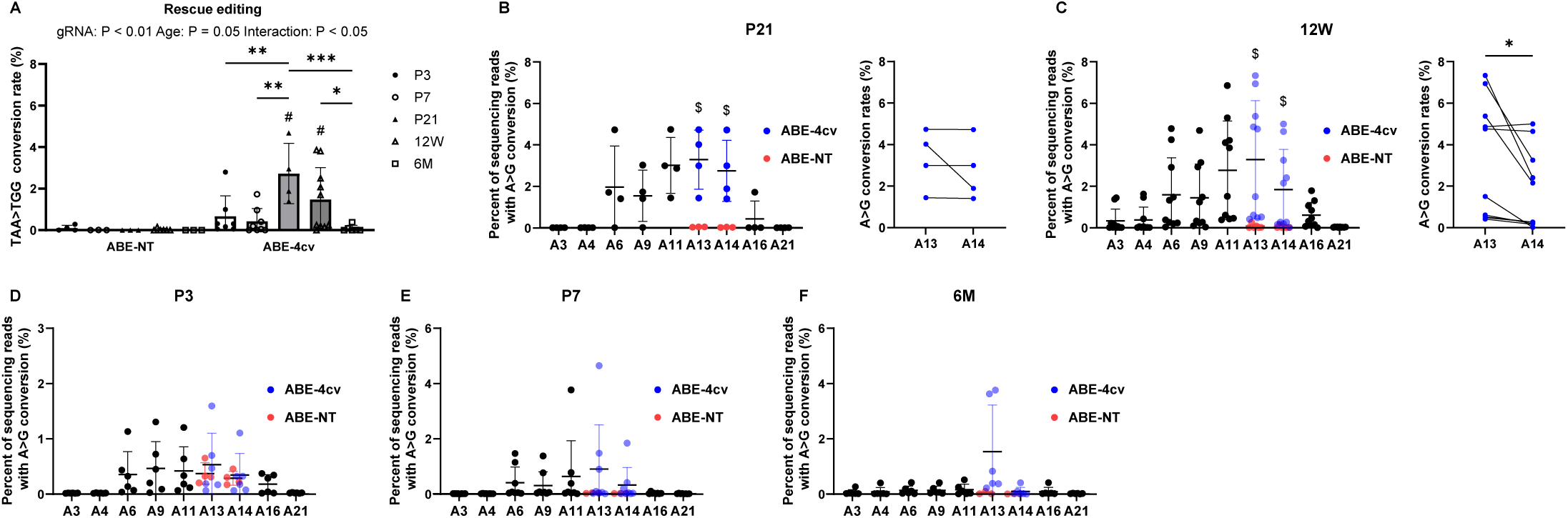
Rescue editing of *mdx^4cv^* mutation in muscle satellite cells following systemic delivery of MyoAAV-SaABE8e^4cv^. 4E13 VG/kg MyoAAV-SaABE8e4cv or MyoAAV-SaABE8eNT were injected RO into P3, P7, P21, 12W, or 6M *mdx^4cv^* mice. One month post injection, satellite cells were FACS-isolated (Fig. S1), and genomic DNA extracted for amplicon sequencing at the *mdx^4cv^* mutation site. (A) Percent of reads in which the *mdx^4cv^* nonsense mutation is converted to a non-stop (missense) mutation (TAA>TGG). (B-F) A>G conversion rates of *mdx4cv* mice injected with MyoAA-SaABE8e4cv at the indicated age, shown for individual adenines (A) of MyoAAV-SaABE8e^4cv^-injected *mdx^4cv^* and MyoAAV-SaABE8e^NT^-injected mice. Percentages calculated from sequencing reads. Adenine conversions in the *mdx^4cv^* premature stop codon from MyoAAV-SaABE8e^4cv^-injected mice are shown in dark blue (A13 and A14). A13 and A14 A>G conversion rates from MyoAAV-SaABE8e^NT^-injected *mdx^4cv^* mice are shown in red. $ indicates a significant difference between MyoAAV-SaABE8e^4cv^- and MyoAAV-SaABE8e^NT^-injected *mdx^4cv^* mice in A13 or A14 A>G conversion rates (determined by unpaired T-test). For B and C, the paired A13 and A14 conversion rates for each mouse (same data as in adjacent graph) are shown on the right. Representative allele frequency tables are shown in Fig. S27. n = 3-10 biological replicates/group; A is analyzed with two-way ANOVA, and B and C are analyzed with paired T-test. Data are shown as mean ± SD. *, **, and *** indicate P < 0.05, < 0.01, and <0.001, respectively.

## Discussion

In this study, we leveraged a newly developed size-minimized ABE^20^ together with a muscle-tropic AAV (MyoAAV)^21^ to establish a single-vector *in vivo* gene correction strategy for dystrophin-deficient muscular dystrophy in *mdx^4cv^*mice. This ABE strategy is expected to be safer than CRISPR-Cas9-mediated pathogenic exon excision approaches^8^, as ABE induces substantially fewer large deletions and chromosomal translocations in human cells^36^. Additionally, compared with older versions of ABEs that are too large to be efficiently packaged in a single AAV, and therefore require an intein-split dual-AAV delivery strategy for reconstitution^16^, this size-minimized SaABE8e enables a single-vector design, which reduces the total dose of AAV required for therapy and simplifies vector design and production. Use of MyoAAV additionally enhances skeletal and cardiac muscle transduction while minimizing liver targeting, further improving safety and efficacy. Altogether, this proof-of-concept pre-clinical study establishes a new, effective, and likely safer gene correction strategy for DMD.

To correct the *mdx^4cv^* mutation with ABE, we designed a gRNA targeting the *mdx^4cv^* TAA premature stop codon (4cv-gRNA1). MyoAAV delivery of this gRNA, along with ABE (MyoAAV-SaABE8e^4cv^), successfully converted the *mdx^4cv^* premature stop codon (TAA) to a tryptophan coding sequence (TGG). Importantly, this change requires simultaneous conversion of both adenines to guanines to restore a full-length coding sequence as single-base edits, yielding TAG or TGA codons, which still encode termination signals. This unfavorable editing context renders ∼70% of SaABE8e^4cv^ editing events non-therapeutic in muscle. Despite this limitation, we achieved readily detectable rescue of the nonsense codon to generate a tryptophan-substituted sequence in striated muscles and satellite cells of juvenile male *mdx^4cv^* mice. This strategy supported restoration of dystrophin and DAPC components in up to 30.8 ± 6.4% of myofibers and 33.7 ± 6.2% of cardiomyocytes and improved gastrocnemius muscle function by 25% in treated mice within one month after injection (**Fig. 2 and Fig. S5**). Notably, the suboptimal target sequence also provided a stringent framework for comparing therapeutic efficacy across neonatal, juvenile, and young adult mice. Our findings identify the juvenile stage (P21 in mice) as optimal for therapeutic efficacy and reveal a significant impact of age in reducing editing efficiency in both neonatal and older adult animals.

Compared with juvenile *mdx^4cv^* mice injected retro-orbitally with MyoAAV-SaABE8e^4cv^, *mdx^4cv^* mice injected as neonates (P3) exhibited substantially lower rates of mutation recoding in satellite cells, TA, and heart (**Fig. 4**). As AAV genomes predominantly persist as non-replicating episomes within host nuclei and become diluted during cell division^6^, we speculate that rapid AAV genome dilution in actively growing neonatal tissues might be the cause of the inefficient *mdx^4cv^* mutation recoding rates. Aligned with this theory, both satellite cells^37^ and cardiomyocytes^38^ are highly proliferative at P3. Although myofibers are generally considered post-mitotic, rapid myonuclear accretion during neonatal growth can be expected to substantially reduce the proportion of myonuclei harboring TAA>TGG therapeutic edit in each myofiber if those new myonuclei must draw on a pool of poorly edited satellite cells as their source. Given the restriction of dystrophin expression to discrete myonuclear domains^39^, diminution of the proportion of myonuclei bearing TAA>TGG therapeutic edits can be expected to result in a smaller proportion of each myofiber exhibiting dystrophin-positivity. Prior work has also shown that both transgene expression and the ratio of AAV vector genomes to diploid host genome progressively decreases when AAV is administered during periods of heightened satellite cell proliferation, such as following muscle injury^40^. Unexpectedly, in our studies, AAV-to-host genome ratio was 3.6-fold higher in diaphragm than in TA following RO delivery, correlating with higher *mdx^4cv^* recoding and dystrophin restoration rates (**Fig. 4**). IP delivery further increased the AAV-to-host genome ratio to 9.4-fold higher in the diaphragm than in TA and correlated with even further enhanced *mdx^4cv^* recoding and dystrophin restoration (**Fig. 4**). These findings suggest that the AAV genome copy number retained within target cells is a critical determinant of efficient gene recoding in neonatal muscle tissues. Accordingly, employing combined or optimized AAV delivery routes to achieve higher local AAV transduction rate may represent a practical strategy to improve therapeutic efficacy in neonatal tissues.

DMD is a progressive disease in both human patients and pre-clinical rodent models. In *mdx^4cv^* mice, the nearly complete loss of full-length dystrophin and disruption of the DAPC results in a more severe and rapid dystrophic progression than occurs in the more widely used *Dmd* exon 23 *mdx* mice (mdx-23)^41^, with *mdx^4cv^* mice exhibiting detectable myofibrosis, myonecrosis, chronic inflammation, and lipid deposition^42^. While disease progression is already pronounced in muscle by 12 weeks of age^43^, cardiomyopathy typically develops later, between 6 and 9 months of age ^30^. The onset of these complications is expected to compromise the effectiveness of AAV-mediated gene therapy applied at later stages of the disease, due to irreversible muscle damage and increased collagen and lipid depositions posing a physical barrier for AAV delivery. Consistent with this concept, we observed a gradual decrease in *mdx^4cv^* mutation recoding rate and dystrophin-positive cell frequency in adult *mdx^4cv^* mice treated at older ages (**Fig. 5** and **Fig. S23**). In contrast to satellite cells from P21 MyoAAV-SaABE8e^4cv^ injected mice, which showed comparable A13 and A14 A>G conversion rates (**Fig. 6C**), the A14 A>G conversion rate (1.84 ± 1.95%) was significantly lower than the A13 A>G conversion rate (3.28 ± 2.85%) in satellite cells from 12W treated *mdx^4cv^* mice (**Fig. 6D**), rendering A14 A>G conversion the limiting factor for rescue editing. Considering that DMD satellite cells develop cell-autonomous myogenic defects^4^, this disparity suggests that disease progression within satellite cells may also influence genome-editing outcomes. Overall, our findings suggest that treatment at the young adult stage in *mdx^4cv^* mice represents a transitional window during which the therapeutic effectiveness of MyoAAV-SaABE8e^4cv^ begins to decrease in an age-dependent manner. Interventions that slow dystrophic progression might extend this treatable window and enhance long-term therapeutic benefit.

To test the persistence of MyoAAV-SaABE8e^4cv^-mediated editing and therapeutic effect, we analyzed tissues from juvenile-treated mice at one and six months after MyoAAV-SaABE8e^4cv^ delivery. Overall, we found that dystrophin restoration remained persistent in the heart but became lower in TA and diaphragm at six months post treatment compared to one-month post-treatment (**Fig. 2** and **Fig. 3**). Considering that the persistence of dystrophin-editing depends on dystrophin levels in skeletal muscle but not heart^44^, this result suggests that a higher initial dystrophin restoration rate is required for a long-term therapeutic effect in skeletal muscle. As previously noted, ∼70% of MyoAAV-SaABE8e^4cv^ introduced edits were not therapeutic because both A13 and A14 must be edited to bypass the *mdx^4cv^*mutation. Thus, it may be easier to achieve long-term therapeutic effects with a single-vector ABE strategy if the sequence targeted allows for a more efficient mutation recoding rate (i.e., requiring only a single adenine edit). Notably, although both dystrophin expression levels and *mdx^4cv^* editing rates in the TA and diaphragm were much lower at 6 months than at 1 month after MyoAAV-SaABE8e^4cv^ delivery (**Fig. 2** and **Fig. 3**), the frequency of dystrophin-positive myofibers in muscle cross-sections remained comparable in the diaphragm. Given that AAV-mediated gene editing results in segmental dystrophin restoration^39^, it is possible that dystrophin expression levels in individual myofibers of both TA and diaphragm are reduced over time, but there is a higher probability of detecting a dystrophin-positive region in a given cross-sectional plane of myofibers in the diaphragm. This theory is supported by the concept of dystrophin myonuclear domain^39^: because myofibers in the diaphragm generally have higher myonuclear densities^45^, the degree of overlap between basal sarcolemmal dystrophin units (BSDU, sarcolemma-associated dystrophin protein organized in focal and immobile nuclear domains) is expected to be greater in the diaphragm than in limb muscles.

Altogether, these data provide proof-of-principle evidence in a disease-relevant preclinical model that a single MyoAAV vector-delivered, ABE-mediated mutation-recoding strategy can correct the genetic lesions that give rise to dystrophic pathologies and improve muscle phenotype and function, even when the target sequence is not optimal for base-editing-mediated strategies. Given the heterogeneity of mutations in DMD patients and the known challenge that target sequences in patients do not always permit the most efficient designs for therapeutic base editing, these findings offer clinically relevant insights and establish a useful baseline for the continuing development of advanced genomic medicine approaches to enable curative therapies for DMD and related muscle pathologies.

## Methods

### Mice

Mice (B6Ros.Cg-*Dmd^mdx-4Cv^*/J, JAX stock #002378; C57BL/6J, JAX stock #000664) were purchased from the Jackson Laboratory and housed and maintained at the Harvard Biological Research Infrastructure in accordance with animal use guidelines. All mice were given food and water ad libitum and kept in a room maintained at 25 °C with a 12-h light: dark cycle. All experimental protocols were approved by the Harvard University Institutional Animal Care and Use Committee (IACUC).

### Plasmid construction

The AAV-EFS-SaABE8eV106W-bGH-U6-sgRNA-BsmBI was a gift from David Liu (Addgene plasmid # 189924 ; http://n2t.net/addgene:189924 ; RRID:Addgene_189924). gRNA targeting *mdx^4cv^* premature stop codon (4cv-gRNA1: 5’ AGAACAGGAGACAACAGTTGA 3’ or 4cv-gRNA2: 5’ CTGCAGAACAGGAGACAACAG 3’) or a non-targeting gRNA (GCTTTCACGGAGGTTCGACG) was cloned with HindIII (NEB) and NdeI (NEB) to replace the original gRNA sequence in the parental plasmid. Possible 4cv-gRNA off-targets were predicted with an algorithm designed by Hsu et al^46^.

### Production of AAV

MyoAAV vectors were generated as previously described^47^. Briefly, HEK293 cells cultured in Dulbecco’s modified eagle’s medium (DMEM, ThermoFisher Scientific) with 10% fetal bovine serum (FBS, GeminiBio) and 1% Penicillin-Streptomycin (Pen-Strep, ThermoFisher Scientific) were plated in 15 cm dishes and expanded in culture until reaching ∼70% confluency. Then, four 15 cm dishes of HEK293 cells were seeded into each HYPERFlask cell culture vessel. Two days later, each HYPERFlask was transfected with 260 µg of the d16 helper plasmid, 130 µg of the MyoAAV2A Rep/Cap plasmid, and 130 µg of the ITR-containing SaABE8e^4cv-gRNA1^ or SaABE8e^NT^ transfer plasmid using 715 ug PEI Max® (Polysciences) in DMEM with 1% Pen-Strep. Recombinant AAV was harvested from transfected HEK293 cells and media four to five days later and purified by high-performance lipid chromatography (HPLC) purification using a POROS GoPure AAVX pre-packed column (ThermoFisher Scientific). AAV titers were quantified by a droplet digital PCR-based assay^48^ with primers and probe targeting bovine growth hormone (bGH) poly(A) signal on MyoAAV genome. Primers and probe sequences for ddPCR are listed in the key resource table.

### Satellite cell isolation

Muscle satellite cells for *in vitro* gene editing were isolated from three-week-old male *mdx^4cv^* mice as previously described^24^. For isolation of *in vivo*-edited satellite cells, triceps, pectoral, abdominal, and hindlimb muscles (excluding the TA) from MyoAAV-injected *mdx^4cv^* mice were harvested and processed for satellite cell isolation using a previously described two-step digestion protocol^49^. Briefly, pooled muscles were digested with 0.2 % collagenase type II (285 U/mg, Lifetech) in Dulbecco’s modified Eagle medium (DMEM) for 90 min at 37°C in a shaking water bath (rpm = 25). The enzyme was inactivated with 20% fetal bovine serum (FBS) in Ham’s F-10 Nutrient Mix (F10, ThermoFisher Scientific), and muscles were washed with phosphate-buffered saline (PBS) and triturated to mechanically dissociate muscle fibers from muscle tissue. These fibers were then gravitationally sedimented through a series of settling steps at 37°C for 25, 15, and 10 min, respectively. The sedimented fibers were then digested with 0.0125 % collagenase type II and 0.05 % dispase (1.81 U/mg, Lifetech) in F10 for 30 min at 37 °C in a shaking water bath to release mononuclear cells from fibers. Enzymes were inactivated by the addition of FBS, and cells were further dissociated by pipette before centrifugation and filtration through a 70 μm cell strainer. Cells were then stained with an antibody cocktail containing APC-Cy7-CD45 (Biolegend, 1:200), APC-Cy7-CD11b (Biolegend, 1:200), APC-Cy7-TER119 (Biolegend, 1:200), APC-Sca-1 (Biolegend, 1:200), PE-CD29 (Biolegend, 1:100), and Biotin-CXCR4 (BD Biosciences, 1:100) on ice for 30 minutes. After primary antibody incubation, cells were washed with staining media (Hank’s Balanced Salt Solution, HBSS, with 2% FBS) and then stained with Streptavidin-PE-Cy7 (Biolegend, 1:200) on ice for 20 minutes. After secondary antibody incubation, cells were washed twice in staining media, and resuspended in staining media with propidium iodide (PI, 1:200) to mark dead cells and Calcein Blue, AM (1:200) to mark live cells. Satellite cells were FACSorted using a FACS Aria II (BD Biosciences) based on the following cell marker profile: PI-, Calcein^+^, CD11b^-^, CD45^-^, Ter119^-^, Sca-1^-^, CD29^+^, CXCR4^+^ (Fig. S1). FACSorting was performed at the Harvard Stem Cell Institute Flow Cytometry core, and flow cytometry data were analyzed using FlowJo (BD Biosciences) analysis software.

### Satellite cell culture and transfection

Freshly isolated muscle satellite cells were cultured in growth media containing 20% horse serum (ThermoFisher Scientific), 1% Pen-Strep, and 1% Glutamax (ThermoFisher Scientific) in F10 on plates that were coated with collagen type I (1 µg/mL, Sigma Aldrich) and laminin (10 µg/mL, Invitrogen) for at least 1 hour at 37°C. Cultured satellite cells were supplemented daily with 5 ng/mL bFGF (Sigma Aldrich).

For *in vitro* gene editing, 10,000 satellite cells were FACSorted from P21 *mdx^4cv^* mice into collagen type I and laminin-coated 96-well plates containing growth media described above. One hour after FACSorting, adherent satellite cells were transfected using Lipofectamine^TM^ Stem (Invitrogen) per manufacturer’s instructions with 100 ng of SaABE8e^4cv-gRNA1^, SaABE8e^4cv-gRNA2^, SaABE8e^NT^, or CAG-EGFP reporter plasmid per well. For satellite cells transfected with the CAG-EGFP plasmid, four days after plasmid transfection, cells were detached by 2.5 mM EDTA for 10 minutes. Detached cells were resuspended in staining media and stained with PI and Calcein Blue, AM for live cell gating. EGFP+ cell frequency was analyzed using a BD Symphony A5 (BD Biosciences) to estimate the transfection rate for a specific batch of experiments (Fig. S2).

### Systemic delivery of AAV vectors *in vivo*

For systemic AAV delivery experiments, 3-day, 7-day, 3-week, 12-week, or 6-month-old male *mdx^4cv^* mice were injected with 4E13 vg/kg MyoAAV-SaABE^4cv^ or MyoAAV-SaABE^NT^ *mdx^4cv^* via retro-orbital or intraperitoneal injection. Unless otherwise specified, 4 weeks post-injection, mice were harvested.

### Extraction of genomic DNA and RNA

Fresh mouse muscle tissues were flash frozen with liquid nitrogen, and satellite cell pellets were frozen in -20℃ freezer. Genomic DNA from frozen mouse muscle tissues and satellite cell pellets were extracted using DNeasy Blood & Tissue Kit (Qiagen) per manufacturer’s instructions. Total RNA was extracted from homogenized mouse muscle tissues using a gentleMACS Octo Dissociator (Miltenyi Biotec) for 1 min 23 sec at 3170 rpm with gentleMACS M tubes. The homogenized mouse muscles tissues were then used for Trizol (Invitrogen) RNA extraction to obtain the RNA-containing aqueous phase for use in the RNeasy MiniPrep Kit (Qiagen) per manufacturer’s instructions. 0.5 µg of the extracted total RNA was digested with ezDNase (ThermoFisher Scientific) according to the manufacturer’s instructions.

DNase-treated RNA was used as template for first-strand cDNA synthesis using SuperScript™ IV First-Strand Synthesis System with oligo d(T)_20_ per manufacturer’s instructions.

### Amplicon sequencing

PCR reactions for amplicon sequencing were carried out by Q5 Hot Start polymerase (NEB) with 100 ng genomic DNA or RNA-derived cDNA from treated mouse tissues or transfected satellite cells per 25-μL reaction. Primer sequences for genomic DNA and cDNA samples are listed in the key resource table. The PCR products were purified using the QIAquick PCR Purification Kit (Qiagen). Purified PCR products were amplicon sequenced at the Massachusetts General Hospital DNA Core. FASTQ files were analyzed using the CRISPResso2 algorithm through the Docker platform. Quantification window size was set at 12, and quantification window center was set at 10. Output style was base_editor_output. A>G conversion rates for individual adenines were calculated by dividing G frequency by the sum of A and G frequencies.

### Viral genomes copy number per diploid ratio quantification

100 ng of genomic DNA extracted from TA and diaphragm was used to perform ddPCR with ddPCR Supermix for Probes (Bio-Rad) per manufacturer’s instructions. QX200 Droplet Generator was used to generate droplets, and QX200 Droplet Reader with QX Manager Software were used to determine the quantity of probe-positive droplets. Primers and probes targeting bovine growth hormone (bGH) poly(A) signal on MyoAAV genome or Glyceraldehyde-3-phosphate dehydrogenase (*Gapdh*) on mouse chromosome were used for the ddPCR reactions. For each tissue, the number of probe-positive droplets from the bGH reaction was divided by the number from the *Gapdh* reaction to obtain the viral genome copy numbers per diploid ratio. Primers and probe sequences for ddPCR are listed in the key resource table.

### Sectioning and immunofluorescence analysis

For immunofluorescence analysis, tissues were embedded in optimal cutting temperature compound (OCT, Tissue-Tek) and flash-frozen in liquid-nitrogen-cold isopentane for cryosectioning. Preserved tissues were stored at -80℃ and sectioned using CM1860 cryostat (Leica Biosciences) at a thickness of 12 μm. For Laminin and DYSTROPHIN immunostaining, muscle sections were permeabilized with 0.2% Triton X-100 in PBS for 10 minutes, followed by 3 × 5-minute DPBS washes. Sections were blocked with 10% Normal Goat Serum (NGS, Jackson ImmunoResearch) and 2% Bovine Serum Albumin (BSA, Sigma Aldrich) for 2 hours at room temperature. Sections were subsequently incubated with mouse monoclonal IgG2b anti-dystrophin primary antibody (1:100, MANDYS8, Sigma Aldrich Aldrich) and rabbit polyclonal anti-laminin primary antibody (1:200, Sigma Aldrich, L9393) in DPBS with 5% NGS and 0.01% TWEEN-20 (Sigma Aldrich) for 1 hour at room temperature, followed by 3 × 5-minute DPBS washes. Sections were then incubated with goat anti-mouse IgG2b Alexa Fluor 594 (1:1000, ThermoFisher Scientific) and goat anti-rabbit IgG Alexa Fluor 488 (1:250, Invitrogen) in DPBS with 5% NGS and 0.01% TWEEN-20 for 2 hours at room temperature, followed by 3 × 5-minute DPBS washes.

Subsequently, sections were stained with Hoechst (1:1,000) for 5 minutes, followed by 3 × 5-minute DPBS washes. All images were captured using an ECHO confocal microscope on wide-field mode (ECHO, San Diego, CA, USA) or a Zeiss 880 Inverted Microscope (ZEISS, Germany). Laminin-positive fibers were quantified using Myotally for automated quantification, followed by manual validation in ImageJ^50^. Dystrophin-positive muscle fibers were quantified manually using ImageJ in a blind analysis. The amount of dystrophin-positive muscle fibers is represented as a percentage of total laminin-positive muscle fibers.

For alpha-sarcoglycan, beta-sarcoglycan, beta-dystroglycan, and n-Nos stainings, cryosections were washed with PBS for 10 minutes, then permeabilized with 0.2% Triton X-100 in PBS for 10 minutes, followed by 3 × 5-minute PBS washes. Sections were then blocked with 5% Normal Goat Serum (NGS, Jackson ImmunoResearch), 2% Bovine Serum Albumin (BSA, Sigma Aldrich), 2% protein concentrate (M.O.M. Kit, Vector Laboratories), 1 drop/ml of M.O.M. blocking reagent (M.O.M. Kit, Vector Laboratories), and 0.1% TWEEN-20 (Sigma Aldrich) for 1 hour at room temperature, followed by 3 × 5 minute PBS washes. Sections were subsequently stained with mouse monoclonal IgG1 anti-alpha-Sarcoglycan (1:50, GeneTex), mouse monoclonal IgG1 anti-beta-Sarcoglycan (1:100, Leica), mouse monoclonal IgG2a anti-beta-Dystroglycan (1:100, Leica), or rabbit polyclonal IgG anti-n-Nos (1:1000, Immunostar) primary antibody and mouse monoclonal IgG2b anti-dystrophin primary antibody (1:100, MANDYS8, Sigma Aldrich Aldrich) in PBS with 3% Bovine Serum Albumin (BSA, Sigma Aldrich), 8% protein concentrate (M.O.M. Kit, Vector Laboratories), and 0.1% TWEEN-20 (Sigma Aldrich) at 4°C overnight, followed by 4 × 10-minute PBS washes each.

Sections were then incubated with goat anti-mouse IgG1 Alexa Fluor 488 (1:250, ThermoFisher Scientific), goat anti-rabbit IgG Alexa Fluor 647 (1:250, ThermoFisher Scientific), or goat anti-mouse IgG2a Alexa Fluor 647 (1:250, ThermoFisher Scientific), and goat anti-mouse IgG2b Alexa Fluor 594 (1:1000, ThermoFisher Scientific) secondary antibody in PBS with 3% Bovine Serum Albumin (BSA, Sigma Aldrich) and 0.1% TWEEN-20 (Sigma Aldrich) for 1 hour at room temperature, followed by 4 × 10-minute PBS washes. Sections were subsequently stained with Hoechst (1:1,000) for 5 minutes at room temperature, followed by 3 × 5-minute PBS washes.

### Western blot analysis

Protein lysates were prepared from frozen mouse tissue using a gentleMACS Octo Dissociator (Miltenyi Biotec) with 500 µL of RIPA buffer per sample (ThermoFisher Scientific) supplemented with 1% Halt protease and phosphatase inhibitor cocktail (ThermoFisher Scientific) and 1% EDTA for 53 sec at 2753 rpm with gentleMACS M tubes. Total protein concentration was quantified using the Qubit Protein Broad Range Assay Kit (ThermoFisher Scientific) following the manufacturer’s instructions.

For Western blot analysis, equal amounts of total proteins were separated on NuPAGE 3–8% Tris-acetate precast gels (ThermoFisher Scientific) using the NuPAGE electrophoresis system (Invitrogen PowerEase 350W) at 150 V for approximately 75 minutes. Proteins were transferred onto 0.2 µm nitrocellulose membranes using a Trans-Blot Turbo Transfer System (Bio-Rad) with Turbo Transfer Packs. The standard high molecular weight transfer program was modified to 15 minutes at 2.0-2.5 A constant current.

Membranes were blocked in 5% (w/v) nonfat milk (Biorad) in TBST (1× Tris-buffered saline (Bio-Rad) with 1% TWEEN-20) for 1h at room temperature (RT). Membranes were then incubated with rabbit anti-dystrophin primary antibody (Abcam; dilution 1:2,000; ab154168) diluted in 5% milk in TBST at 4°C overnight. Following primary antibody incubation, TBST-washed membranes were incubated with HRP-conjugated anti-rabbit secondary antibody (ThermoFisher Scientific) for 1 hour at room temperature. After final TBST washes, chemiluminescent detection was performed using Invitrogen SuperSignal West Femto Maximum Sensitivity Substrate (ThermoFisher Scientific). Membranes were imaged using a ChemiDoc MP Imaging System (Bio-Rad). After chemiluminescent detection, membranes were incubated with Coomassie blue staining solution (0.1% Coomassie Brilliant Blue G-250 (ThermoFisher Scientific) in 40% methanol and 10% acetic acid) for 10 minutes. After a brief water rinse, membranes were dried overnight for imaging. Band intensities were quantified using ImageJ (NIH).

### Muscle contractility measurements

P21 male WT and *mdx^4cv^* mice injected with MyoAAV-SaABE8e^4cv^ or MyoAAV-SaABE8e^NT^ were subjected to functional analysis one month after MyoAAV injection. Muscle performance was evaluated using the 300C-LR muscle lever system (Aurora Scientific Inc., Aurora, CAN) as previously described^51^. Mice were anesthetized with isoflurane (∼3% for induction and ∼2% for maintenance) and positioned on a temperature-controlled platform, with the knee immobilized and the foot secured to the motor shaft. Plantarflexor contractions were induced by percutaneous stimulation of the tibial nerve (0.2 ms pulse duration) at a current sufficient to elicit maximal isometric twitch torque. The torque–frequency relationship was assessed using 500-ms pulse trains delivered at 1, 20, 40, 50, 60, 80, 100, and 150 Hz. To evaluate susceptibility to contraction-induced injury, mice were subjected to 20 eccentric contractions at maximal isometric torque (150 ms duration, 0.2 ms pulses at 150 Hz), generated by rotating the footplate 40° backward at 800°/s following the initial 100 ms of isometric contraction. Muscle damage was quantified as the reduction in peak isometric torque.

### Statistical analysis

GraphPad Prism 10.0 software (GraphPad Software, La Jolla, CA) was used to plot data and perform unpaired T-test, one-way ANOVA, and two-way ANOVA analyses.

Illustrations were generated using BioRender.com.

### Reagents

A complete list of reagents is included in the supplemental Information.

## Data availability

Raw data are available from the corresponding author upon reasonable request.

## Acknowledgements.

We thank A. Lin for performing immunohistochemistry image quantification, N. Kheradmand at the Harvard Stem Cell and Regenerative Biology-Harvard Stem Cell Institute Flow Cytometry Core for assistance with FACS, the DNA Core at Massachusetts General Hospital Center for Computational and Integrative Biology for CRISPR sequencing services, and the members of the Wagers laboratory and HSCRB Flow Cytometry Core for input and technical support. This work was funded in part by awards and grants from Harvard University, Sarepta Therapeutics, and DP1 AG048917 (to A.J.W.)

## Author Contributions

K.H.L., A.L., and A.J.W. conceived the study and designed experiments. K.H.L, A.L., M.M., R.E., and K.M. conducted animal experiments and tissue harvests. S.v.O. and R.E. performed Western blot analyses. K.H.L. and M.M. performed protein structure prediction analyses. K.H.L., A.L., and G.K. performed cryo-sectioning and image acquisition. A.L., G.K., and R.E. performed IHC quantification. K.H.L., A.L., M.M., R.E., and K.M. performed satellite cell isolation and culture. K.H.L. and A.L. performed FACS analysis. A.L., M.M., and R.E. prepared amplicons for NGS, and K.H.L. analyzed the NGS results. K.H.L. designed and cloned the AAV transfer plasmids. K.H.L. and A.L. generated and titered AAVs. J.M. and R.K. performed *in vivo* muscle function assessments. A.J.W. supervised the study. K.H.L., A.L., and A.J.W. wrote the manuscript with input from all co-authors.

