## Supplementary Material for "*In vivo* base editing via single myotrophic adeno-associated viruses in dystrophic mouse muscle and satellite cells"

### Supplemental Information

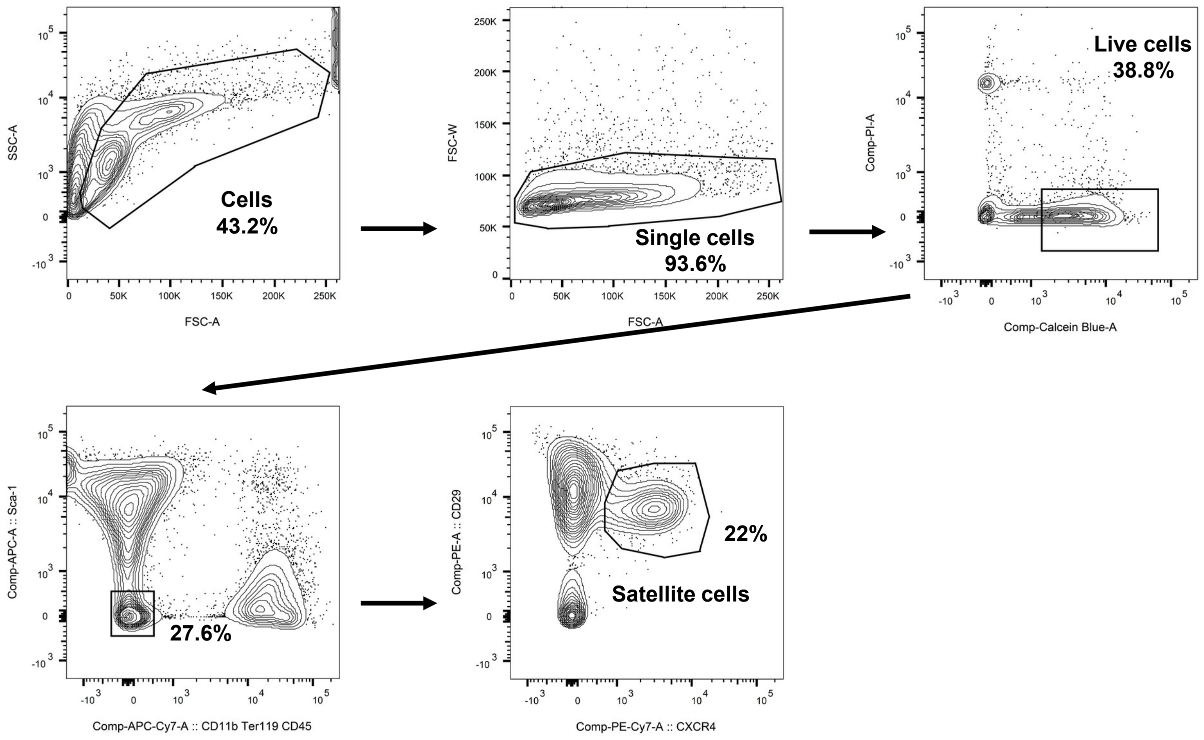


**Fig. S1 FACS isolation of muscle satellite cells, defined as Sca-1$^-$CD11b$^-$Ter119$^-$CD45$^-$CD29$^+$CXCR4$^+$ cells, from juvenile *mdx*$^{4cv}$ male mice.** (A) Muscle mononuclear cells were prepared as previously described[1]. Arrows depict sequential gating strategy for selection of Sca-1$^-$CD11b$^-$Ter119$^-$CD45$^-$CD29$^+$CXCR4$^+$ satellite cells.

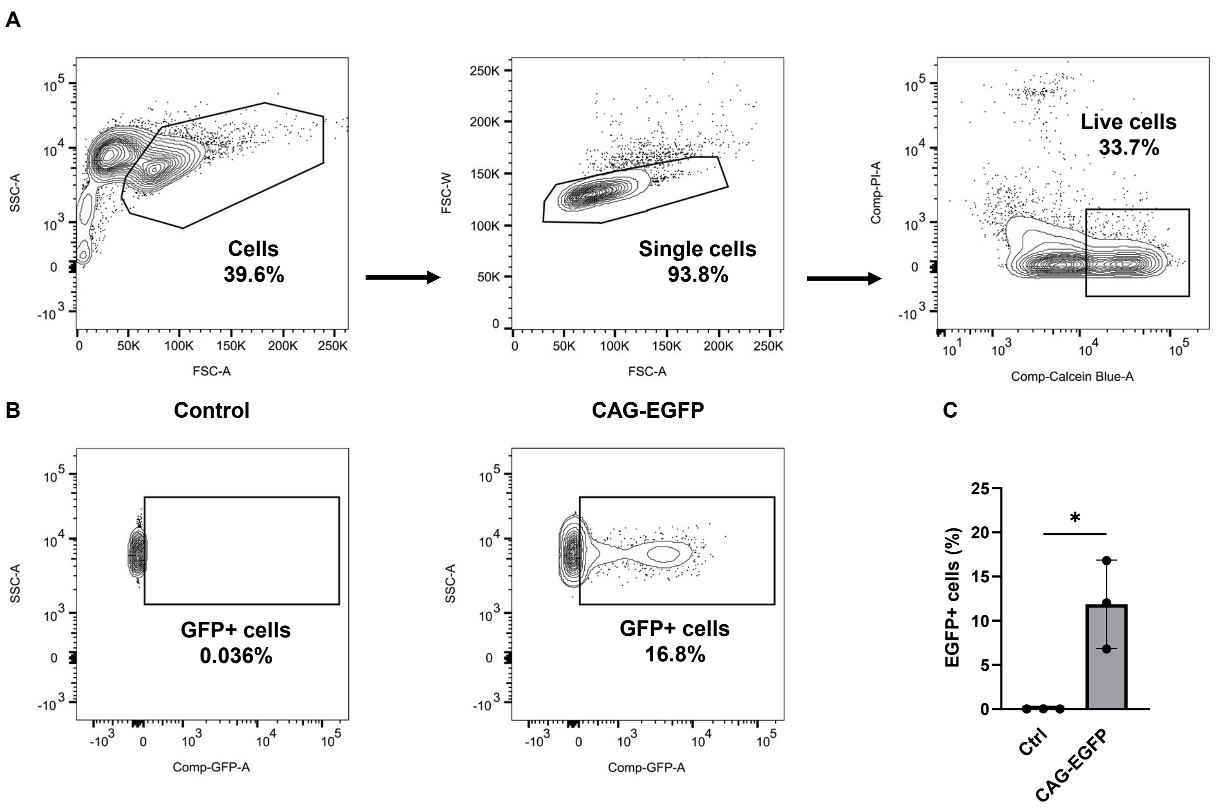


**Fig. S2.** ***In vitro*** **transfection efficiency of FACS-isolated mouse satellite cells.** (A) Gating strategy to select live cells from preparations of muscle mononuclear cells by FACS. (B) Representative flow data showing gating for EGFP+ satellite cells in control (no plasmid) and CAG-EGFP plasmid-transfected satellite cells. Cells were analyzed 4 days after transfection. (C) Quantification of %EGFP+ satellite cells in control (ctl) or CAG-EGFP transfected satellite cell cultures. n = 3 biological replicates/group; analyzed with paired T-test. Data are shown as mean ± SD. * indicates $P < 0.05$

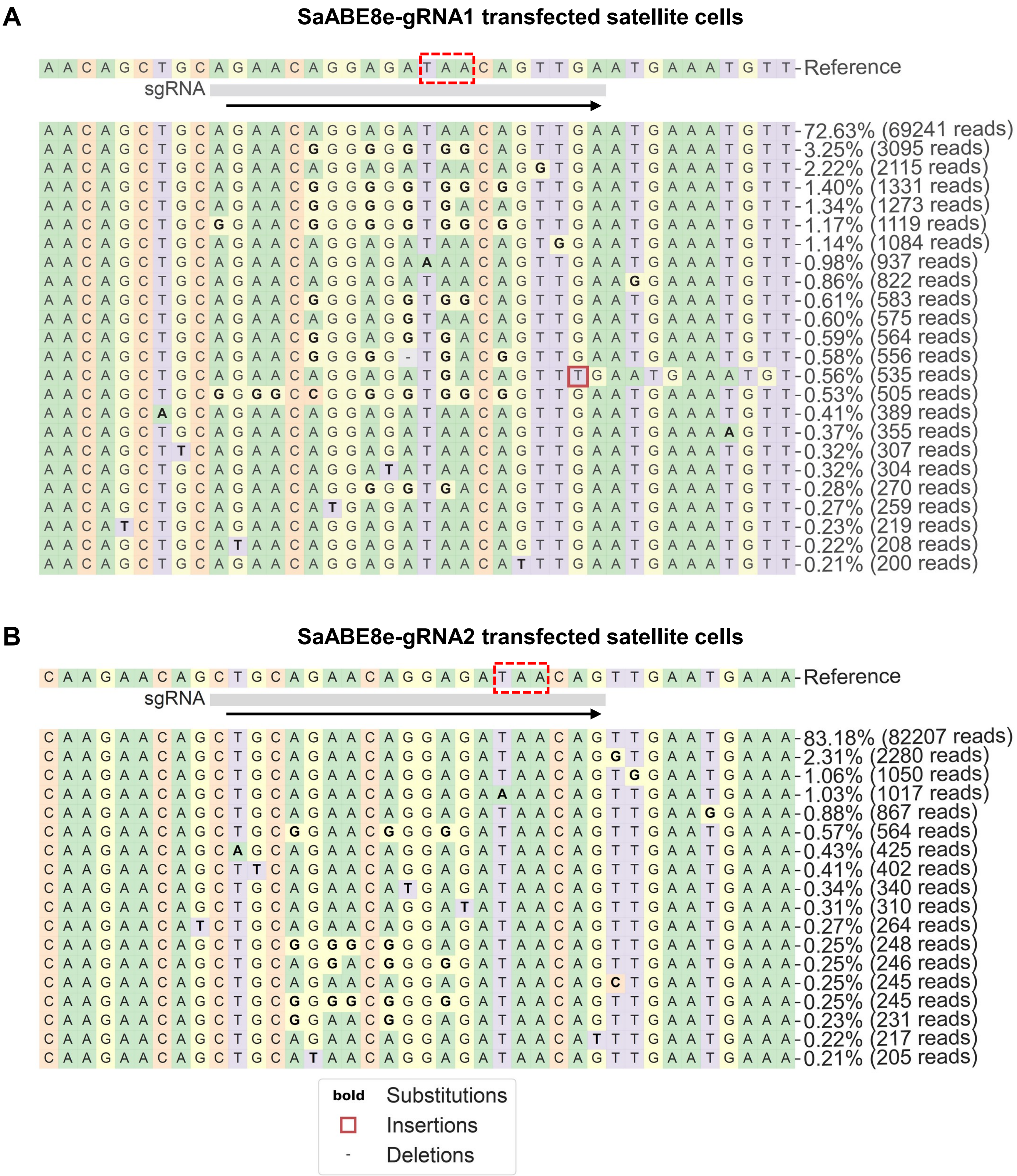


**Fig. S3 Amplicon sequencing read analysis for SaABE8e-gRNA1 and SaABE8e-gRNA2 editing outcomes focusing on residues surrounding the *mdx*$^{4cv}$ nonsense mutation and gRNA binding site.** (A,B) Sequencing results were analyzed with CRISPResso2 algorithm. Representative allele frequency tables for SaABE-gRNA1 (A) and SaABE-gRNA2-transfected (B) satellite cells are shown. Dashed red box indicates the *mdx*$^{4cv}$ nonsense mutation, and arrow indicates the orientation of gRNA binding.

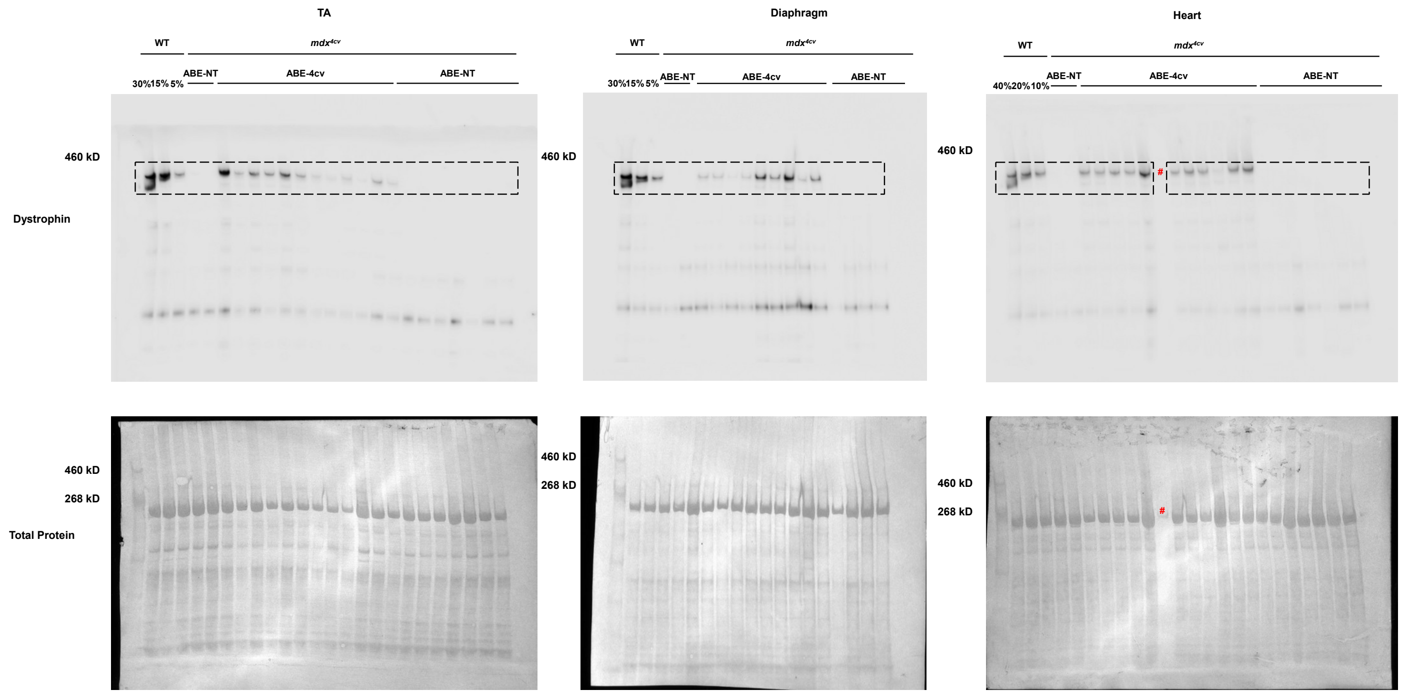


**Fig. S4. Systemic delivery of MyoAAV-SaABE8e$^{4cv}$ restores dystrophin expression in *mdx$^{4cv}$* mice.** Juvenile (P21) male *mdx$^{4cv}$* mice were injected retro-orbitally injected with 4E13 VG/kg MyoAAV-SaABE8e$^{4cv}$ or MyoAAV-SaABE8e$^{NT}$, and tissues harvested one month later. Full membrane images of dystrophin protein expression detected by Western Blot of protein lysate from TA (left), diaphragm (middle), or heart (right) of the indicated wild-type (WT) mice, or *mdx$^{4cv}$* mice injected with MyoAAV-SaABE8e$^{NT}$ (NT), or MyoAAV-SaABE8e$^{4cv}$ (4cv-gRNA1). To estimate the efficiency of dystrophin protein rescue, the first 3 lanes were loaded with different percentages (30% - 15% - or 5% for TA and diaphragm, and 40% - 20% - or 10% for heart) of WT lysate. Coomassie blue staining of each membrane is shown below each blot, demonstrating equivalent protein loading. Dashed squares highlight the regions of each blot used for quantification of dystrophin protein (427 kDa). # indicates a lane in which a technical error prevented successful protein loading; data from this lane were not included in the analysis.

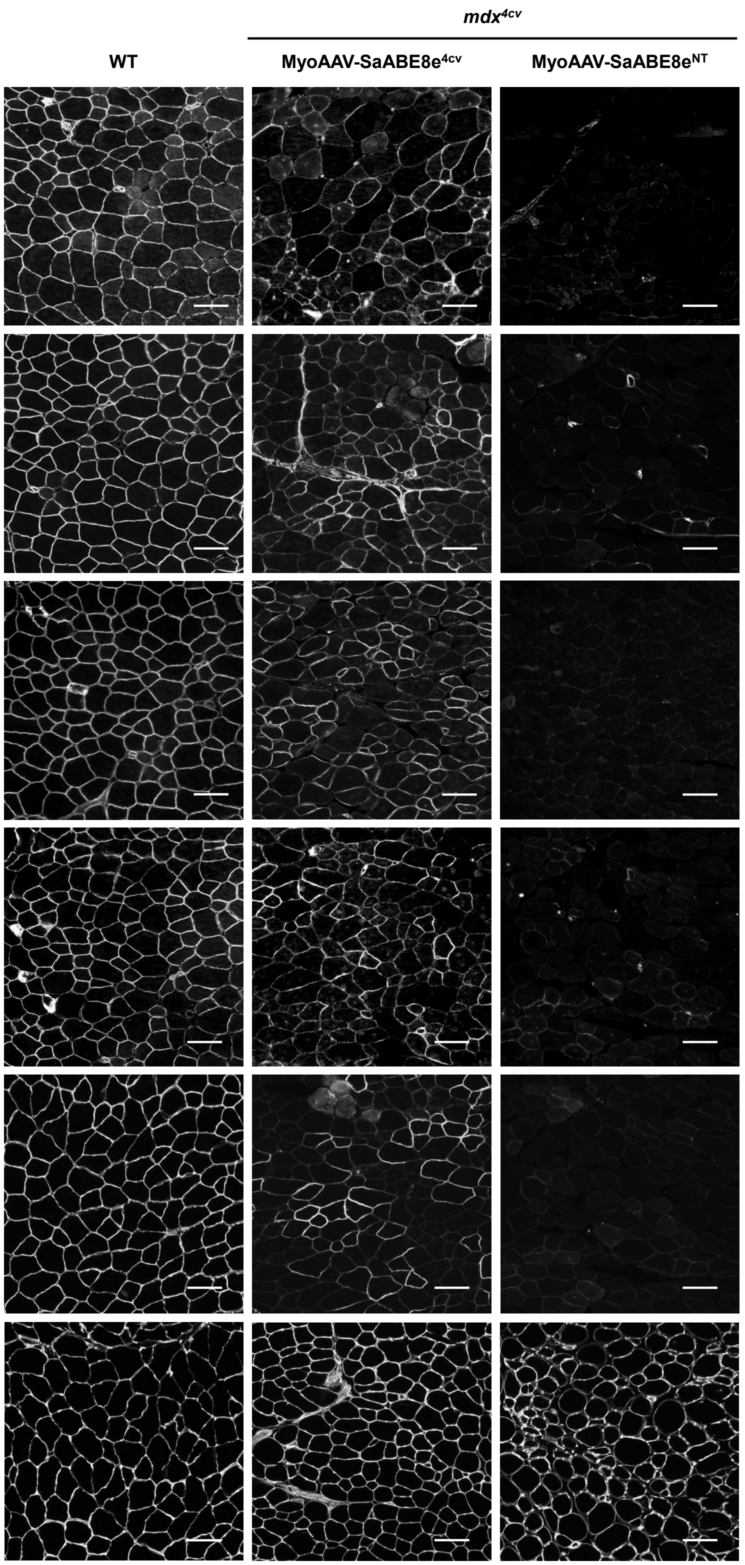


**Fig. S5 Restoration of dystrophin-associated protein complex (DAPC) and sarcolemmal nNOS in MyoAAV-SaABE8e$^{4cv}$ injected muscles.** Immunofluorescence staining for the indicated dystrophin-associated protein complex (DAPC) components or neuronal nitric oxide synthase (nNos) in muscle sections from wild-type mice (WT, left) or *mdx*$^{4cv}$ mice injected retro-orbitally with MyoAAV-SaABE8e$^{4cv}$ (middle) or MyoAAV-SaABE8e$^{NT}$ (right). Scale bar = 100 µm.

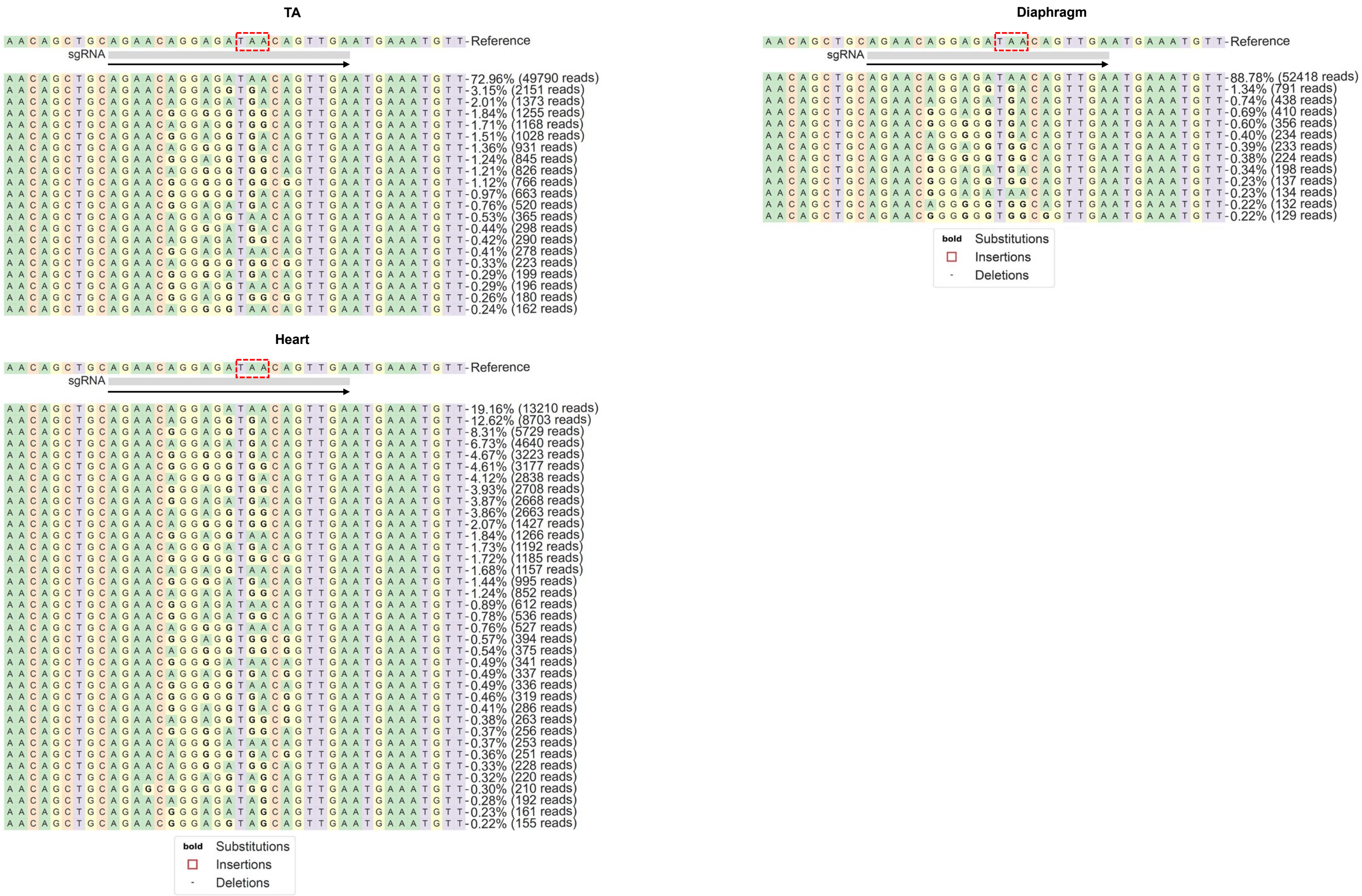


**Fig. S6. Amplicon sequencing read analysis at the site of the mdx$^{4cv}$ nonsense mutation for the indicated tissues from MyoAAV-SaABE8e$^{4cv}$ injected *mdx$^{4cv}$* mice.** Mice were retro-orbitally injected at P21 with 4E13 VG/kg MyoAAV-SaABE8e$^{4cv}$ and TA, diaphragm, and heart were harvested one month later for amplicon sequencing. cDNA amplicon sequencing results were analyzed with CRISPREsso2 algorithm. Representative allele frequency tables for each sample type are shown. Dashed red box indicates the *mdx$^{4cv}$* nonsense mutation site, and arrow indicates the direction of the gRNAs.

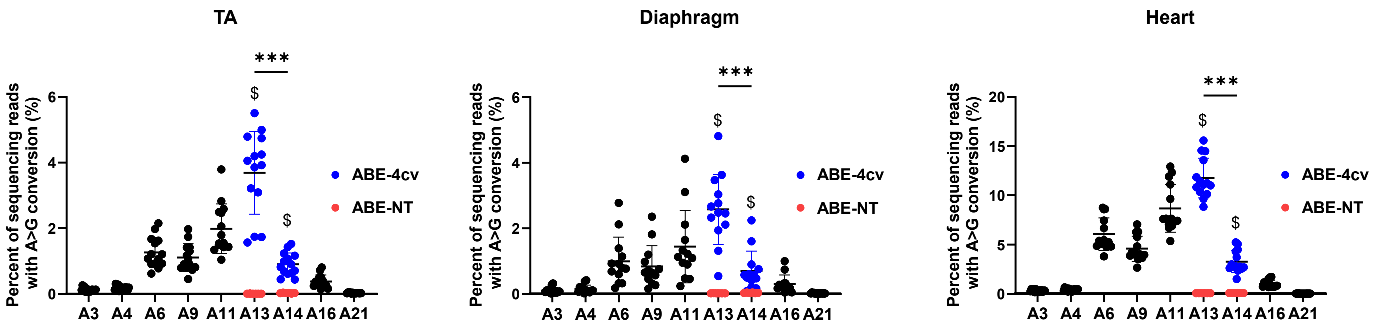


**Fig. S7. Amplicon sequencing read analysis of genomic DNA from TA, Diaphragm, and heart of male *mdx*$^{4cv}$ mice injected retro-orbitally with MyoAAV-SaABE8e$^{4cv}$ at P21.** Mice received 4E13 VG/kg MyoAAV-SaABE8e$^{4cv}$ and TA, diaphragm, and heart were harvested one month later. Genomic DNA from TA, diaphragm, and heart were used for amplicon sequencing at the *mdx*$^{4cv}$ mutation site. A>G conversion rates for individual adenines (A) were calculated from sequencing reads. Adenine conversions in the *mdx*$^{4cv}$ premature stop codon of MyoAAV-SaABE8e$^{4cv}$-injected mice are shown in dark blue (A13 and A14). A13 and A14 A>G conversion rates from MyoAAV-SaABE8e$^{NT}$-injected *mdx*$^{4cv}$ mice are shown in red. $ indicates a significant difference between MyoAAV-SaABE8e$^{4cv}$- and MyoAAV-SaABE8e$^{NT}$-injected *mdx*$^{4cv}$ mice in A13 or A14 A>G conversion rates (determined by unpaired T-test). n = 13-14 biological replicates/group; analyzed with one-way repeated measures ANOVA. Data are shown as mean ± SD. *** indicates $P < 0.001$.

**A**

| Sequence | PAM | Off-target Score | Gene | Chromosome | Strand | Position | Mismatches |
|---|---|---|---|---|---|---|---|
| CTTACAGGAGATAACAGTTGT | GTGAA | 0.595238095 | | chr15 | + | 48935095 | 4 |
| CTAACAGGAAATAACAGTTTA | AAGGA | 0.561465721 | | chr1 | - | 95644468 | 4 |
| GGAAAAGGAGGTAACATTTGA | GAGAG | 0.418871252 | | chr12 | + | 54182057 | 4 |
| TGAACATGAGATAACAGAGGA | TGGGA | 0.418871252 | | chr1 | - | 8975204 | 4 |
| TGAAAAGGAGAAAACTGTTGA | AAGAG | 0.401861252 | | chr13 | - | 91402201 | 4 |
| AGAAAAGCAGATACCAGTTGA | CAGAA | 0.371674491 | | chr9 | - | 33164855 | 3 |
| GGAACAGGAGAAAACAGTGGT | GAGAG | 0.371674491 | | chr7 | - | 113202558 | 4 |
| TGAACATAAGATAATAGTTGA | ATGAA | 0.358220211 | | chr5 | - | 88541343 | 4 |
| AGAACAGCAGATATGAGTTGA | CTGAG | 0.345705968 | | JH584298.1 | - | 24533 | 3 |
| AGAACAGCAGATATGAGTTGA | CTGAG | 0.345705968 | | GL456354.1 | + | 119105 | 3 |

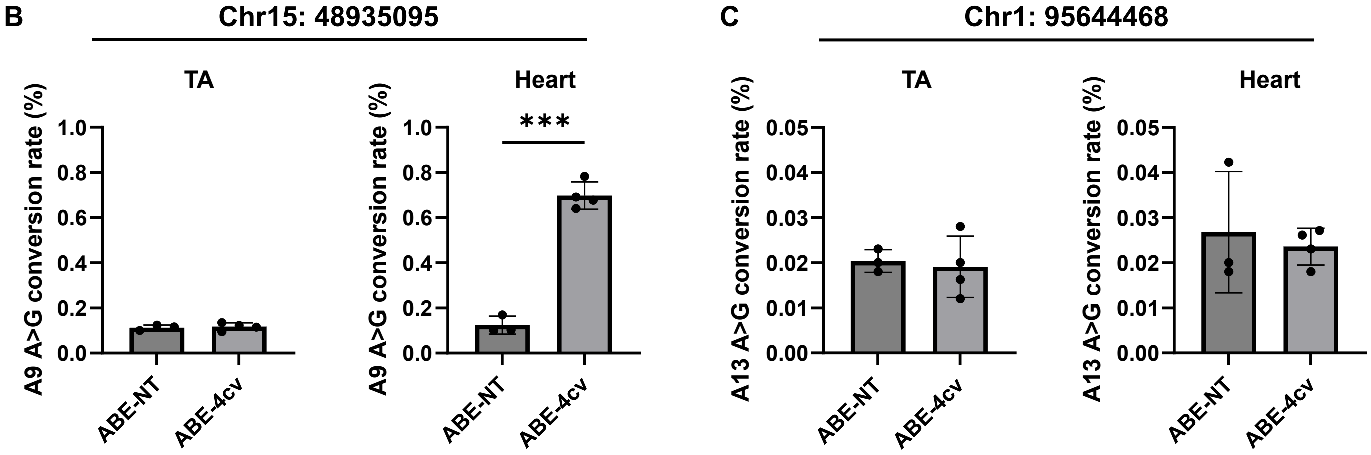


**Fig. S8. MyoAAV-SaABE8e$^{4cv-gRNA1}$ exhibits minimal activity at *in silico* predicted off-target sites, none of which are in gene coding regions.** (A) Top ten *in silico* predicted off-targets with their highest predicted off-target score, targeting site sequence, PAM for gRNA1, gene encoded at this site, chromosomal location, positive or negative strand orientation, position at the indicated chromosome, and number of mismatches between the off-target sites and gRNA1. (B-C) Juvenile (P21) male *mdx*$^{4cv}$ mice were injected with either MyoAAV-SaABE8e$^{4cv}$ or MyoAAV-SaABE8e$^{NT}$ (4E13VG/kg). Genomic DNA from TA muscle and heart was extracted one month later for amplicon sequencing at predicted off-target sites at Chr15: 48935095 (B) and Chr1: 95644468 (C). (B) A>G conversion rate at the most efficiently converted position (A9) at the off-target site on Chr15: 48935095 was used to quantify ABE activity in TA muscle and heart. (C) A13 A>G conversion rate at the off-target site on Chr1: 95644468 was used to quantify ABE activity in TA muscle and heart (no ABE activity was detected at this off-target site). n = 3-4 biological replicates/group; unpaired T-test. Data are shown as mean ± SD. *** indicates $P < 0.001$.

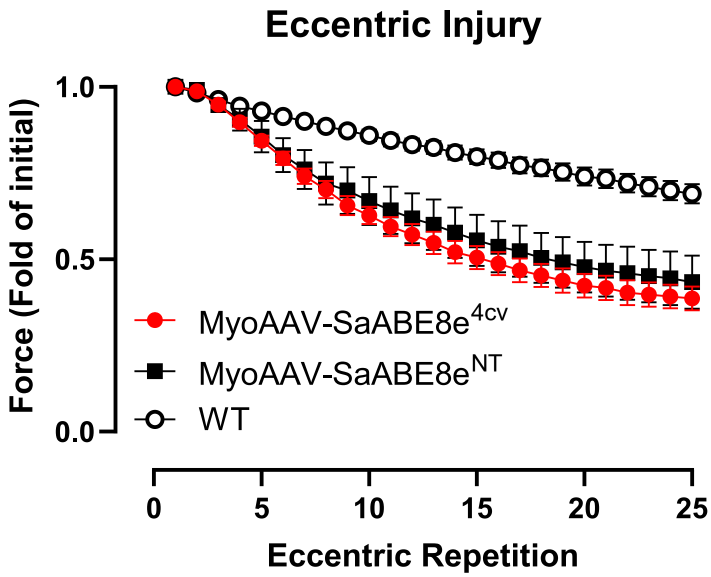


**Fig. S9. Systemic delivery of MyoAAV-SaABE8e$^{4cv\text{-}gRNA1}$ does not rescue susceptibility to repetitive eccentric contraction-induced injury at four weeks after gene editing.** Juvenile (P21) male *mdx*$^{4cv}$ mice were retro-orbitally injected with 4E13 VG/kg MyoAAV-SaABE8e$^{4cv}$ or MyoAAV-SaABE8e$^{NT}$. One month post-injection, male wildtype (WT), MyoAAV-SaABE8e$^{4cv}$-injected *mdx*$^{4cv}$, and MyoAAV-SaABE8e$^{NT}$-injected *mdx*$^{4cv}$ mice were assessed for *in vivo* gastrocnemius muscle force output. The force drop of gastrocnemius muscles from the indicated mice after the indicated number of experimentally induced eccentric contractions was measured. n = 6-12 biological replicates/group. Data analyzed with two-way repeated measures ANOVA showed no statistical difference between mice receiving MyoAAV-SaABE8e$^{4cv}$ and mice receiving MyoAAV-SaABE8e$^{NT}$. Data plotted as mean ± SD.

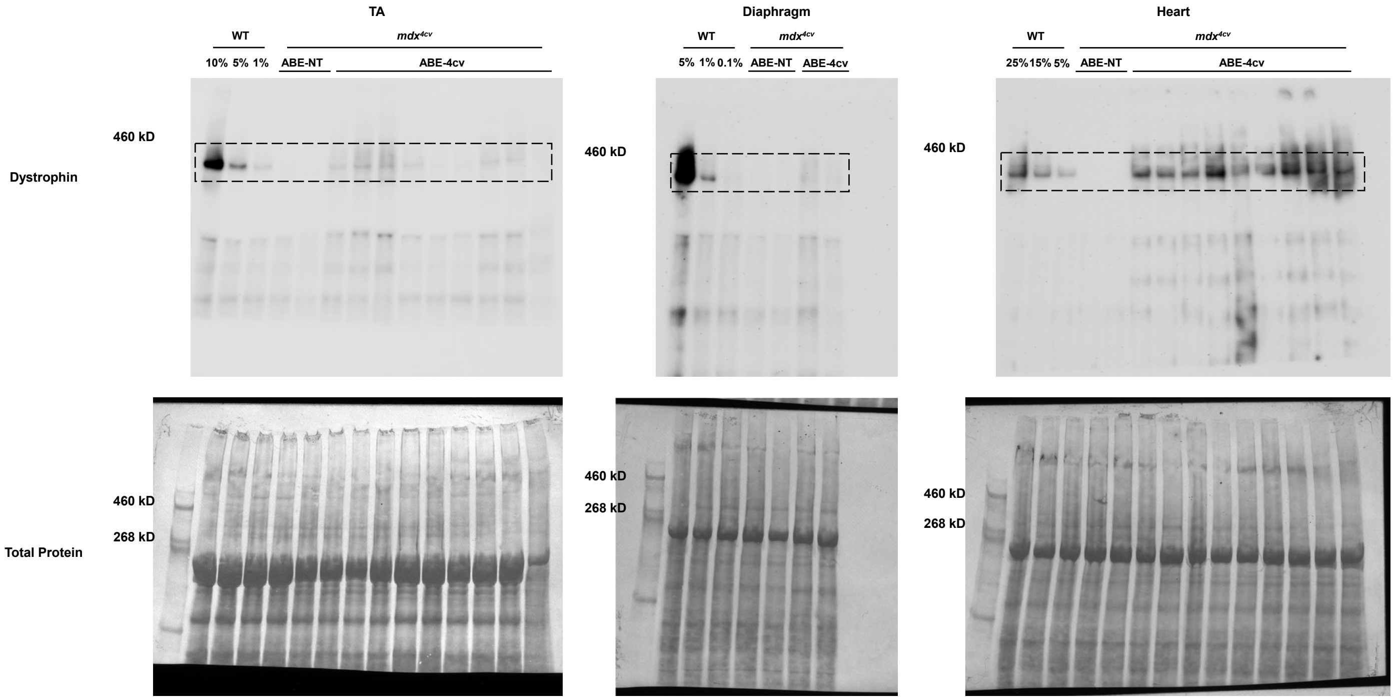


**Fig. S10. Systemic delivery of MyoAAV-SaABE8e$^{4cv}$ in *mdx*$^{4cv}$ mice sustains production of edited Dystrophin protein throughout adulthood.** Juvenile (P21) male *mdx*$^{4cv}$ mice were injected retro-orbitally with 4E13 VG/kg MyoAAV-SaABE8e$^{4cv}$ or MyoAAV-SaABE8e$^{NT}$, and tissues harvested six months later. Full membrane images of dystrophin protein expression detected by Western Blot of protein lysate from TA (left), diaphragm (middle), or heart (right) of the indicated wild-type (WT) mice, or *mdx*$^{4cv}$ mice injected with MyoAAV-SaABE8e$^{NT}$ (NT), or MyoAAV-SaABE8e$^{4cv}$ (4cv-RNA1). To estimate the efficiency of dystrophin protein rescue, the first 3 lanes were loaded with different percentages (10% - 5% - or 1% for TA, 5% - 1% - or 0.1% for diaphragm, and 25% - 15% - or 5% for heart) of WT lysate. Coomassie blue staining of each membrane is shown below each blot, demonstrating equivalent protein loading. Dashed squares highlight the regions of each blot used for quantification of dystrophin protein (427 kDa).

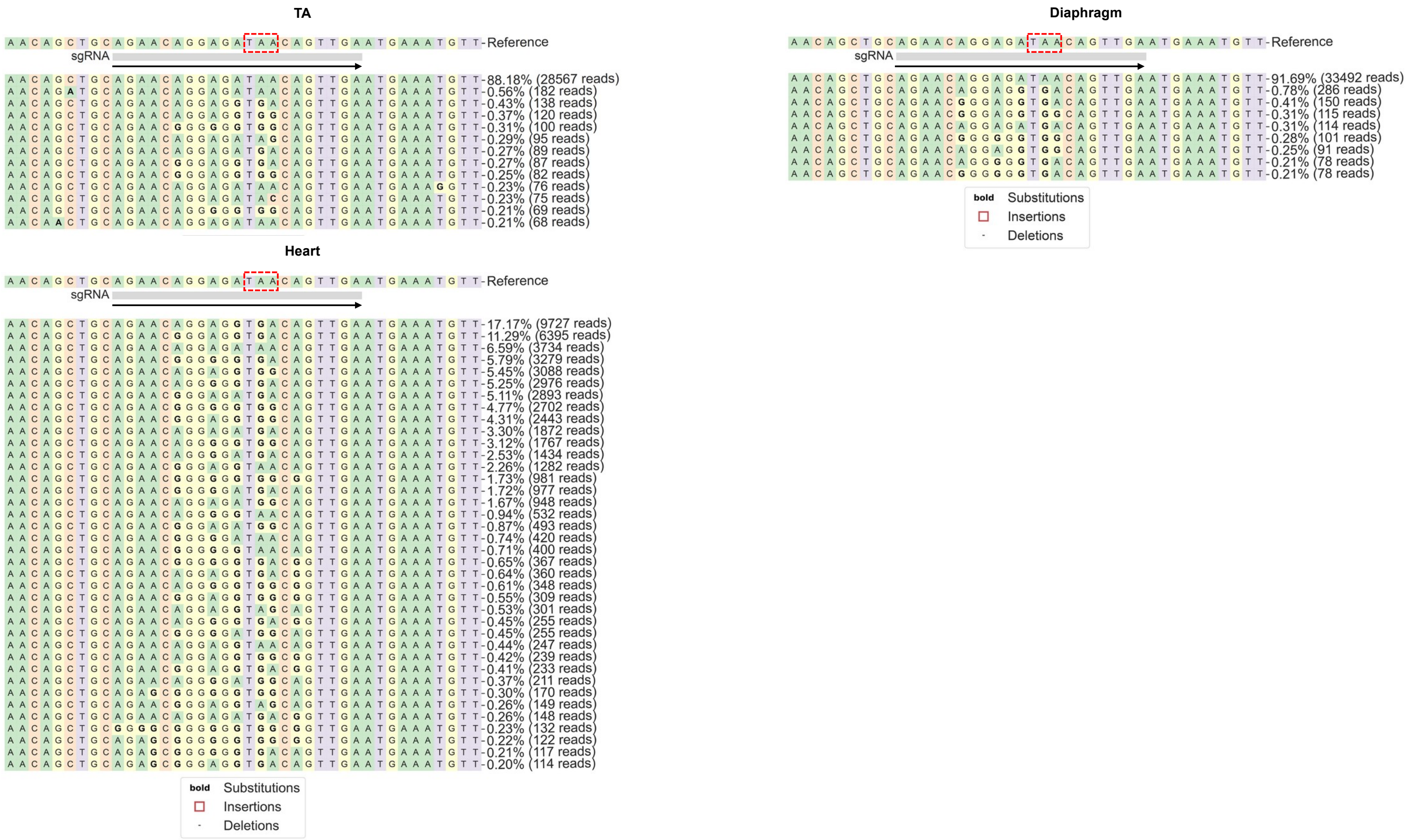


**Fig. S11. Amplicon sequencing read analysis at the site of the *mdx*$^{4cv}$ nonsense mutation for the indicated tissues from *mdx*$^{4cv}$ mice injected with MyoAAV-SaABE8e$^{4cv}$.** Mice were retro-orbitally injected at P21 with 4E13 VG/kg MyoAAV-SaABE8e$^{4cv}$ and TA, diaphragm, and heart muscle harvested six months later. cDNA amplicon sequencing results were analyzed with CRISPResso2 algorithm. Representative allele frequency tables for each sample type are shown. Dashed red box indicates the *mdx*$^{4cv}$ nonsense mutation site, and arrow indicates the direction of the gRNAs.

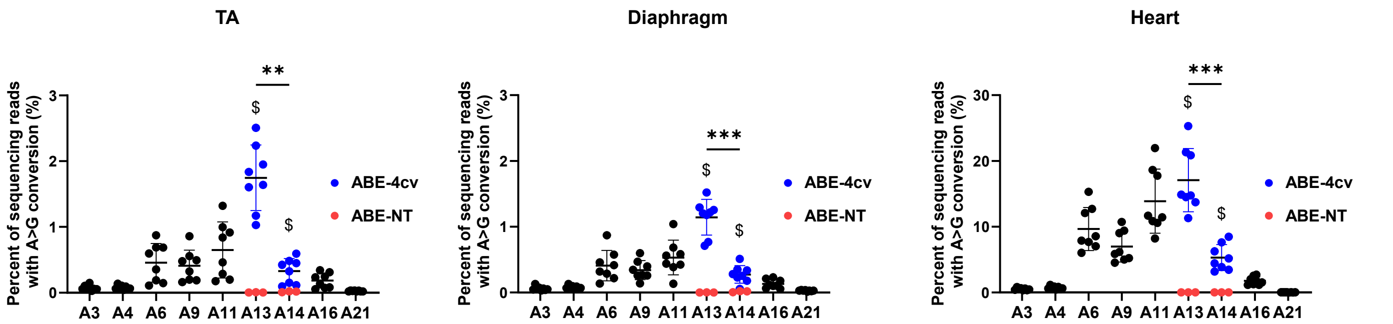


**Fig. S12. Amplicon sequencing read analysis of genomic DNA from TA, diaphragm, and heart of male *mdx*$^{4cv}$ mice injected retro-orbitally with MyoAAV-SaABE8e$^{4cv}$ at P21.** Mice were administered 4E13 VG/kg MyoAAV-SaABE8e$^{4cv}$ and TA, diaphragm, and heart harvested six months later. Genomic DNA from TA, diaphragm, and heart were used for amplicon sequencing at the *mdx*$^{4cv}$ mutation site. A>G conversion rates for individual adenines (A) of MyoAAV-SaABE8e$^{4cv}$-injected *mdx*$^{4cv}$ mice were calculated from sequencing reads. Adenine conversions in the *mdx*$^{4cv}$ premature stop codon from MyoAAV-SaABE8e$^{4cv}$-injected mice are shown in dark blue (A13 and A14). A13 and A14 A>G conversion rates from MyoAAV-SaABE8e$^{NT}$-injected *mdx*$^{4cv}$ mice are shown in red. $ indicates a significant difference between MyoAAV-SaABE8e$^{4cv}$- and MyoAAV-SaABE8e$^{NT}$-injected *mdx*$^{4cv}$ mice in A13 or A14 A>G conversion rates (determined by unpaired T-test). n = 8 biological replicates/group; analyzed with one-way repeated measures ANOVA. Data are shown as mean ± SD. ** and *** indicate P < 0.01 and < 0.001, respectively.

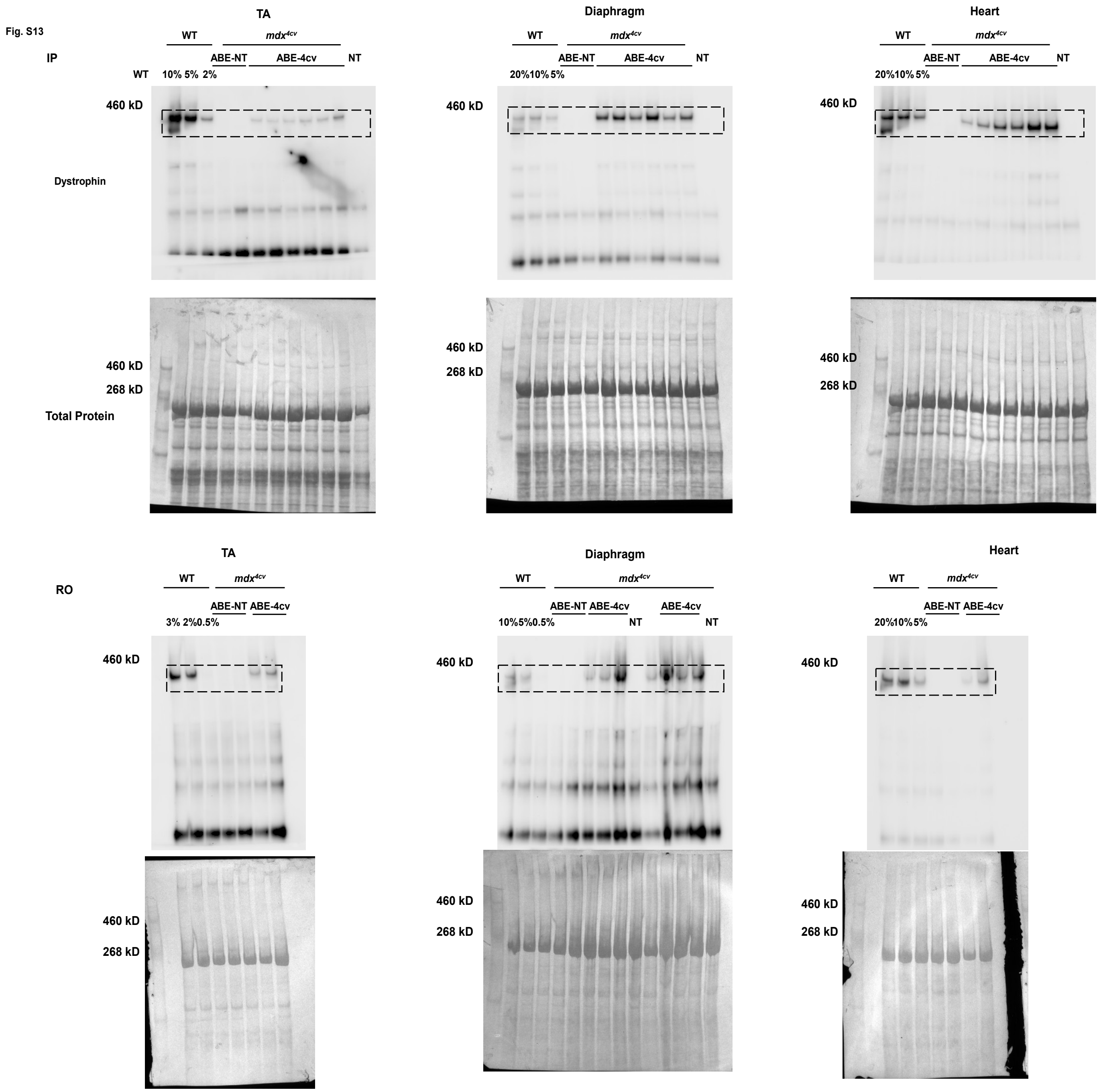


**Fig. S13. Inefficient rescue of dystrophin deficiency following systemic delivery of MyoAAV-SaABE8e$^{4cv}$ in neonatal *mdx$^{4cv}$* mice.** Neonatal (P3) male *mdx$^{4cv}$* mice were injected intraperitoneally or retro-orbitally with 4E13 VG/kg MyoAAV-SaABE8e$^{4cv}$ or MyoAAV-SaABE8e$^{NT}$, and tissues harvested one month later. Full membrane images of dystrophin protein expression detected by Western Blot of protein lysate from TA (left), diaphragm (middle), or heart (right) of the indicated wild-type (WT) mice, or mdx$^{4cv}$ mice injected with MyoAAV-SaABE8e$^{NT}$ (NT), or MyoAAV-SaABE8e$^{4cv}$ (4cv-gRNA1). To estimate the efficiency of dystrophin protein rescue, the first 3 lanes were loaded with different percentages (%) of WT lysate, as indicated. Coomassie blue staining of each membrane is shown below each blot, demonstrating equivalent protein loading. Dashed squares highlight the regions of each blot used for quantification of dystrophin protein (427 kDa).

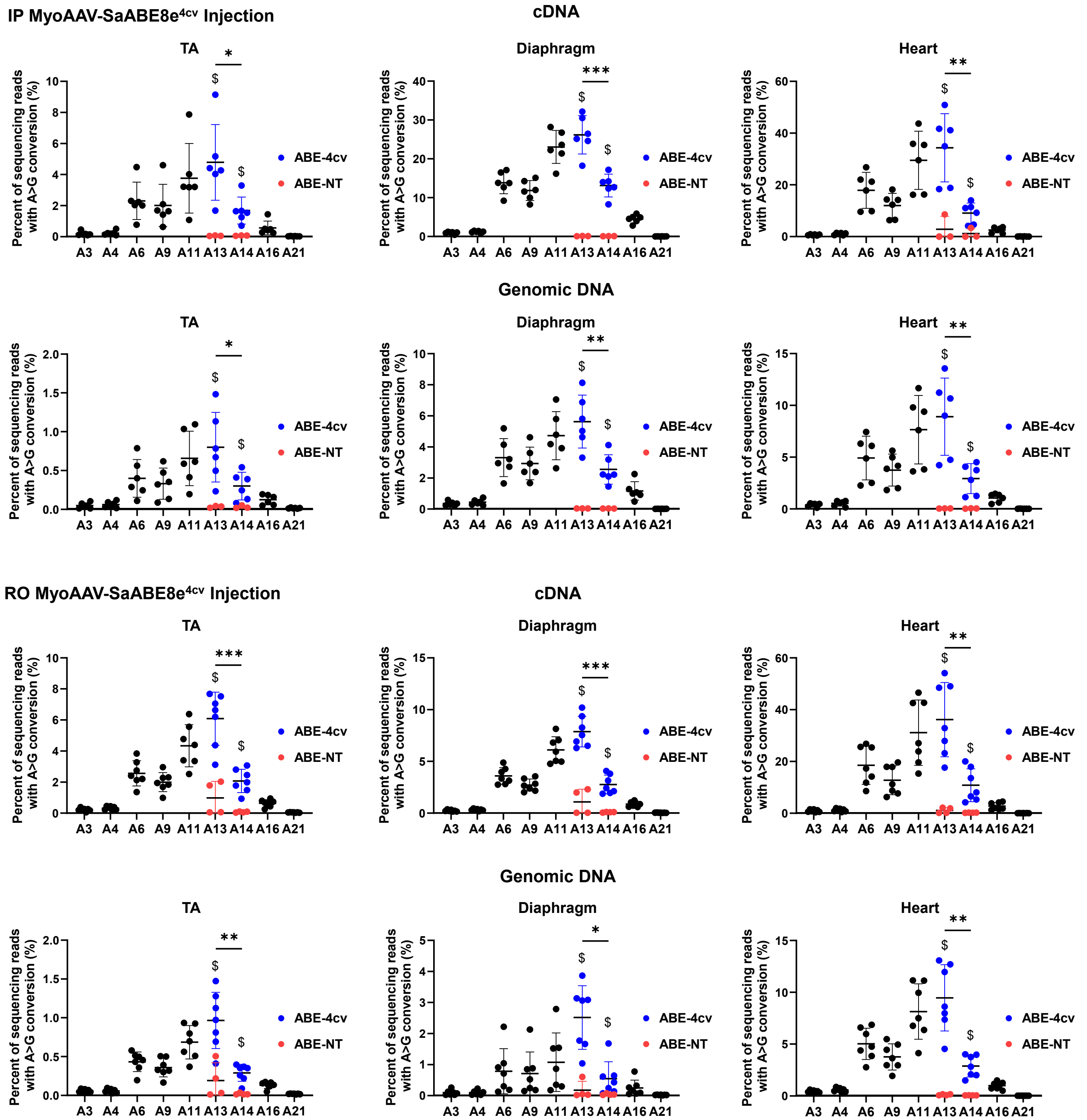


**Fig. S14. Amplicon sequencing read analysis of cDNA and genomic DNA from TA, Diaphragm, and heart of male *mdx*$^{4cv}$ mice injected retro-orbitally with MyoAAV-SaABE8e$^{4cv}$ or MyoAAV-SaABE8e$^{NT}$ at P3.** Mice were injected intraperitoneally (IP) or retro-orbitally (RO) at P3 with 4E13 VG/kg MyoAAV-SaABE8e$^{4cv}$ and TA, diaphragm, and heart harvested one month later. cDNA and genomic DNA from TA, diaphragm, and heart were used for amplicon sequencing at the *mdx*$^{4cv}$ mutation site. A>G conversion rates for individual adenines (A) of MyoAAV-SaABE8e$^{4cv}$-injected *mdx*$^{4cv}$ mice were calculated from sequencing reads. Adenine conversions in the *mdx*$^{4cv}$ premature stop codon from MyoAAV-SaABE8e$^{4cv}$-injected mice are shown in dark blue (A13 and A14). A13 and A14 A>G conversion rates from MyoAAV-SaABE8e$^{NT}$-injected *mdx*$^{4cv}$ mice are shown in red. $ indicates a significant difference between MyoAAV-SaABE8e$^{4cv}$- and MyoAAV-SaABE8e$^{NT}$-injected *mdx*$^{4cv}$ mice in A13 or A14 A>G conversion rates (determined by unpaired T-test). n = 6-7 biological replicates/group; analyzed with one-way repeated measures ANOVA. Data are shown as mean ± SD. *, **, and *** indicate $P < 0.05$, $< 0.01$, and $<0.001$, respectively.

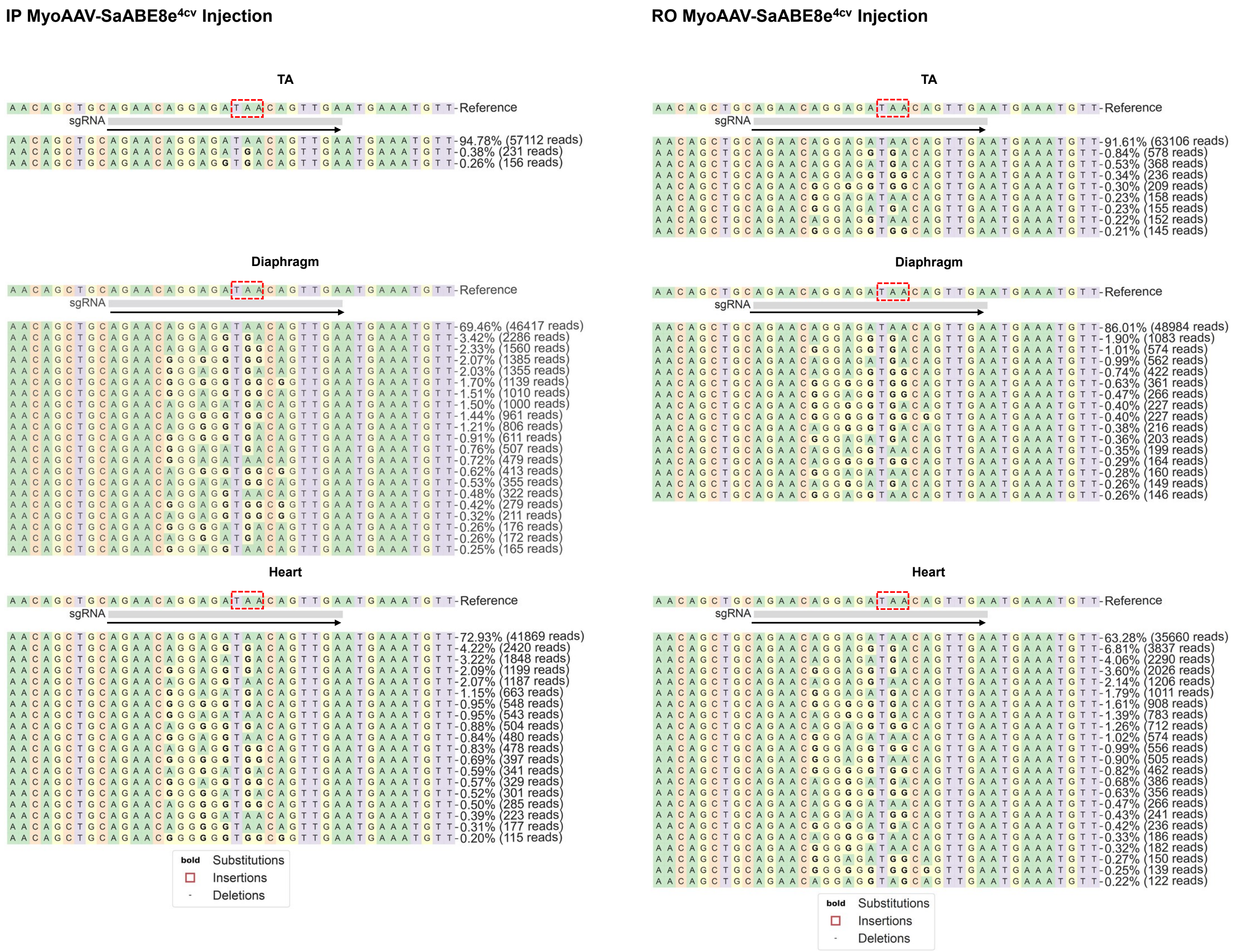


**Fig. S15. Amplicon sequencing read analysis at the site of the mdx$^{4cv}$ nonsense mutation for the indicated tissues from *mdx$^{4cv}$* mice injected with MyoAAV-SaABE8e$^{4cv}$ at P3.** Mice were injected either intraperitoneally or retro-orbitally at P3 with 4E13 VG/kg MyoAAV-SaABE8e$^{4cv}$ and TA, diaphragm, and heart harvested one month later. cDNA amplicon sequencing results were analyzed with CRISPResso2 algorithm. Representative allele frequency tables for each sample type are shown. Dashed red box indicates the *mdx$^{4cv}$* nonsense mutation site, and arrow indicates the direction of the gRNAs.

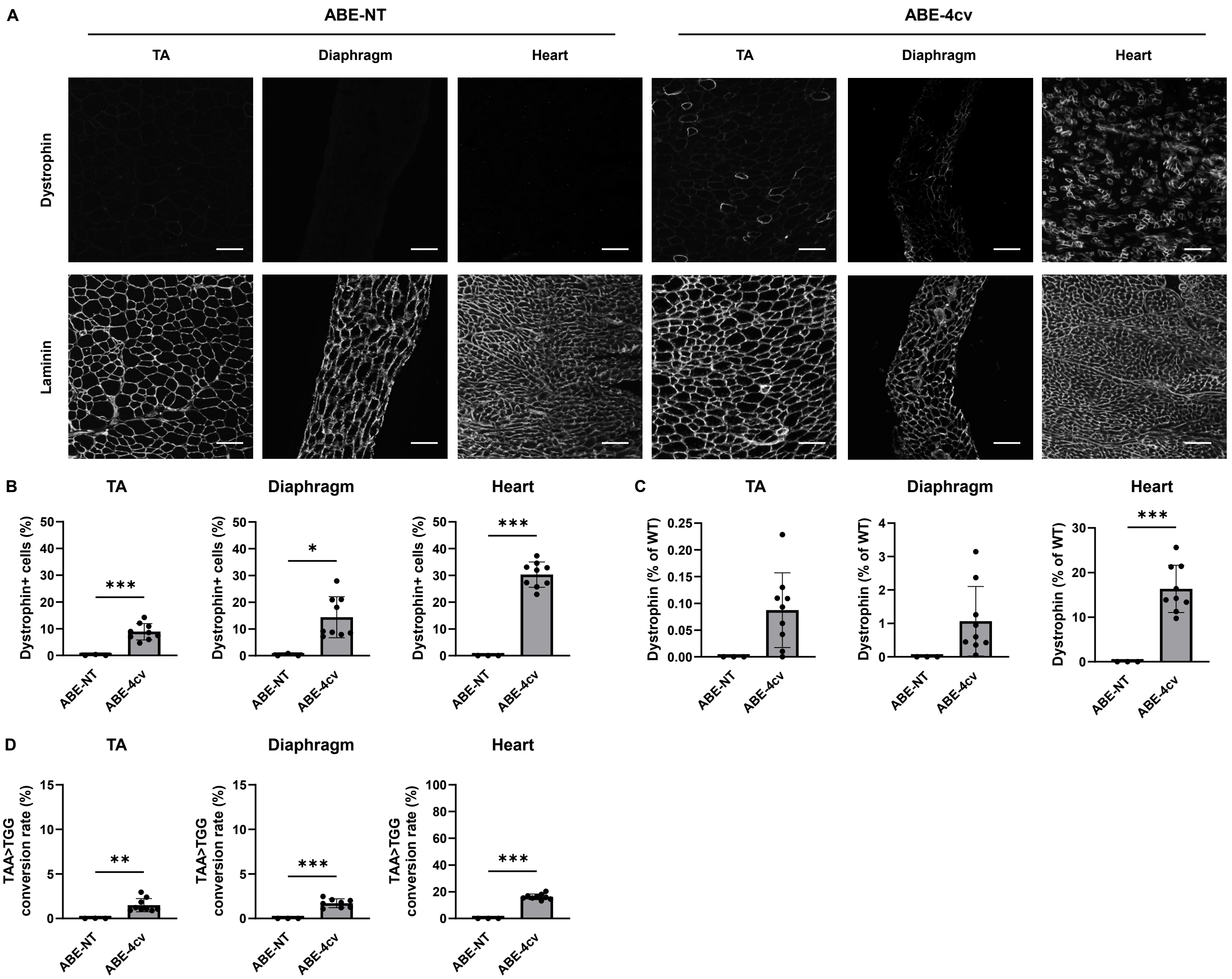


**Fig. S16. Systemic delivery of MyoAAV-SaABE8e$^{4cv}$ in P7 *mdx*$^{4cv}$ mice efficiently rescues dystrophin deficiency in heart.** P7 male *mdx*$^{4cv}$ mice were retro-orbitally injected with 4E13 VG/kg MyoAAV-SaABE8e$^{4cv}$ or MyoAAV-SaABE8e$^{NT}$, and tissues harvested one month later. (A) Representative single-channel immunofluorescence images from ABE-NT or ABE-4cv injected mice showing the same field of tibialis anterior (TA), diaphragm, or heart stained for dystrophin (top) or laminin (bottom), as indicated. Scale bar = 100 μm. (B) Quantification of dystrophin-positive myofibers or cardiomyocytes, calculated as percent (%) of total cells (derived from laminin staining). (C) Semi-quantitative analysis of dystrophin protein expression in TA, diaphragm, and heart of mice injected with MyoAAV-SaABE8e$^{NT}$ or MyoAAV-SaABE8e$^{4cv}$. Representative blots are shown in Fig. S17. (D) cDNA from TA, diaphragm, and heart were used for amplicon sequencing at the *mdx*$^{4cv}$ mutation site. The A>G conversion rates of individual adenines are shown in Fig. S18, and representative allele frequency tables are shown in Fig. S19. Percentages of reads from cDNA with *mdx*$^{4cv}$ nonsense mutation converted to non-stop (missense) mutation (TAA>TGG) are shown. n = 3-9 biological replicates/group; B-D are analyzed with unpaired T-test. Data are shown as mean ± SD. *, **, and *** indicate $P < 0.05$, $< 0.01$, and $<0.001$, respectively.

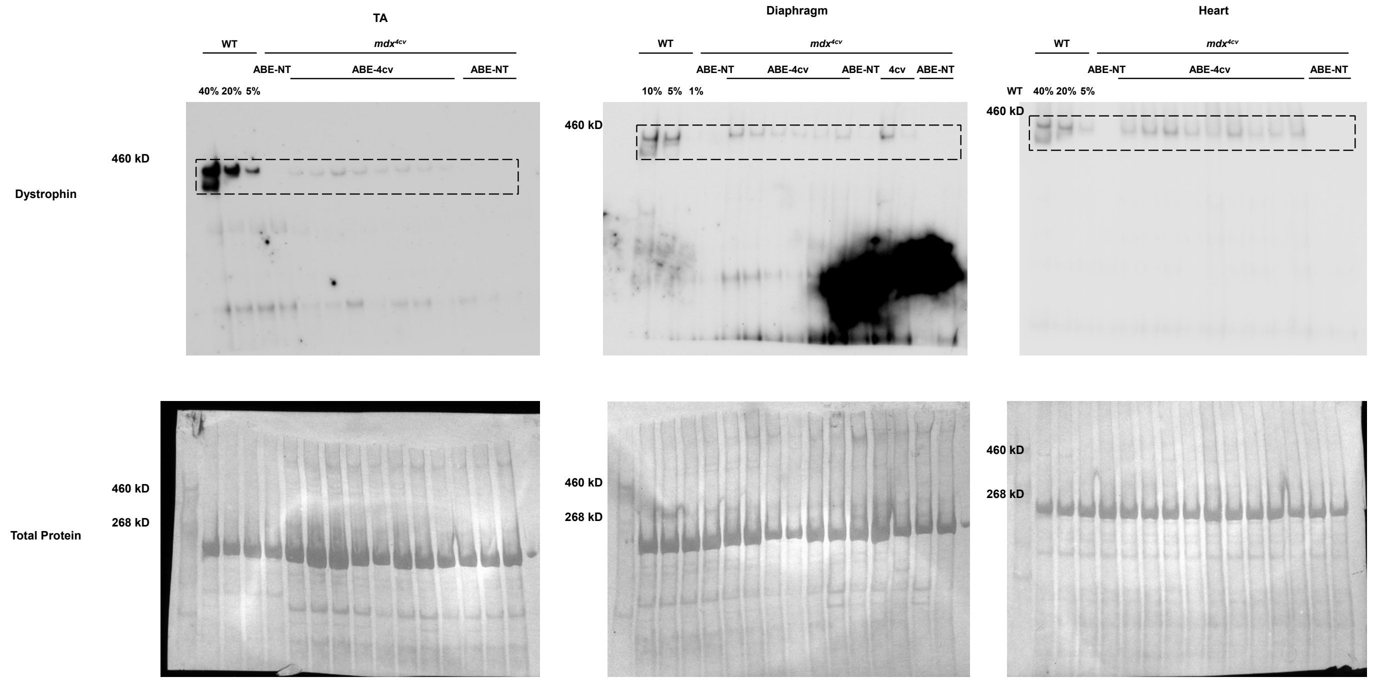


**Fig. S17. Systemic delivery of MyoAAV-SaABE8e$^{4cv}$ in P7 *mdx$^{4cv}$* mice rescues dystrophin deficiency in heart.** Neonatal (P7) male *mdx$^{4cv}$* mice were injected retro-orbitally with 4E13 VG/kg MyoAAV-SaABE8e$^{4cv}$ or MyoAAV-SaABE8e$^{NT}$, and tissues harvested one month later. Full membrane images of dystrophin protein expression detected by Western Blot of protein lysate from TA (left), diaphragm (middle), or heart (right) from wild-type (WT) mice, or *mdx$^{4cv}$* mice injected with MyoAAV-SaABE8e$^{NT}$ (NT), or MyoAAV-SaABE8e$^{4cv}$ (4cv-RNA1), as indicated. To estimate efficiency of dystrophin protein rescue, the first 3 lanes were loaded with different percentages (40% - 20% - or 5% for TA and heart, and 10% - 5% - or 1% for diaphragm) of WT lysate. Coomassie blue staining of each membrane is shown below each blot, demonstrating equivalent protein loading. Dashed squares highlight the regions of each blot used for quantification of dystrophin protein (427 kDa).

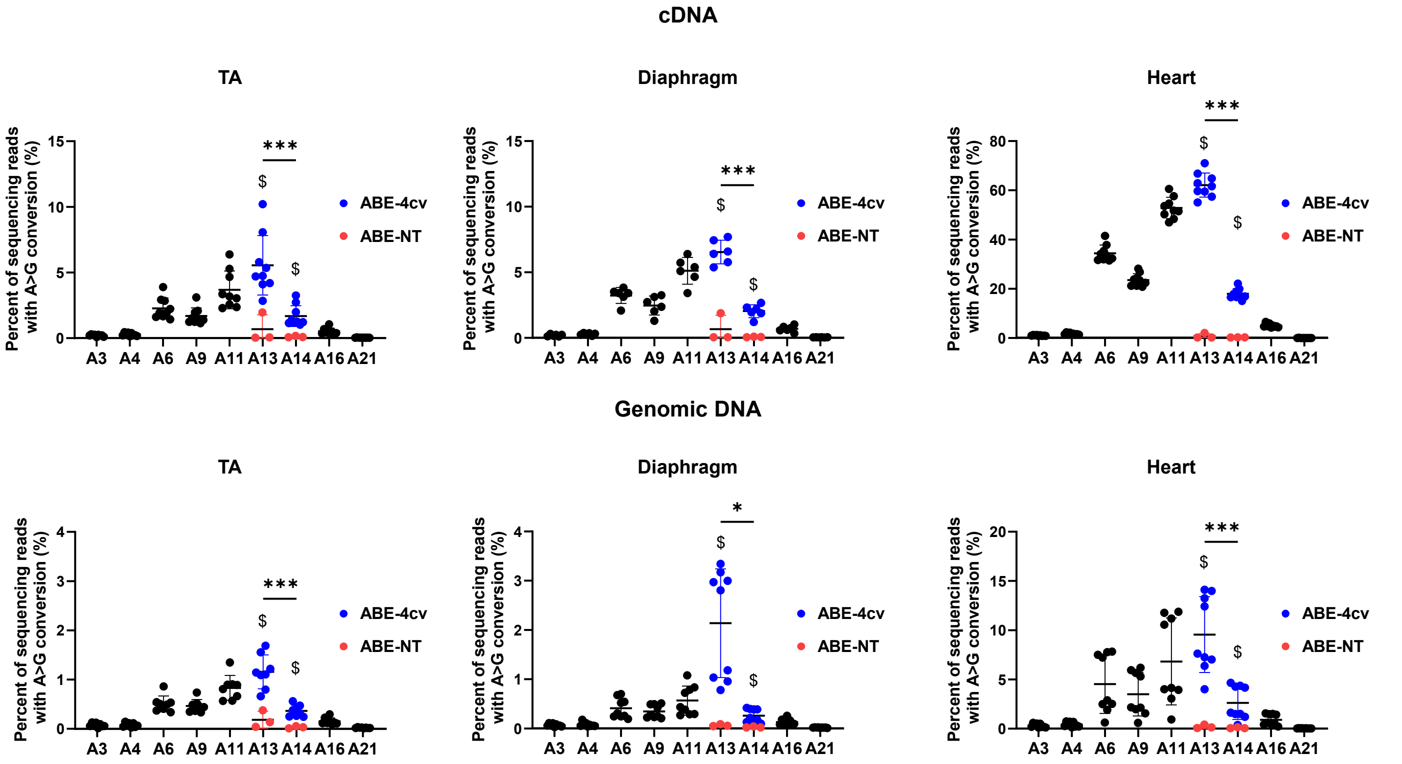


**Fig. S18. Amplicon sequencing read analysis of cDNA and genomic DNA from TA, diaphragm, and heart of male *mdx*$^{4cv}$ mice injected retro-orbitally with MyoAAV-SaABE8e$^{4cv}$ at P7.** Mice were retro-orbitally (RO) injected at P7 with 4E13 VG/kg MyoAAV-SaABE8e$^{4cv}$ and TA, diaphragm, and heart harvested one month later. cDNA and genomic DNA from TA, diaphragm, and heart were used for amplicon sequencing at the *mdx*$^{4cv}$ mutation site. A>G conversion rates for individual adenines (A) of MyoAAV-SaABE8e$^{4cv}$-injected *mdx*$^{4cv}$ mice were calculated from sequencing reads. Adenine conversions in the *mdx*$^{4cv}$ premature stop codon from MyoAAV-SaABE8e$^{4cv}$-injected mice are shown in dark blue (A13 and A14). A13 and A14 A>G conversion rates from MyoAAV-SaABE8e$^{NT}$-injected *mdx*$^{4cv}$ mice are shown in red. $ indicates a significant difference between MyoAAV-SaABE8e$^{4cv}$- and MyoAAV-SaABE8e$^{NT}$-injected *mdx*$^{4cv}$ mice in A13 or A14 A>G conversion rates (determined by unpaired T-test). n = 6-9 biological replicates/group; analyzed with one-way repeated measures ANOVA. Data are shown as mean ± SD. * and *** indicate $P < 0.05$ and <0.001, respectively.

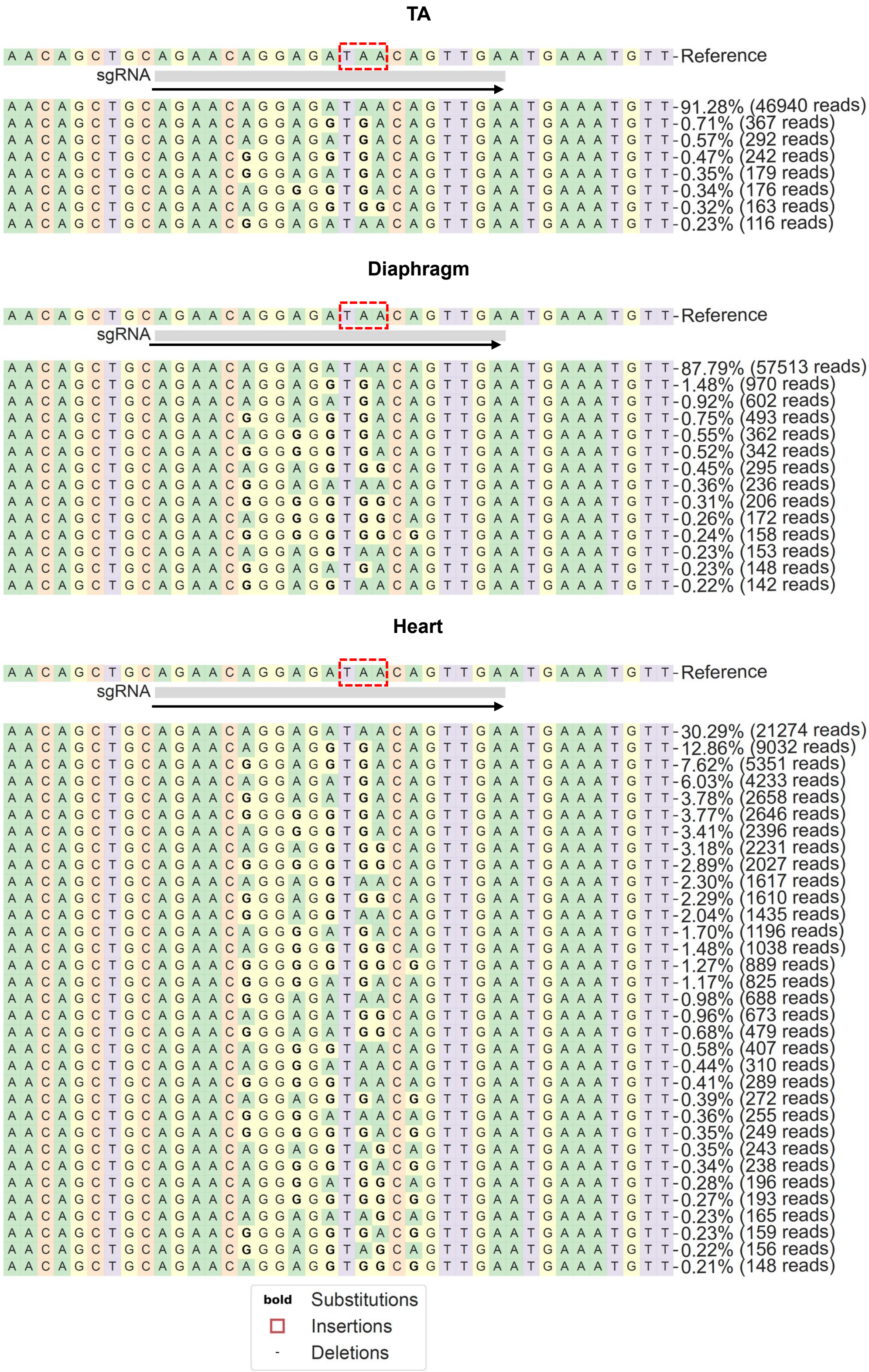


**Fig. S19. Amplicon sequencing read analysis at the site of the *mdx*$^{4cv}$ nonsense mutation for the indicated tissues from MyoAAV-SaABE8e$^{4cv}$ injected P7 *mdx*$^{4cv}$ mice.** Mice were retro-orbitally injected at P7 with 4E13 VG/kg MyoAAV-SaABE8e$^{4cv}$ and TA, diaphragm, and heart harvested one month later. cDNA amplicon sequencing results were analyzed with CRISPResso2 algorithm. Representative allele frequency tables for each sample type are shown. Dashed red box indicates the *mdx*$^{4cv}$ nonsense mutation site, and arrow indicates the direction of the gRNAs.

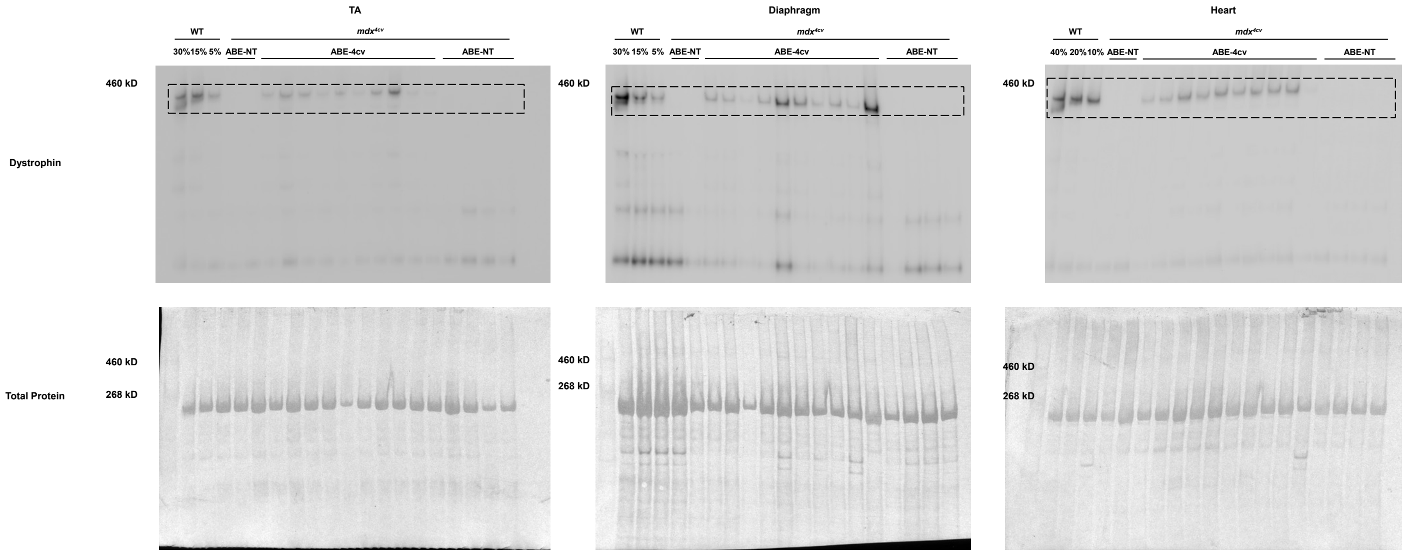


**Fig. S20. Diminished efficiency of dystrophin restoration with systemic administration of MyoAAV-SaABE8e$^{4cv}$ to young adult *mdx$^{4cv}$* mice.** Young adult (12W) male *mdx$^{4cv}$* mice were retro-orbitally injected with 4E13 VG/kg MyoAAV-SaABE8e$^{4cv}$ or MyoAAV-SaABE8e$^{NT}$, and tissues harvested one month later. Full membrane images of dystrophin protein expression detected by Western Blot of protein lysate from TA (left), diaphragm (middle), or heart (right) of the indicated wild-type (WT) mice, or *mdx$^{4cv}$* mice injected with MyoAAV-SaABE8e$^{NT}$ (NT), or MyoAAV-SaABE8e$^{4cv}$ (4cv-RNA1). To estimate efficiency of dystrophin protein rescue, the first 3 lanes were loaded with different percentages (30% - 15% - or 5% for TA and diaphragm, and 40% - 2% - or 10% for heart) of WT lysate. Coomassie blue staining of each membrane is shown below each blot, demonstrating equivalent protein loading. Dashed squares highlight the regions of each blot used for quantification of dystrophin protein (427 kDa).

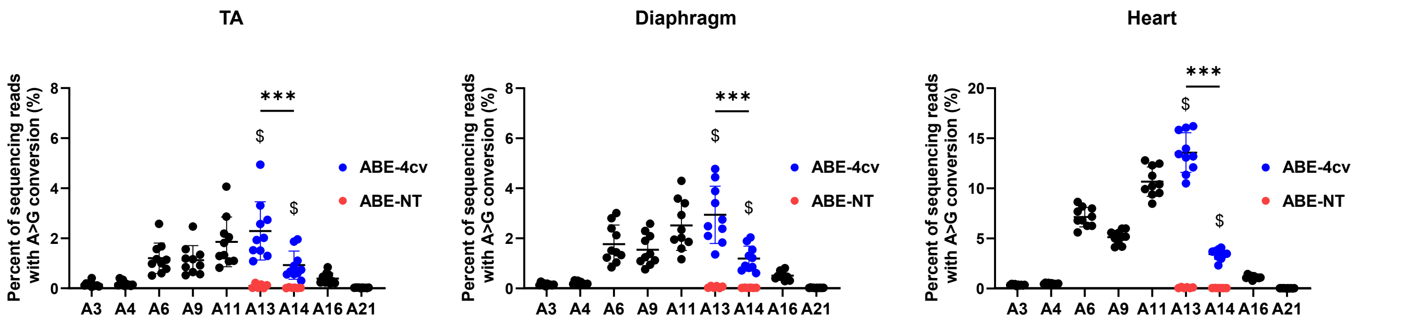


**Fig. S21. Amplicon sequencing read analysis of genomic DNA from TA, diaphragm, and heart of male *mdx*$^{4cv}$ mice injected retro-orbitally with MyoAAV-SaABE8e$^{4cv}$ at 12 weeks old.** Mice were retro-orbitally (RO) injected at 12W with 4E13 VG/kg MyoAAV-SaABE8e$^{4cv}$ or MyoAAV-SaABE8e$^{NT}$ and TA, diaphragm, and heart harvested one month later. Genomic DNA from TA, diaphragm, and heart were used for amplicon sequencing at the *mdx*$^{4cv}$ mutation site. A>G conversion rates for individual adenines (A) of MyoAAV-SaABE8e$^{4cv}$-injected *mdx*$^{4cv}$ mice were calculated from sequencing reads. Adenine conversions in the *mdx*$^{4cv}$ premature stop codon of MyoAAV-SaABE8e$^{4cv}$-injected *mdx*$^{4cv}$ mice are shown in dark blue (A13 and A14). A13 and A14 A>G conversion rates from control, MyoAAV-SaABE8e$^{NT}$-injected *mdx*$^{4cv}$ mice are shown in red. $ indicates a significant difference between MyoAAV-SaABE8e$^{4cv}$- and MyoAAV-SaABE8e$^{NT}$-injected *mdx*$^{4cv}$ mice in A13 or A14 A>G conversion rates (determined by unpaired T-test). n = 10 biological replicates/group; analyzed with one-way repeated measures ANOVA. Data are shown as mean ± SD. *** indicate $P < 0.001$.

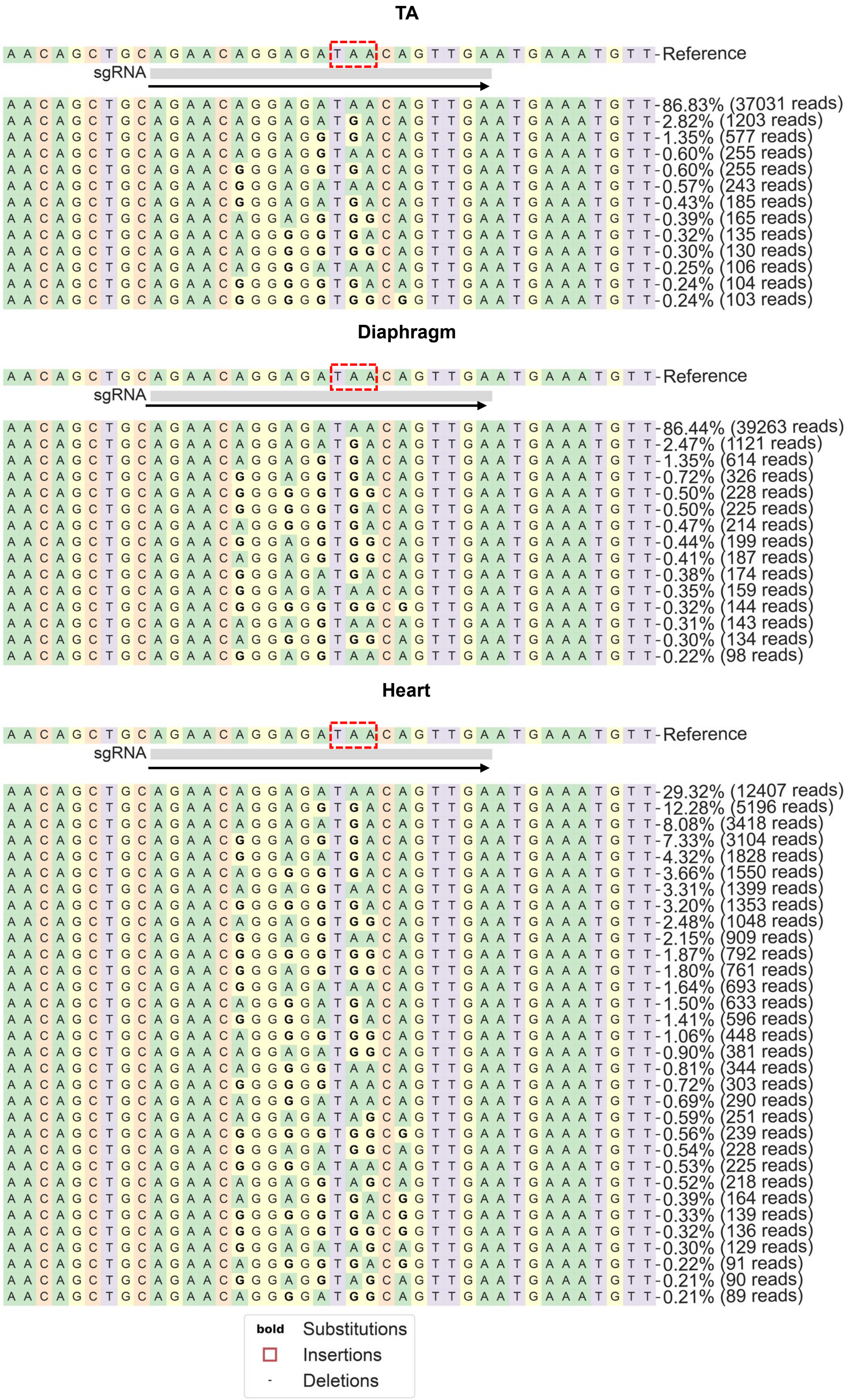


**Fig. S22. Amplicon sequencing read analysis at the site of the *mdx$^{4cv}$* nonsense mutation for the indicated tissues from *mdx$^{4cv}$* mice injected with MyoAAV-SaABE8e$^{4cv}$ at 12 weeks of age.** Mice were injected retro-orbitally at 12 weeks old with 4E13 VG/kg MyoAAV-SaABE8e$^{4cv}$ and TA, diaphragm, and heart were harvested one month later. Amplicon sequencing results from cDNA were analyzed with CRISPREsso2 algorithm. Representative allele frequency tables for each sample type are shown. Dashed red box indicates the *mdx$^{4cv}$* nonsense mutation site, and arrow indicates the direction of the gRNAs.

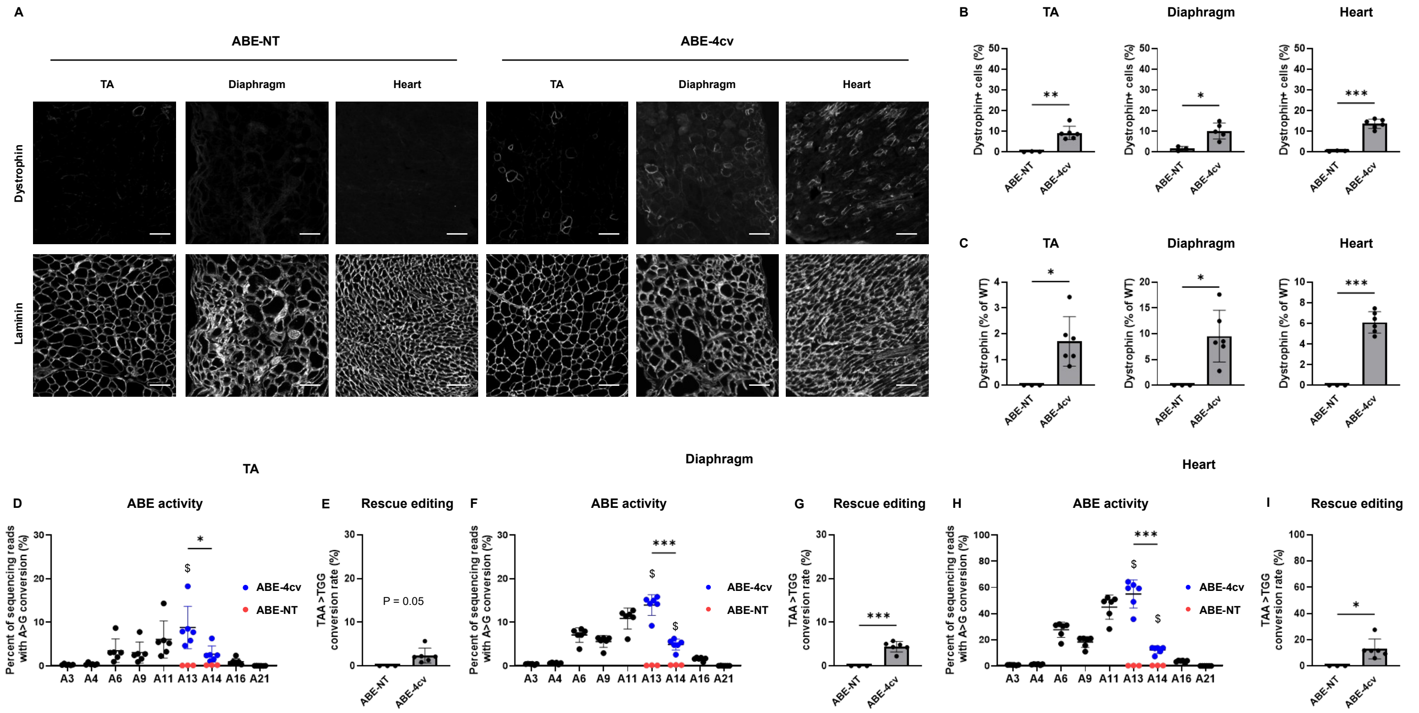


**Fig. S23. Systemic delivery of MyoAAV-SaABE8e$^{4cv}$ in 6-month-old *mdx*$^{4cv}$ mice yields inefficient rescue of dystrophin deficiency.** 6-month-old male *mdx*$^{4cv}$ mice were injected retro-orbitally with 4E13 VG/kg MyoAAV-SaABE8e$^{4cv}$ or MyoAAV-SaABE8e$^{NT}$, and tissues harvested one month later. (A) Representative single-channel immunofluorescence images from ABE-NT or ABE-4cv injected mice showing the same field of tibialis anterior (TA), diaphragm, or heart stained for dystrophin (top) or laminin (bottom), as indicated. Scale bar = 100 µm. (B) Quantification of dystrophin-positive myofibers or cardiomyocytes, calculated as percent (%) of total cells (derived from laminin staining). (C) Semi-quantitative analysis of dystrophin protein expression in TA, diaphragm, and heart of mice injected with MyoAAV-SaABE8e$^{NT}$ or MyoAAV-SaABE8e$^{4cv}$. Representative blots are shown in Fig. S24. (D-I) cDNA from TA (D-E), diaphragm (F-G), and heart (H-I) were analyzed by amplicon sequencing at the *mdx*$^{4cv}$ mutation site. Representative allele frequency tables are shown in Fig. S26. (D,F,H) A>G conversion rates for individual adenines (A) of MyoAAV-SaABE8e-injected *mdx*$^{4cv}$ mice were calculated from sequencing reads. Adenine conversions in the *mdx*$^{4cv}$ premature stop codon of MyoAAV-SaABE8e$^{4cv}$ -injected mice are shown in dark blue (A13 and A14). A13 and A14 A>G conversion rates from MyoAAV-SaABE8e$^{NT}$-injected *mdx*$^{4cv}$ mice are shown in red. $ indicates a significant difference between MyoAAV-SaABE8e$^{4cv}$- and MyoAAV-SaABE8e$^{NT}$-injected *mdx*$^{4cv}$ mice in A13 or A14 A>G conversion rates (determined by unpaired T-test). (E,G,I) Percentages (%) of reads with *mdx*$^{4cv}$ nonsense mutation converted to a non-stop (missense) mutation (TAA>TGG). n = 3-6 biological replicates/group; B, C, E, G, and I are analyzed with unpaired T-test, D, F, and H are analyzed with one-way repeated measures ANOVA. Data are shown as mean ± SD. *, **, and *** indicate $P < 0.05$, $< 0.01$, and $<0.001$, respectively.

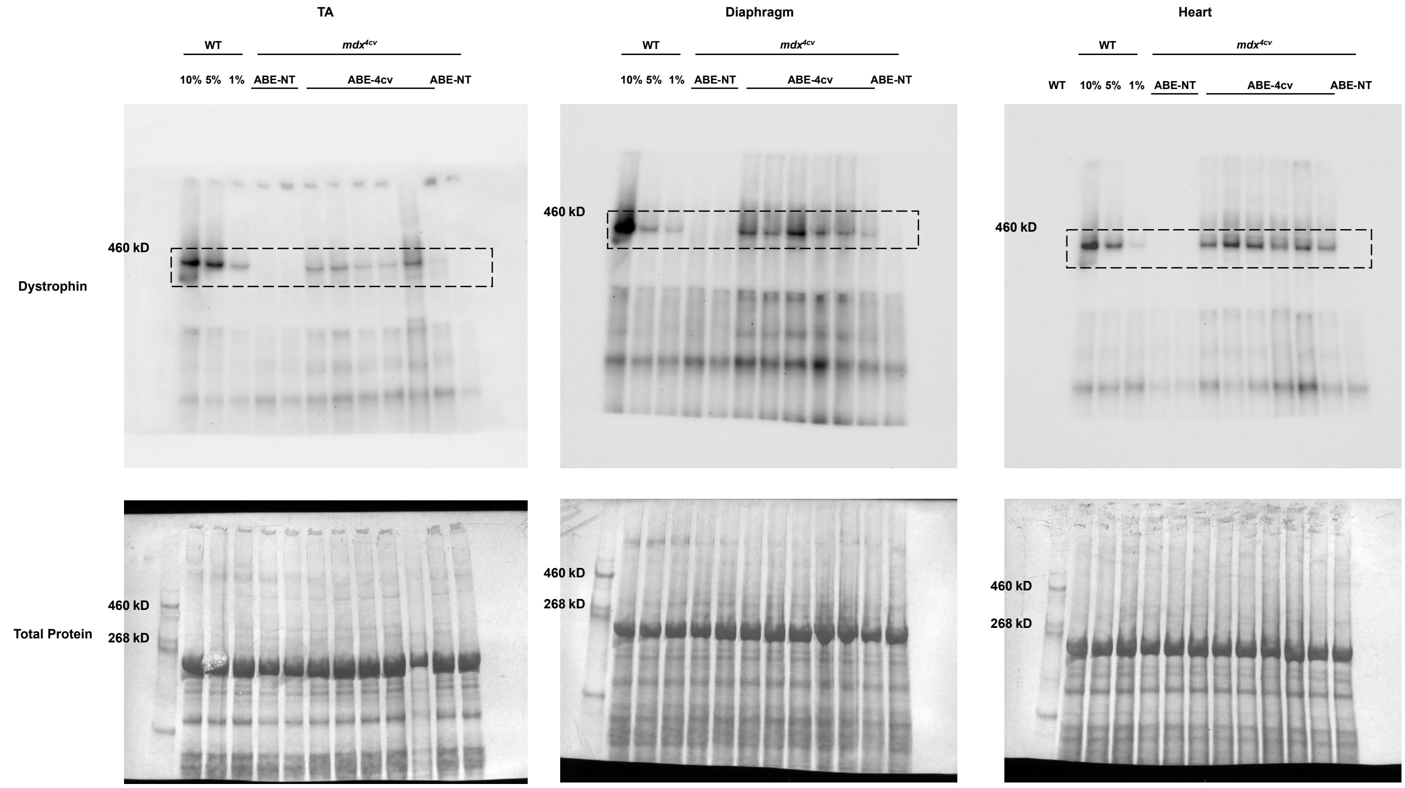


**Fig. S24. Systemic delivery of MyoAAV-SaABE8e$^{4cv}$ in 6-month-old *mdx$^{4cv}$* mice rescued dystrophin deficiency inefficiently.** Adult (6M) male *mdx$^{4cv}$* mice were injected retro-orbitally with 4E13 VG/kg MyoAAV-SaABE8e$^{4cv}$ or MyoAAV-SaABE8e$^{NT}$, and tissues harvested one month later. Full membrane images of dystrophin protein expression detected by Western Blot of protein lysate from TA (left), diaphragm (middle), or heart (right) of wild-type (WT) mice, or *mdx$^{4cv}$* mice injected with MyoAAV-SaABE8e$^{NT}$ (NT), or MyoAAV-SaABE8e$^{4cv}$ (4cv-RNA1), as indicated. To estimate efficiency of dystrophin protein rescue, the first 3 lanes were loaded with different percentages (10% - 5% - or 1%) of WT lysate. Coomassie blue staining of each membrane is shown below each blot, demonstrating equivalent protein loading. Dashed squares highlight the regions of each blot used for quantification of dystrophin protein (427 kDa).

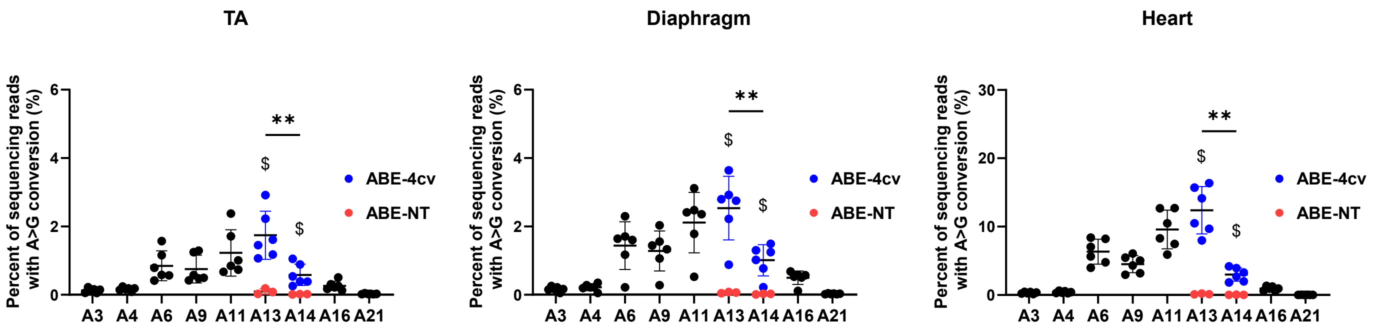


**Fig. S25. Amplicon sequencing read analysis of genomic DNA from TA, diaphragm, and heart of male *mdx*$^{4cv}$ mice injected retro-orbitally with MyoAAV-SaABE8e$^{4cv}$ at 6 months old.** Mice were retro-orbitally (RO) injected at 6 months old with 4E13 VG/kg MyoAAV-SaABE8e$^{4cv}$ and TA, diaphragm, and heart harvested one month later. Genomic DNA from TA, diaphragm, and heart were used for amplicon sequencing at the *mdx*$^{4cv}$ mutation site. A>G conversion rates for individual adenines (A) of MyoAAV-SaABE8e-injected *mdx*$^{4cv}$ mice were calculated from sequencing reads. Adenine conversions in the *mdx*$^{4cv}$ premature stop codon of MyoAAV-SaABE8e$^{4cv}$-injected mice are shown in dark blue (A13 and A14). A13 and A14 A>G conversion rates from MyoAAV-SaABE8e$^{NT}$-injected *mdx*$^{4cv}$ mice are shown in red. $ indicates a significant difference between MyoAAV-SaABE8e$^{4cv}$- and MyoAAV-SaABE8e$^{NT}$-injected *mdx*$^{4cv}$ mice in A13 or A14 A>G conversion rates (determined by unpaired T-test). n = 6 biological replicates/group; analyzed with one-way repeated measures ANOVA. Data are shown as mean ± SD. *** indicate P < 0.001.

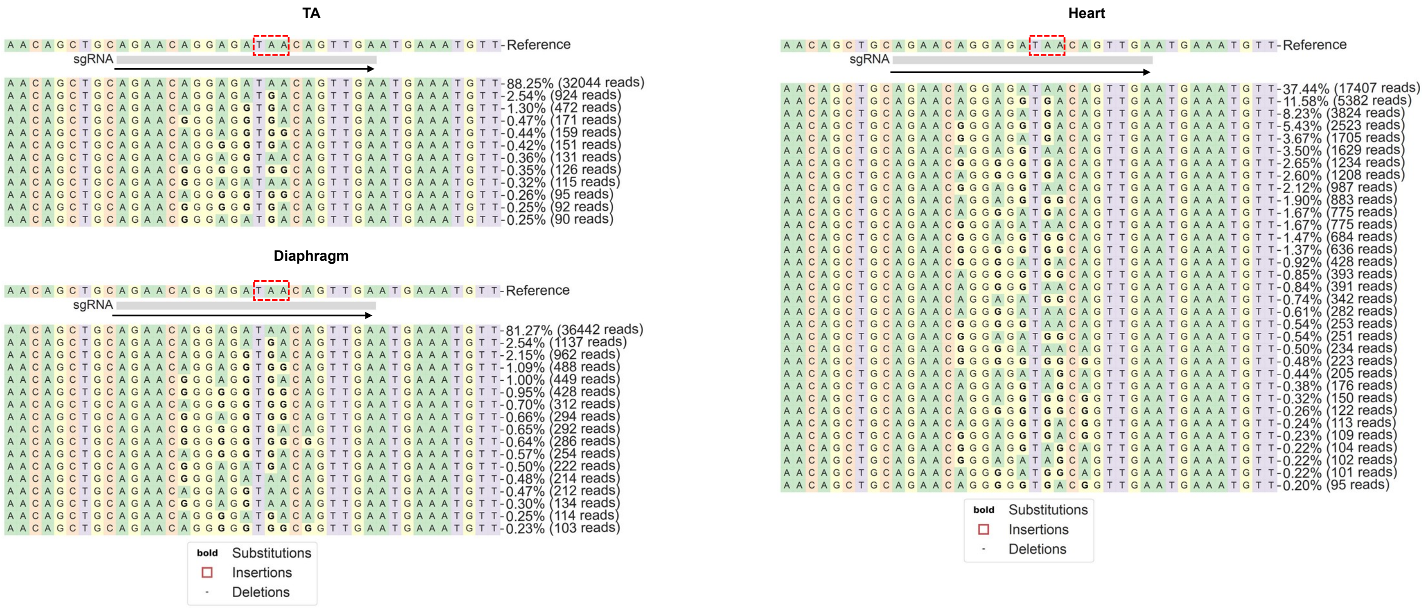


**Fig. S26. Amplicon sequencing read analysis at the site of the $mdx^{4cv}$ nonsense mutation for the indicated tissues $mdx^{4cv}$ mice injected with from MyoAAV-SaABE8e$^{4cv}$ at 6 months of age.** Mice were injected retro-orbitally at 6 months old with 4E13 VG/kg MyoAAV-SaABE8e$^{4cv}$ and TA, diaphragm, and heart were harvested one month later. cDNA from TA, diaphragm, and heart were used for amplicon sequencing. Amplicon sequencing results from cDNA were analyzed with CRISPREsso2 algorithm. Representative allele frequency tables for each sample type are shown. Dashed red box indicates the $mdx^{4cv}$ nonsense mutation site, and arrow indicates the direction of the gRNAs.

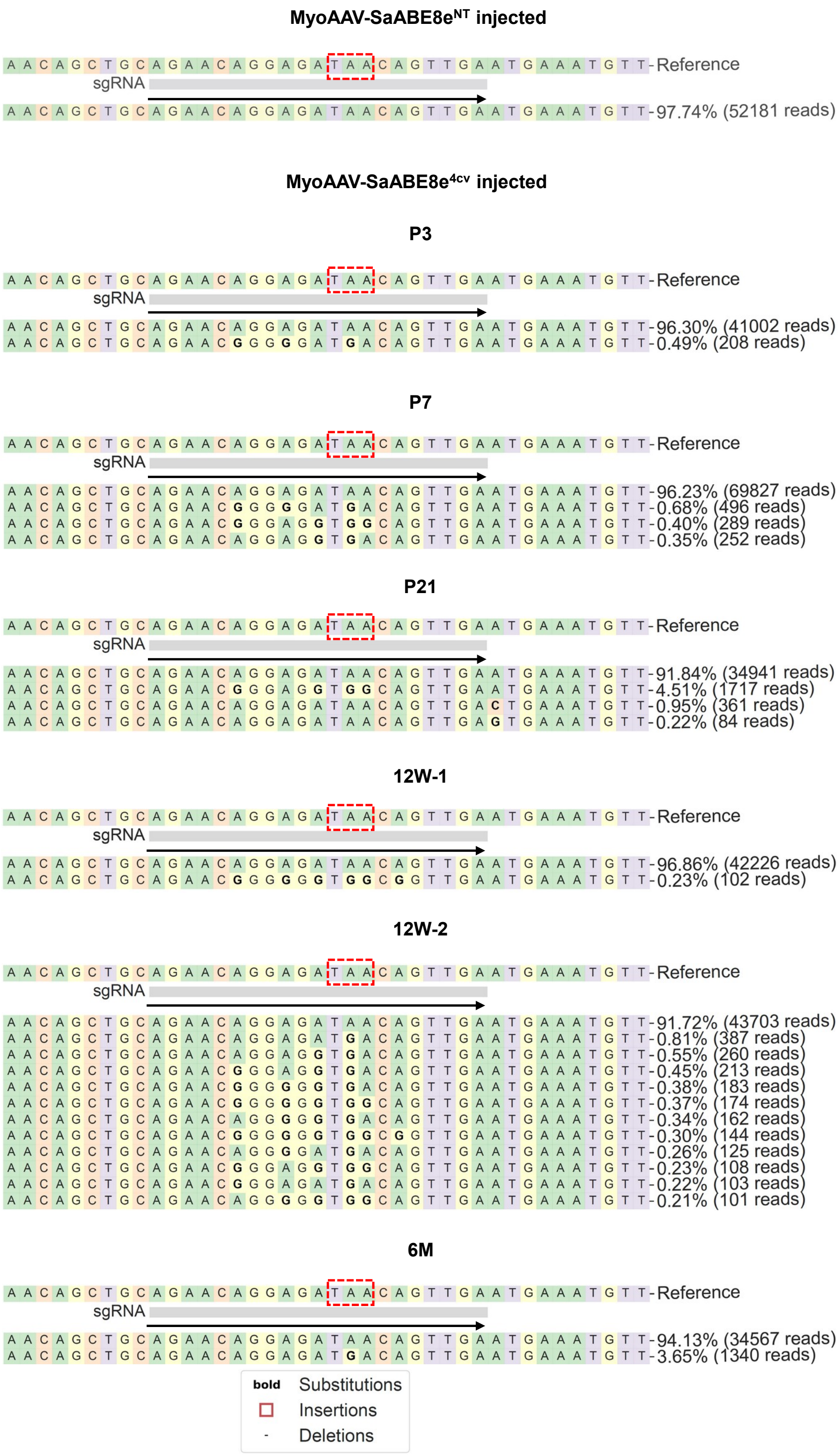


**Fig. S27. Amplicon sequencing read analysis at the site of the *mdx*$^{4cv}$ nonsense mutation in satellite cells from *mdx*$^{4cv}$ mice treated with MyoAAV-SaABE8e$^{4cv}$ at the indicated age.** Mice were injected retro-orbitally at the indicated age with 4E13 VG/kg MyoAAV-SaABE8e$^{4cv}$ or MyoAAV-SaABE8e$^{NT}$, and satellite cells were FACSorted one month later. Genomic DNA of these cells was used for amplicon sequencing, and the amplicon sequencing results were analyzed with CRISPREsso2 algorithm. Representative allele frequency tables for the indicated conditions are shown. Because of the heterogeneous outcomes in 12W MyoAAV-SaABE8e$^{4cv}$ injected mice, 2 allele frequency tables (12W-1 and 12W-2) are shown. Dashed red box indicates the *mdx*$^{4cv}$ nonsense mutation site, and arrow indicates the direction of the gRNAs.

1 Maesner, C. C., Almada, A. E. & Wagers, A. J. Established cell surface markers efficiently isolate highly overlapping populations of skeletal muscle satellite cells by fluorescence-activated cell sorting. *Skelet Muscle* **6**, 35 (2016). https://doi.org/10.1186/s13395-016-0106-6

**Key resources table**

**Oligo Sequences:**

| ID | Sequence (5’>3’) | Description |
|---|---|---|
| **Amplicon Sequencing Primers** | | |
| *mdx4cv* genomic DNA FWD | TCAGATTCAGTGGGATGAGGT | Genomic DNA amplicon sequencing |
| *mdx4cv* genomic DNA REV | ATGGCATGCTTTCATTTGCTAT | Genomic DNA amplicon sequencing |
| *mdx4cv* cDNA FWD | GATTTGGAACAGAGACGCCCC | cDNA amplicon sequencing |
| *mdx4cv* cDNA REV | TGCCTCTGACCTGTCCTATG | cDNA amplicon sequencing |
| **Droplet Digital PCR** | | |
| bGH FWD | GCCAGCCATCTGTTGT | AAV viral genome quantification |
| bGH REV | GGAGTGGCACCTTCCA | AAV viral genome quantification |
| bGH-probe | /56-FAM/TCCCCCGTG/ZEN/CCTTCCTTGACC/3IABkFQ/ | AAV viral genome quantification |
| Gapdh-F | CGCCCTGATCTGAGGTTAAAT | Host genome quantification |
| Gapdh-R | CGGAGCAACAGATGTGTGTA | Host genome quantification |
| Gapdh-probe | /5HEX/AGCCGTGTG/ZEN/ACCTTTCTGGATCTG/3IABkFQ/ | Host genome quantification |

**Plasmids:**

| ID | Source | Description |
|---|---|---|
| AAV-EFS-SaABE8eV106W-bGH-U6-sgRNA-BsmBI (Addgene_189924) | Addgene | Parental plasmid for SaABE8e$^{4cv}$ and SaABE8e$^{NT}$ |

| | | |
|---|---|---|
| EGFP reporter plsamid | Addgene | |
| SaABE8e$^{4cv\text{-}gRNA1}$ | Constructed in this study | 4cv-gRNA1 was cloned into Addgene_189924 |
| SaABE8e$^{4cv\text{-}gRNA2}$ | Constructed in this study | 4cv-gRNA2 was cloned into Addgene_189924 |
| SaABE8e$^{NT}$ | Constructed in this study | A non-targeting gRNA was cloned into Addgene_189924 |
| MyoAAV2A Rep/Cap plasmid | Gift from Dr. Jeff Molkentin's laboratory | MyoAAV production |
| pAdDeltaF6 | Puresyn | MyoAAV production |

**Antibodies and Dyes:**

| **ID** | **Source** | **Catalog Number** | **Dilution Factor** |
|---|---|---|---|
| **Satellite Cell Isolation** | | | |
| APC-Cy7-CD45 Ab | Biolegend | 103116 | 1:200 |
| APC-Cy7-CD11b Ab | Biolegend | 101226 | 1:200 |
| APC-Cy7-TER119 Ab | Biolegend | 116223 | 1:200 |
| APC-Sca-1 Ab | Biolegend | 108112 | 1:200 |
| PE-CD29 | Biolegend | 102208 | 1:100 |
| Biotin-CXCR4 | BD Biosciences | BDB551968 | 1:100 |
| Streptavidin-PE-Cy7 | Biolegend | 405206 | 1:200 |
| Propidium Iodide | Biolegend | 421301 | 1:200 |
| Calcein Blue, AM | ThermoFisher | C1429 | 1:200 |
| **Immunofluorescence** | | | |
| anti-Dystrophin Ab (MANDYS8) | Sigma Aldrich | D8168 | 1:100 |
| anti-Laminin Ab (LAMA1) | Sigma Aldrich | L9393 | 1:200 |

| | | | |
|---|---|---|---|
| α-sarcoglycan Ab [Ad1/20A6] | GeneTex | GTX01939 | 1:50 |
| β-sarcoglycan Ab [B-SARC-L-A] | Leica | NCL-L-b-SARC | 1:100 |
| β-dystroglycan Ab [B-DG-A] | Leica | NCL-b-DG | 1:100 |
| n-NOS:N-Terminal Ab | Immunostar | 24431 | 1:1000 |
| goat anti-mouse IgG2b Alexa Fluor 594 | ThermoFisher | A-21145 | 1:1000 |
| goat anti-rabbit IgG Alexa Fluor 488 | Invitrogen | A11008 | 1:250 |
| goat anti-mouse IgG1 Alexa Fluor 488 | ThermoFisher | A-21121 | 1:250 |
| goat anti-rabbit IgG Alexa Fluor 647 | ThermoFisher | A-21245 | 1:250 |
| goat anti-mouse IgG2a Alexa Fluor 647 | ThermoFisher | A-21241 | 1:250 |
| Hoechst 33342 | BD Biosciences | 561908 | 1:1000 |
| **Western Blot** | | | |
| anti-Dystrophin Ab [EPR9598(ABC)] | abcam | ab154168 | 1:2000 |
| Coomassie Brilliant Blue G-250 | ThermoScientific | 20279 | |

**Animal, Cell line, and Bacteria:**

| **ID** | **Source** | **Description** |
|---|---|---|
| B6Ros.Cg-*Dmd*$^{mdx\text{-}4Cv}$/J | Jackson Laboratory, #002378 | Mouse DMD disease model |
| C57BL/6J | Jackson Laboratory, #000664 | Mouse wild type |
| HEK293 | ATCC | AAV production |

| | | |
|---|---|---|
| NEB® Stable Competent *E. coli* | NEB | Inverted Terminal Repeat sequence containing plasmids |

**Reagents:**

| **Name** | **Source** | **Catalog Number** |
|---|---|---|
| Puromycin Dichloride | ThermoFisher | A1113802 |
| HindIII-HF® | NEB | R3104T |
| NdeI | NEB | R0111S |
| Dulbecco's modified eagle's medium (DMEM) | ThermoFisher | 11965-118 |
| Fetal bovine serum (FBS) | GeminiBio | 26140079 |
| Penicillin-Streptomycin (Pen-Strep) | ThermoFisher | 15140148 |
| Corning™ HYPERFlask™ M Cell Culture Vessels | VWR | 10031 |
| POROS GoPure AAVX Pre-packed Column | ThermoFisher | A36651 |
| ddPCR Supermix for Probes (No dUTP) | Bio-Rad | 186-3024 |
| Droplet Generation Oil for Probes | Bio-Rad | 1863005 |
| Collagenase type II | ThermoFisher | 17101015 |
| Ham's F-10 Nutrient Mix | ThermoFisher | 11550043 |
| Dispase | ThermoFisher | 17105041 |
| Hank's Balanced Salt Solution (HBSS) | ThermoFisher | 14025134 |
| Horse serum | ThermoFisher | 16050122 |
| Glutamax | ThermoFisher | 35050061 |
| Collagen type I | ThermoFisher | A1048301 |

| | | |
|---|---|---|
| Laminin | ThermoFisher | 23017015 |
| Fibroblast Growth Factor-Basic, human | Sigma Aldrich | F0291-25ug |
| Lipofectamine™ Stem | ThermoFisher | STEM00001 |
| DNeasy Blood & Tissue Kit | Qiagen | 69506 |
| TRIzol™ Reagent | ThermoFisher | 15596026 |
| RNeasy MiniPrep Kit | Qiagen | 74106 |
| ezDNase™ Enzyme | ThermoFisher | 11766051 |
| SuperScript™ IV First-Strand Synthesis System | ThermoFisher | 18091050 |
| Q5® Hot Start High-Fidelity 2X Master Mix | NEB | M0494L |
| QIAquick PCR Purification Kit | Qiagen | 28106 |
| Optimal Cutting Temperature Compound | Tissue-Tek | 4583 |
| Triton™ X-100 | Sigma Aldrich | T8787 |
| Normal Goat Serum | Jackson ImmunoResearch | 005-000-121 |
| Bovine Serum Albumin | Sigma Aldrich | A4737 |
| TWEEN® 20 | Sigma Aldrich | P2287 |
| M.O.M. (Mouse On Mouse) Immunodetection Kit, Basic | VectorLabs | BMK-2202 |
| Qubit Protein Broad Range Assay | ThermoFisher | A50669 |
| Pierce RIPA Lysis and Extraction Buffer | ThermoFisher | 89901 |

| | | |
|---|---|---|
| Halt™ Protease and Phosphatase Inhibitor Cocktail (100X) | ThermoScientific | 78429 |
| NuPAGE™ Tris-Acetate Mini Protein Gels, 3 to 8%, 1.0 mm, WedgeWell™ format | ThermoFisher | TA03815BOX |
| Trans-Blot Turbo Mini 0.2 µm PVDF Transfer Packs | BioRad | 1704156 |
| SuperSignal™ West Femto Maximum Sensitivity Substrate | ThermoFisher | 34096 |
